# POLCAM: Instant molecular orientation microscopy for the life sciences

**DOI:** 10.1101/2023.02.07.527479

**Authors:** Ezra Bruggeman, Oumeng Zhang, Lisa-Maria Needham, Markus Körbel, Sam Daly, Matthew Cheetham, Ruby Peters, Tingting Wu, Andrey S. Klymchenko, Simon J. Davis, Ewa K. Paluch, David Klenerman, Matthew D. Lew, Kevin O’Holleran, Steven F. Lee

## Abstract

Current methods for single-molecule orientation localization microscopy (SMOLM) require optical setups and algorithms that can be prohibitively slow and complex, limiting the widespread adoption for biological applications. We present POLCAM, a simplified SMOLM method based on polarized detection using a polarization camera, that can be easily implemented on any wide-field fluorescence microscope. To make polarization cameras compatible with single-molecule detection, we developed theory to minimize field of view errors, used simulations to optimize experimental design, and developed a fast algorithm based on Stokes parameter estimation which can operate over 1000 fold faster than the state of the art, enabling near instant determination of molecular anisotropy. To aid in the adoption of POLCAM, we developed open-source image analysis software, and a website detailing hardware installation and software use. To illustrate the potential of POLCAM in the life sciences, we applied our method to study alpha-synuclein fibrils, the actin cytoskeleton of mammalian cells, fibroblast-like cells and the plasma membrane of live human T cells.

## Introduction

Single-molecule localization microscopy (SMLM) [1–4] is a super-resolution microscopy technique that is widely used in biology to study cellular structures below the diffraction limit [5–7]. Single-molecule orientation localization microscopy (SMOLM) is a multidimensional variant of SMLM in which in addition to the precise spatial position, the orientation of individual fluorescent molecules is also measured. The ability to measure the orientation of single molecules provides information about how molecules organize, orient, rotate, and wobble in their environment, which is of key relevance across biological systems [8–13]. The widespread use of SMOLM by the biological imaging community has thus far been limited by the need for complex experimental setups and often computationally expensive image analysis. Additionally, this lack of accessibility has also slowed down the necessary development of a wider range of labeling protocols that are appropriate for SMOLM: labeling methods in which the orientation of the fluorescent probe is relatively fixed and rotationally restricted with respect to its target [14–17].

Fluorescent molecules are not isotropic point sources, *i*.*e*. they do not emit light equally in all directions. Fundamentally, fluorescent molecules emit like oscillating electric dipoles: the intensity *I* of the emitted fluorescence depends on the relative observation direction and follows the relationship *I ∝*sin^2^(*η*), where *η* is the angle between the observation direction and the orientation of the emission dipole moment of the molecule [18]. In conventional SMLM experiments, this anisotropic emission is typically not noticeable because, with common SMLM labeling protocols, fluorescent molecules are free to rapidly wobble and rotate with respect to their target (*e*.*g*., due to long linker chains), resulting in an orientation-averaged image [19]. However, when a labeling method is used that restricts the rotational freedom of the fluorescent molecules with respect to their target, the anisotropy in the emitted fluorescence allows for the measurement of molecular orientation [20]. Different approaches have been used to achieve molecular orientation imaging. Some methods are based on active modulation of the polarization of the excitation light [21–23], but the majority of methods are based on modification of the detection path of the microscope. The image of a single molecule can be fitted using a dipole-spread function (DSF) that includes the position of the molecule (*x, y* or *x, y, z*), the orientation of the emission dipole moment (*ϕ, θ*) [24, 25] and often a rotational mobility parameter [26].

As the intensity distribution of a standard DSF does not contain significant information about a molecule’s orientation, the DSF can be engineered to increase the orientation information content. A simple example is imaging slightly out-of-focus to exaggerate the DSF shape [27–29]. More advanced DSF engineering can be performed using a spatial light modulator [19, 30–33], a special optic [34–36] or pupil splitting [37]. A drawback of DSF engineering is that the optical setups required are highly complex (with the exception of the Vortex DSF [34]), and are sensitive to optical aberrations (with the exception of pupil splitting [37]) as the orientation estimation algorithms rely on simulated DSF models, which can necessitate performing spatially (in)variant phase-retrieval to match the DSF model to the experimental DSF [34]. Additionally, fitting a 5- or 6-dimensional DSF model is computationally expensive, making data analysis prohibitively slow.

An alternative method is splitting the emission into multiple polarized channels that form separate images on the same camera [10, 38–44], multiple detectors [45], or using continuous image displacement using a rotating calcite crystal [46]. The advantage of polarized detection-based methods is that the orientation estimation can be performed using simple intensity measurements, and does not necessarily require the fitting of a complex DSF model. As a result, the data analysis is fast and easily compatible with high-throughput data collection. The simplest polarized detection setup splits the emission into 2 orthogonal polarized channels using a polarizing beam splitter [38, 39]. This method suffers from some degeneracies since both an isotropic emitter (*e*.*g*., a freely rotating molecule/fluorescent bead), an immobilized molecule with an emission dipole moment oriented at 45° in between the transmission axis of the two channels, or parallel to the optical axis all give rise to equal intensities measured in both channels [8, 47]. Splitting the emission into 3 or more polarized channels/cameras breaks this degeneracy [15, 48, 49], but significantly complicates the experimental setup [39, 41–44].

Here, we present a new experimentally simplified SMOLM method called POLCAM that uses a polarization camera for four-channel polarized detection. Polarization cameras have become popular in the field of computer vision as they provide single-shot multi-channel polarized measurements [50–53]. The pixels of the sensor of a polarization camera are covered by small linear polarizers with transmission axes typically oriented at 0°, 45°, 90° and −45° in a 2 × 2-pixel mosaic pattern that is repeated over the entire sensor (Fig. 1a). As polarizers are integrated into the camera chip, no additional polarization optics are required.

**Figure 1:**
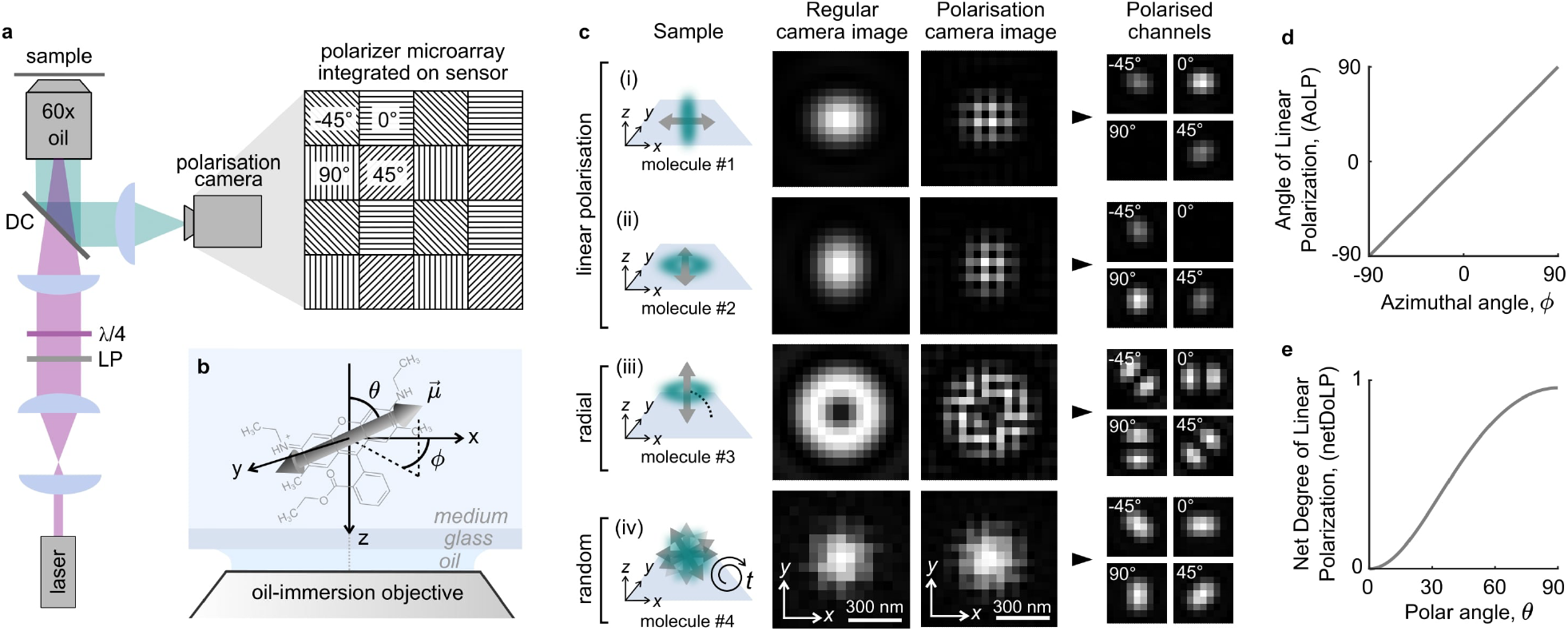
Single-molecule imaging using a polarization camera. **a)** Schematic of the optical setup that includes the polarization camera and a schematic representation of a small region of the four-directional micropolarizer array (transmission axis at 0°, 45°, 90°, or −45°) integrated into the sensor. LP linear polarizer, *>*/4 quarter-wave plate, DC dichroic. **b)** Definition of the in-plane angle *ϕ* and out-of-plane angle *θ* that specify the orientation of the emission dipole moment (arrow) of a molecule (structure of Rhodamine 6G depicted). **c)** Simulated examples of four single fluorescent molecules. From top to bottom, a molecule (i) aligned with the x-axis, (ii) the y-axis, (iii) the optical axis (z-axis), and (iv) a rapidly rotating molecule. For each example, the following is shown: the image plane recorded with a regular monochrome camera, the image plane recorded with a polarisation camera in its raw format, and in a format where the pixels have been rearranged to form four images that are each made up only of pixels that are covered by a micropolariser with the same transmission axis orientation. **d)** Relationship between the average angle of linear polarization (AoLP) and the in-plane angle *ϕ* of the dipole moment. **e)** Relationship between the net degree of linear polarization (netDoLP) and the out-of-plane angle *θ* of the dipole moment of a rotationally immobilized molecule for a 1.4 NA oil-immersion objective and no refractive index mismatch between the sample and immersion medium.

Polarization cameras have reached quantum efficiencies and noise levels that in theory are compatible with single-molecule detection, but the ease of use of polarization cameras does come at a cost in the form of *instantaneous field-of-view* (IFOV) errors near object edges [54–56]. To make POLCAM robust to IFOV errors, we used vectorial diffraction simulations of single dipole emitters to optimize microscope design and developed a Stokes parameter estimation-based reconstruction algorithm, and a DSF-fitting algorithm.

We validated and characterized our method using samples with known structures; single fluorophores immobilized on cover glass, in polymer, and lipid bilayer-coated glass beads labeled with membrane dyes. We next performed SMOLM on amyloid fibrils *in vitro* and on the actin network of fixed mammalian cells. To demonstrate that a polarization camera can also be used for conventional polarized detection microscopy, we imaged the actin network of COS-7 cells, and the membrane of live human T cells interacting with an antibody-coated cover glass.

Our approach can be easily implemented by switching the regular camera on any single-molecule fluorescence microscope for a polarization camera. The camera used in this work is supported by the popular image acquisition software Micro-Manager [57], and we provide image analysis software in the form of MATLAB applications for single-molecule and diffraction-limited image analysis and real-time image processing and rendering during acquisition, and a napari [58] plugin for processing multi-dimensional diffraction-limited polarization camera image datasets. We envisage that the combination of ease of use, ease of implementation, low cost, improved speed, and open-source software will make POLCAM an accessible and powerful tool for the study of molecular orientation across diverse biological applications.

## Results

### Measuring molecular orientation using polarized detection

When a fluorescent molecule has one dominant emission dipole moment *μ* —as is the case for many common fluorescent molecules— its emission resembles the far-field emitted by an oscillating electric dipole [18, 20] (Fig. 1b-c, Supplementary Note S1 and S3). When its emission dipole moment is oriented parallel to the sample plane, the electric field in the back focal plane of the objective will mainly be *linearly polarized* along the direction of the emission dipole moment (Fig. 1c, first and second row). If the emission dipole moment is oriented parallel to the optical axis, the electric field in the back focal plane of the objective will be *radially polarized* [59, 60] (Fig. 1c, third row). When a molecule is rapidly rotating, it can appear unpolarized (Fig. 1c, bottom row). As the tube lens in conventional widefield fluorescence microscopy has a low numerical aperture, the described polarization is mostly conserved in the image plane [61]. As a result, the angle of the axis of maximum polarization determines the in-plane orientation *ϕ* of the emission dipole moment, and the degree of net linear polarization is related to the out-of-plane orientation *θ* (Fig. 1d-e).

Conventionally, polarizing beam splitters are used to split the detected fluorescence into multiple polarized image channels. The number of photons that are detected from a single molecule in the different channels will depend on the 3D orientation and rotational mobility of the molecule. These measured intensities can be used to estimate the angles *ϕ* and *θ* using analytically derived equations. Equations for the case of four polarized detection channels (0°, 45°, 90°, and −45°) were derived by John T. Fourkas [60]. Here, we rewrote these expressions in terms of Stokes parameters (Supplementary Notes S4 and S5):

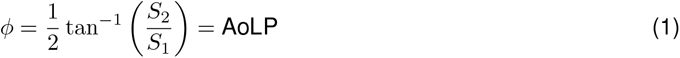

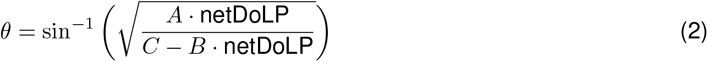

where *A, B* and *C* are constants that are a function of the half maximum collection angle of the objective *α* (Supplementary Notes S4 and S5) and the net degree of linear polarization (netDoLP) given by

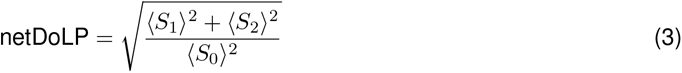

where the brackets ⟨… ⟩ refer to averaging over a small region of interest around a single molecule (Supplementary Note S7), and *S*_0_, *S*_1_ and *S*_2_ are the first three Stokes parameters:

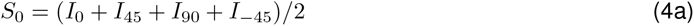

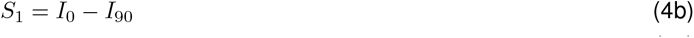

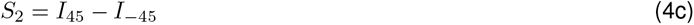

where *I*_0_, *I*_45_, *I*_90_, and *I*_*-*45_ refer to the measured intensities in the four polarized channels. The full derivation of Eq. (46) and (47) can be found in Supplementary Notes S4 and S5, and example simulated images in Supplementary Figures S3–S19.

We note that Eq. (46) is simply the expression for the angle of linear polarization (AoLP) [62] and that Eq. (47) depends only on the net degree of linear polarization and the objective used. This is in line with our intuition (Fig. 1c). Eq. (46) and Eq. (47) are plotted in figure 1d,e for a 1.4 NA oil-immersion objective. We note that estimation of *ϕ* using Eq. (46) is very robust, but that estimation of *θ* using Eq. (47) on the other hand is only possible under strict conditions (perfect rotational immobilization, high signal-to-noise ratio, and no large refractive-index mismatch) and is more reliably performed with DSF-fitting as will be discussed in more detail in a later section.

As a proxy for rotational mobility, we use the average degree of linear polarization (avgDoLP) which we define as a local average of the degree of linear polarization (DoLP):

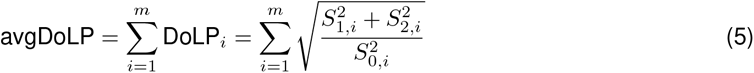

where *m* is the number of pixels in the region of interest around the molecule. If a DSF-fitting algorithm is used instead (see the section on considering the DSF shape), a rotational mobility parameter *γ* can be estimated that inversely relates to the size of a cone in which the molecule has rotational freedom [26]: *γ* is 1 for perfect immobilization, and 0 for complete rotational freedom. The exact mathematical relation between avgDoLP and *γ* is complex, as avgDoLP is also influenced by the signal-to-noise ratio, the out-of-plane angle, and the refractive index of the sample medium. The relationship between avgDoLP and rotational mobility is numerically explored in Fig. S27, showing that it is monotonic under all conditions and can therefore be used qualitatively, but with care. For details, we refer to the Methods and Supplementary Notes S7.

### Overcoming IFOV errors

In conventional polarization-sensitive fluorescence imaging, where polarizing elements are placed in the optical path, a traditional camera sensor captures the full intensity distribution everywhere in the image plane. However, similar to the operation of conventional color image sensors [63], the polarization camera measures the intensity of each polarization channel in a subset of the image (1 in 4 pixels), and the full intensity distributions have to be recovered through interpolation [54–56]. This recovery can be performed accurately provided that the pixel size is small enough. If this is not the case, any measurements taken from the recovered channels will exhibit what is known as *instantaneous field-of-view* (IFOV) errors [54–56]. In standard applications of polarization cameras (*e*.*g*., quality control in the manufacturing industry, removal of reflections in images [64]), IFOV errors can be ignored or avoided, as the pixel size can be much smaller than variations in neighboring pixels, and artefacts mostly appear near the edges of objects. However, when imaging single molecules, the opposite is true as the image of a single emitter varies significantly over each pixel, and the limited photon budget prevents the use of a small pixel size.

We determined the optimal pixel size for the estimation of molecular orientation using vectorial diffraction simulations (Supplementary Note S6). We define the optimal pixel size as the largest pixel size that still allows for accurate recovery of the four polarized channels from a single polarization camera image. To assess whether accurate recovery is possible, we used an approach described by Tyo *et al*. [55] that checks for overlap between the contributions of different Stokes parameters in the Fourier transform of the unprocessed polarization camera image (Supplementary Note S6.1, Supplementary Fig. S20–S25). If the contributions don’t overlap, the recovery is assumed to be accurate.

Using this method, we find that for our setup (1.4 NA oil-immersion objective, 650 nm wavelength, sample in an aqueous medium) optimal sampling is achieved at a pixel size of *∼* 60 nm *×* 60 nm (Supplementary Fig. S20). Practically, a pixel size of 57.5 nm *×* 57.5 nm was achieved using a 60*×* magnification objective and the standard polarization camera pixel size of 3.45 μm *×* 3.45 μm (Supplementary Fig. S30). The calculated ideal pixel size as a function of wavelength and objective numerical aperture and sample medium can be found in Supplementary figure S20.

Next, we compared the performance of different algorithms [54–56] for Stokes parameter estimation and channel interpolation on simulated polarization camera images of immobilized single molecules. We found that a Fourier-based approach [55] and cubic spline interpolation performed the best (Supplementary Note S6.2, Supplementary Fig. S21–S25). Figure 2a shows an experimental polarization camera image of semi-immobilized SYTOX orange molecules on a cover glass. Figure 2b shows the results of the Fourier-based interpolation. Figure 2c shows examples of the emission of single molecules taken from this dataset; two in-plane oriented molecules (molecule 1 and 2), one molecule oriented out of the plane (molecule 3) and a rapidly rotating molecule (molecule 4). An example SYTOX Orange dataset is included in the supporting information (Dataset 1).

**Figure 2:**
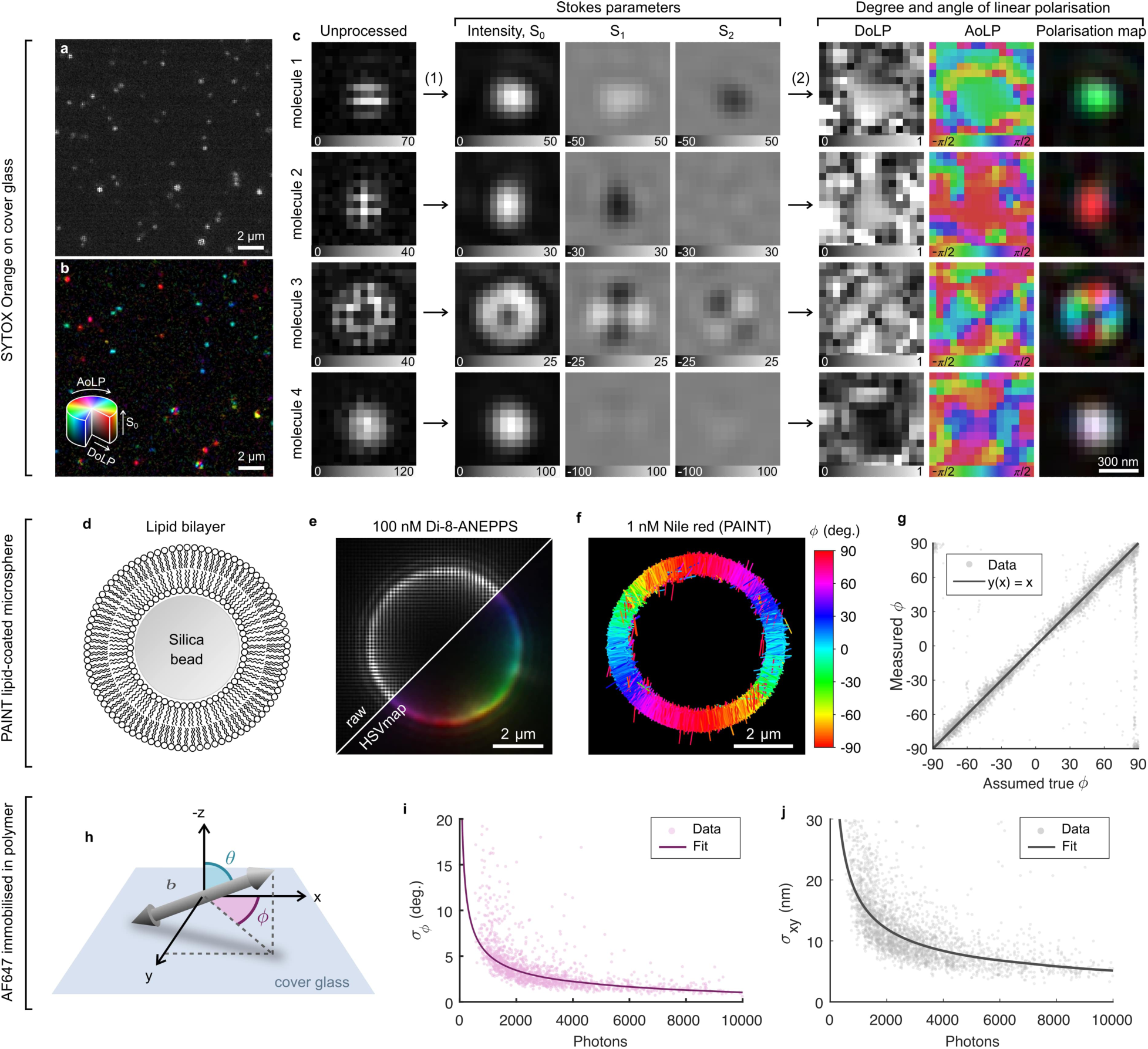
Single-molecule detection, experimental bias and precision. **a)** An unprocessed polarization camera image of SYTOX Orange molecules dispersed on a cover glass in PBS. **b)** The same image as in panel a but processed to reveal polarization information using a polarization colormap that combines the angle of linear polarization (AoLP), degree of linear polarization (DoLP), and intensity (S_0_) in HSV colorspace (hue = AoLP, saturation = DoLP, value = S_0_). **c)** Examples of 4 SYTOX Orange molecules with their emission dipole moment parallel to the sample plane (molecules 1 and 2), parallel to the optical axis (molecule 3), and rapidly rotating (molecule 4). For each, the unprocessed image, estimated Stokes parameter images (S_0_, S_1_ and S_2_), AoLP, DoLP, and polarization colormap images are shown. **d)** Illustration of a silica microsphere (5 μm diameter) coated using a lipid bilayer (DPPC + 40% cholesterol). **e)** Diffraction-limited image of a cross-section at a z-plane in the middle of a lipid bilayer-coated silica microsphere labeled using the membrane dye Di-8-ANEPPS. **f)** POLCAM SMOLM reconstruction of a cross-section of a lipid bilayer-coated silica microsphere acquired through PAINT with Nile red. Each localization is drawn as a rod with a direction indicating the estimated in-plane angle *ϕ*. **g)** An experimental bias curve for the estimation of *ϕ* generated using a PAINT dataset such as the one shown in panel f. **h)** Illustration of the angles specifying the orientation of the emission dipole moment. **i-j)** Experimental precision from repeated localization and orientation estimation on AF647 immobilized in PVA. The precision is the measured standard deviation on repeated measurement of the position (x,y), and in-plane angle *ϕ* of the same molecule. Photon numbers are averages. Measurements between N = 12-40 are used to calculate the standard deviation. A power law was fitted to the data.

### Experimental accuracy and precision

To measure the experimental accuracy of orientation estimation, silica microspheres with a diameter of 5 μm were coated with a lipid bilayer (DPPC containing 40% cholesterol, Fig 2d) as in [33, 37] and labeled using different membrane dyes. Polarization-resolved diffraction-limited images were acquired using bulk labeling concentrations of different membrane dyes (Di-8-ANEPPS, DiI, and Nile red). As expected, the emission dipole moments of Di-8-ANEPPS and Nile red orient perpendicular to the membrane surface, and DiI orients parallel to the membrane surface (Fig. 2e, Supplementary Fig. S33). Nile red were used to collect a PAINT [4] dataset of a lipid-coated microsphere to generate an experimental accuracy curve for the in-plane angle *ϕ* (Fig. 2f-g).

Experimental orientation estimation and localization precision curves as a function of the number of detected photons were generated using single Alexa Fluor 647 (AF647) dyes immobilized in poly(vinyl alcohol) (PVA). The data points in figure 2i are the measured standard deviation on the repeated measurement of the orientation of an AF647 molecule. At 500 detected photons (the default lower threshold on photon number that is used in all datasets), we achieve an experimental in-plane angle precision *σ*_*ϕ*_ of 7.5° (Fig. 2i), *i*.*e*. the upper bound on *σ*_*ϕ*_. The precision converges to a lower bound of 1° at higher photon numbers (Fig. 2i). The experimental localization precision at 500 detected photons is 25-30 nm and converges to 5 nm at high photon numbers (Fig. 2j). The same curves generated using simulations with a realistic noise model largely agree with the experimental data (Supplementary Fig. S35). A complete characterization of the bias and precision on the position, orientation, and rotational mobility estimates based on simulations can be found in Supplementary figures S36–S41. DNA-origami with a spacing of 80 nm between binding sites can also be easily resolved using POLCAM (Supplementary Figure S31 and S32).

### TAB-PAINT imaging of alpha-synuclein fibrils *in vitro*

Orientationally-resolved imaging of *α*-synuclein fibrils labeled with the dye Nile red has previously been demonstrated using TAB-PAINT (Transient Amyloid Binding PAINT) [4, 65]. Nile red reversibly binds to the hydrophobic regions of the fibrils in a defined orientation [12, 35]. Therefore, this serves as an excellent test sample for orientation-resolved super-resolution imaging in biologically relevant samples. Figure 3 shows the POLCAM reconstruction of the *α*-synuclein fibrils, which were immobilised on a cover glass using a PLL-coating and imaged in PBS. POLCAM can super-resolve morphologically consistent *α*-synuclein fibrils with widths of *∼* 50 nm (FWHM) over large fields of views of *∼* 50 μm *×* 50 μm (Fig. 3a-b, Supplementary Fig. S34). When color-coding the reconstructions by the in-plane angle *ϕ* estimate, show that the majority of the Nile red molecules orient parallel to the long axis of the fibril (Fig. 3c-e). The distributions of the measured in-plane angle over short fibril sections have standard deviations around 7° and 9° (Fig. 3f), which approaches the expected precision for this dataset (Fig. 2i, *∼* 5° at 1000 detected photons). This indicates that the width of this distribution can not be explained by measurement precision alone, and must partly be due to Nile red molecules binding at a range of orientations that are not exactly parallel to the fibril axis.

**Figure 3:**
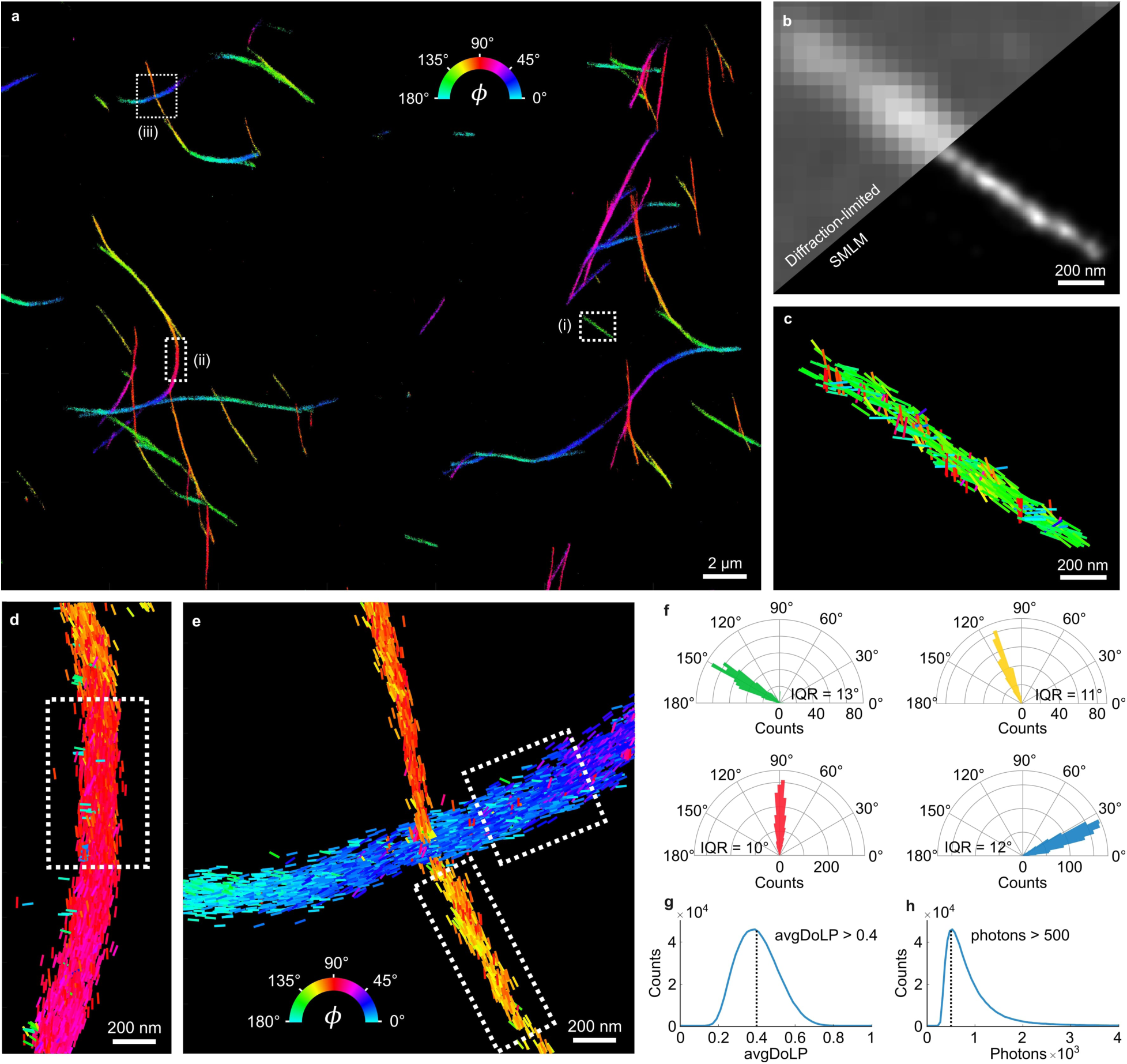
TAB-PAINT imaging of alpha-synuclein fibrils *in vitro*. **a)** A POLCAM SMOLM reconstruction of *α*-synuclein fibrils, color-coded by the in-plane angle *ϕ* of the emission dipole moment of the Nile Red molecules. **b)** Diffraction-limited image and SMLM reconstruction of inset (i) from panel a. **c-e)** Detail of insets (i), (ii), and (iii) in panel a, where individual localizations are drawn as rods. The orientation and color of the rods indicate the measured *ϕ*. **f)** Polar histograms of the fibrils that are shown in panel c (green), and segments indicated by the dashed boxes drawn in panels d (red) and panel f (yellow and blue). The interquartile range (IQR) of the distributions is displayed below the respective histograms. **g)** Histogram of the avgDoLP of the data shown in panel a, including the avgDoLP threshold used to exclude localizations with high rotational mobility when displaying *ϕ*-color-coded reconstructions. **h)** Histogram of the number of detected photons per molecule per frame for the dataset shown in panel a. The minimum number of detected photons (500 photons) is also indicated.

While a range of avgDoLP and photon values were extracted using POLCAM (Fig. 3g-h), we apply lower limit filtering thresholds to ensure high-accuracy results; (i) at least 500 detected photons (applied to all single-molecule data presented in this work), and (ii) avgDoLP > 0.4 when *ϕ*-color-coded data is shown, as localizations with avgDoLP < 0.4 are too rotationally free to estimate a meaningful orientation. An example TAB-PAINT dataset is included in the supporting information (Dataset 2).

### dSTORM imaging of actin in fixed HeLa cells

Next, to demonstrate the versatility of POLCAM, we performed SMOLM on eukaryotic cells using dSTORM. Fixed HeLa cells were labeled using phalloidin-Alexa Fluor 488 (AF488) and phalloidin-Alexa Flour 647 (AF647). In previous literature, this labeling has been shown to result in rotational restriction for AF488 (where the dyes on average align with the axis of actin fibers) and rotational freedom for AF647 [39, 44]. Single AF488 and AF647 molecules can be easily detected with POLCAM in the cellular environment, and the difference in rotational mobility between the rotationally constrained AF488 and randomly oriented AF647 is also directly evident from the images of single molecules (Fig. 4a-b). The difference in rotational mobility between the two labeling approaches is also visible in the resulting super-resolution images (Fig. 4c-d) using a colormap where more orientationally random areas appear white (avgDoLP < 0.4) and the more ordered appear colored. Analysis of the distributions of the avgDoLP revealed an ordered subset of localizations (Fig. 4e) (25 % for AF488 and only 5 % for AF648). By filtering these localizations using this empirically determined threshold (avgDoLP *>* 0.4), a refined POLCAM image of actin in cells can be generated that excludes localizations that are too rotationally free to generate an accurate *ϕ* estimate (Fig. 4g). The resolution of the super-resolved images was estimated using FRC (Fourier Ring Correlation [66]): 70 nm for the AF488 dataset and 55 nm for the AF647 dataset (Fig. 4f).

**Figure 4:**
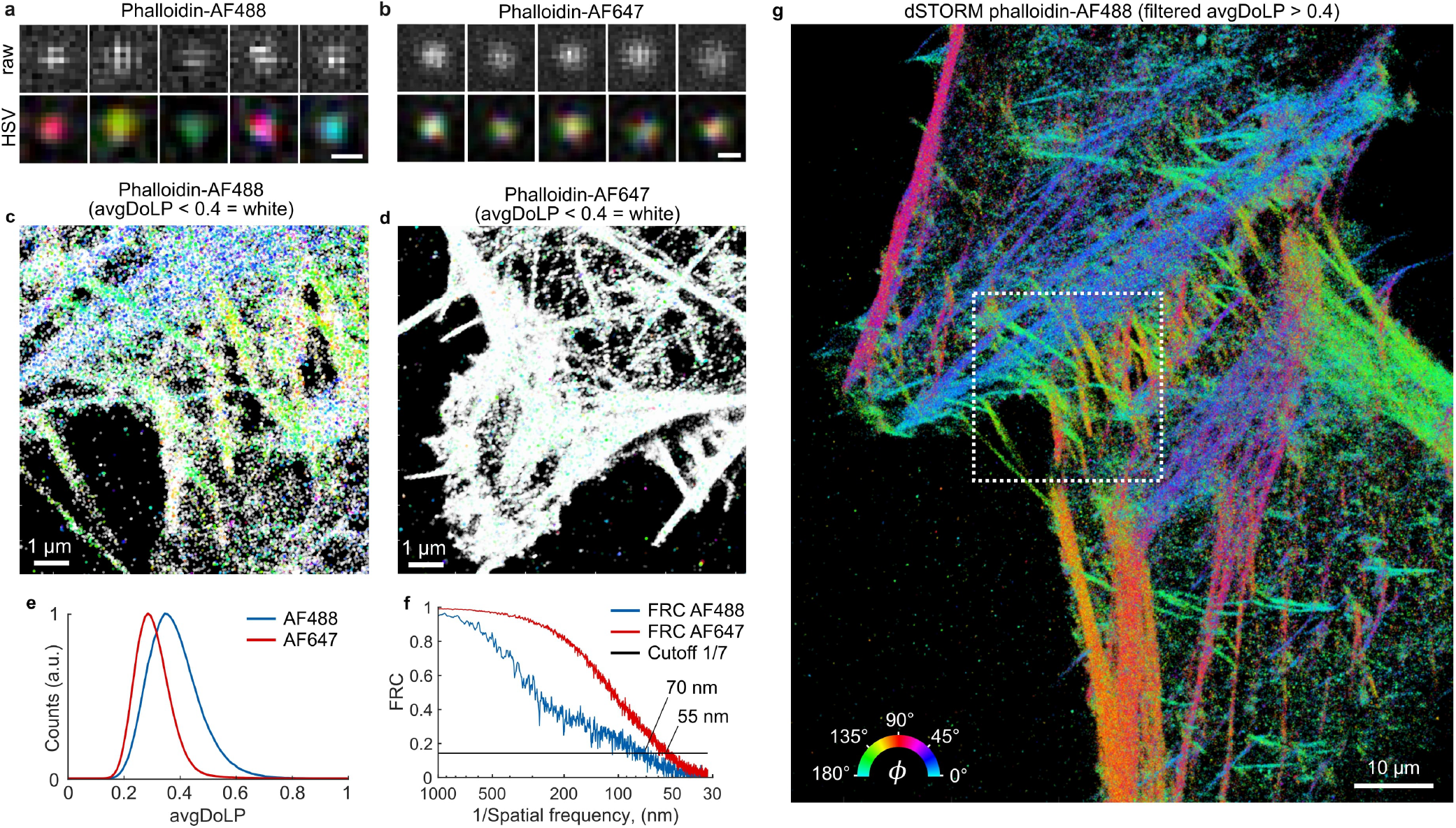
dSTORM imaging of actin in fixed HeLa cells. **a)** Representative examples of images of single AF488 molecules from the dSTORM datasets of phalloidin-AF488, unprocessed (top) and as an HSV (hue = AoLP, saturation = DoLP, value = S0) colormap (bottom). Scale bar 300 nm. **b)** The same as in panel a, but for AF647. Scale bar 300 nm. **c)** POLCAM SMOLM reconstruction of a phalloidin-AF488 dSTORM dataset in the form of a modified *ϕ*-color-coded scatter plot. All localizations with an avgDoLP *ϕ* 0.4 are colored white to indicate they have high rotational mobility. **d)** The same as panel c, but for phalloidin-AF647. **e)** Comparison of the avgDoLP distribution of single molecules from the phalloidin-AF488 and phalloidin-AF647 datasets. **f)** Fourier ring correlation curves for the phalloidin-AF488 (70 nm FRC resolution) and phalloidin-AF647 (55 nm FRC resolution) dSTORM datasets. **g)** *ϕ*-color-coded scatter plot of the full dataset from inset c (marked by the dashed box), displaying only the points with avgDoLP *>* 0.4 (*i*.*e*., localizations with high rotational mobility are not displayed).

### Improving accuracy by considering the DSF shape

From equation (47), it is clear that the estimation of the polar angle *θ* will become biased in the presence of rotational mobility, as netDoLP will decrease with increasing rotational mobility. Additionally, equation (47) becomes biased in the presence of noise, and depends on the refractive index of the sample medium.

As a result, unbiased estimation of *θ* is more reliably performed by additionally taking the shape of the DSF into account. We adapted the previously published DSF-fitting algorithm RoSE-O [67] for use with a polarization camera. This algorithm fits the shape of the image of a single emitter in all four polarized channels to estimate the orientation and rotational mobility of the emitter. Using simulated images, we compared the performance of the *intensity-only* algorithm and the DSF-fitting algorithm. As expected, the DSF-fitting algorithm is able to more accurately estimate *θ* in the presence of rotational mobility (Fig. 5a,b). For in-plane oriented emitters, both algorithms achieve the same localization precision across a wide range of signal-to-noise ratios (SNR), but the DSF-fitting algorithm outperforms the intensity-only algorithm up to 2-fold for emitters that are oriented out-of-plane (Fig. 5c). The precision on the estimation of *ϕ* is similar for in-plane oriented emitters, but the DSF-fitting algorithm is able to more precisely estimate *ϕ* for more out-of-plane oriented emitters, especially at high SNR (Fig. 5d). A complete comparison of the algorithms can be found in Supplementary figures S36–S41.

**Figure 5:**
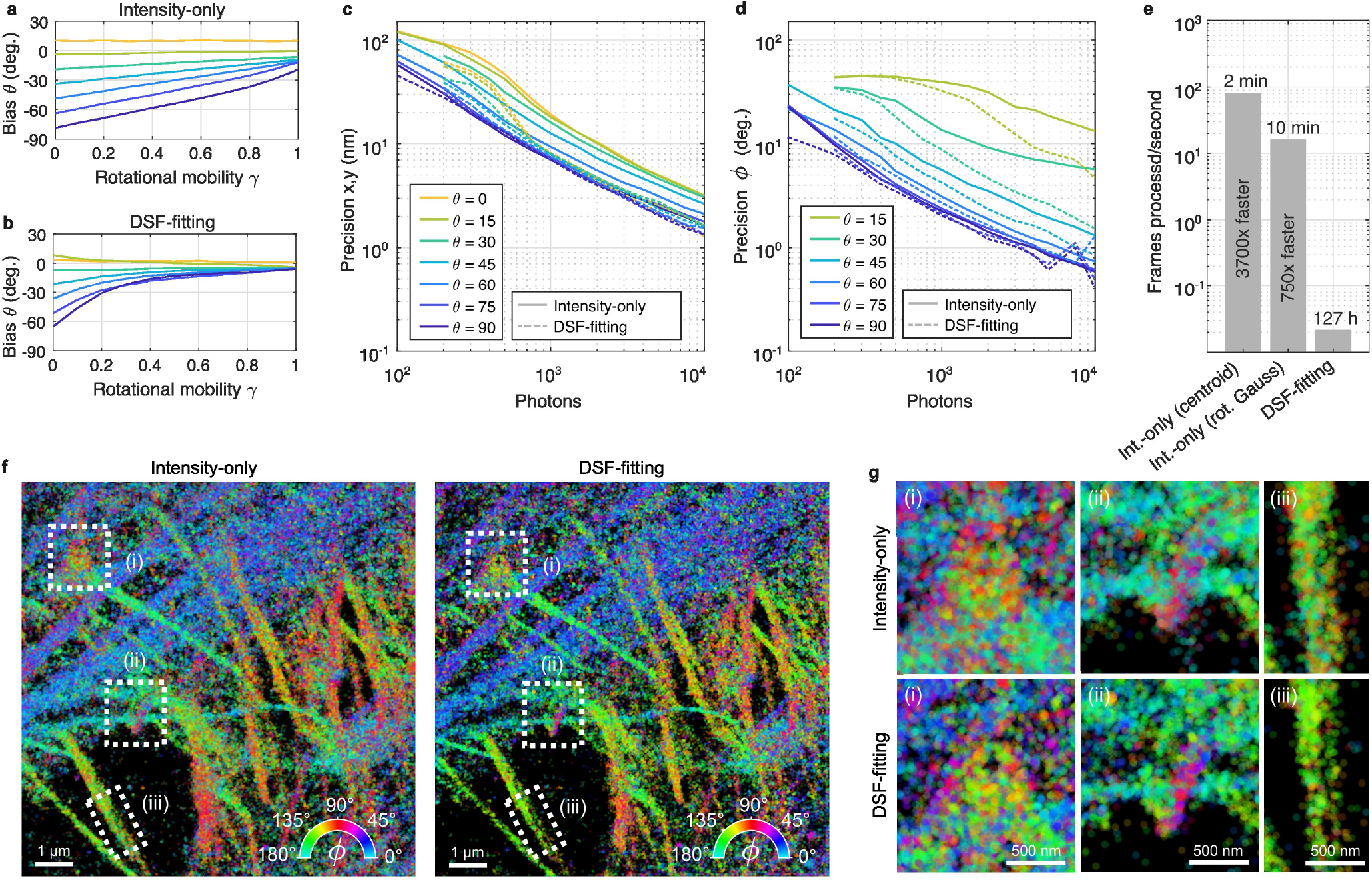
Improving accuracy and precision by considering the DSF shape. **a)** The bias on the estimation of the out-of-plane angle *θ* as a function of rotational mobility *γ* for the intensity-only algorithm, determined using simulated images of single dipole emitters (using 1000 photons/emitter, and 10 background photons/pixel). A rotational mobility parameter *γ* of 0 corresponds to total rotational freedom, and a value of 1 corresponds to perfect rotational immobilization. **b)** The same as panel a but for the DSF-fitting algorithm. **c)** The lateral localization precision as a function of the number of detected photons as determined from simulations (10 background photons/pixel, *γ* = 1). Separate curves are shown for molecules at different out-of-plane orientations. **d)** The same as panel c, but for the precision of *ϕ* estimation. **e)** The computer processing time for the datasets from panel f for the intensity-only algorithm with centroid localization, the intensity-only algorithm with least squares fitting of a rotated asymmetric Gaussian, and the DSF-fitting algorithm. Refer to the methods section for the computer specifications. **f)** A POLCAM SMOLM reconstruction of a 200*×*200 pixel region from figure 4 with 10.000 frames, as analyzed by the intensity-only algorithm (with rotated asymmetric Gaussian fitting) and the DSF-fitting algorithm. Both reconstructions are rendered using approximately the same number of localisations (258.251 localisations for the intensity-only algorithm, 258.172 for the DSF-fitting algorithm). **g)** Insets from the regions in panel f that are marked by dashed boxes. The insets display some structural differences between the reconstructions generated by the two algorithms.

Figure 5f shows super-resolution reconstructions generated by the two algorithms of a subset (200*×*200 pixels and 10,000 frames and *∼*15 localizations/frame) of the phalloidin-AF488 dSTORM dataset from Fig. 4g. Although both algorithms generate similar-looking reconstructions, zooming in on specific regions (Fig. 5g) seems to confirm that the DSF-fitting algorithm produces a slightly higher resolution image. A spatial comparison of the *ϕ* estimates shows that the widths of the *ϕ* distributions generated by the DSF-fitting algorithm are slightly more narrow (Supplementary Fig. S41). On a typical workstation (see Methods for system specifications), the *intensity-only* algorithm is *>* 750-3700 times faster (2 minutes total processing using centroid localization, and 10 minutes using least squares rotated asymmetric Gaussian fitting) compared to the DSF-fitting algorithm. Therefore, the DSF-fitting algorithm is recommended if the complete 3D orientation (*ϕ, θ*) needs to be estimated. If knowledge of *ϕ* and the rotational mobility proxy avgDoLP is sufficient, the fast *intensity-only* algorithm is recommended.

### Live diffraction-limited polarization microscopy

The combination of instrumental simplicity and fast computation makes POLCAM compatible with real-time image processing. We developed napari-polcam —a napari plugin for the open-source multi-dimensional image viewer napari [58] (Fig. 6a)—, and a standalone application for on-the-fly processing and rendering called POLCAM-Live (Fig. 6b). Both software take in unprocessed data and convert it into different formats in an easy-to-use interface. The ability to process data live is useful for fast decision-making during experiments and alignment (Supplementary Note S8).

**Figure 6:**
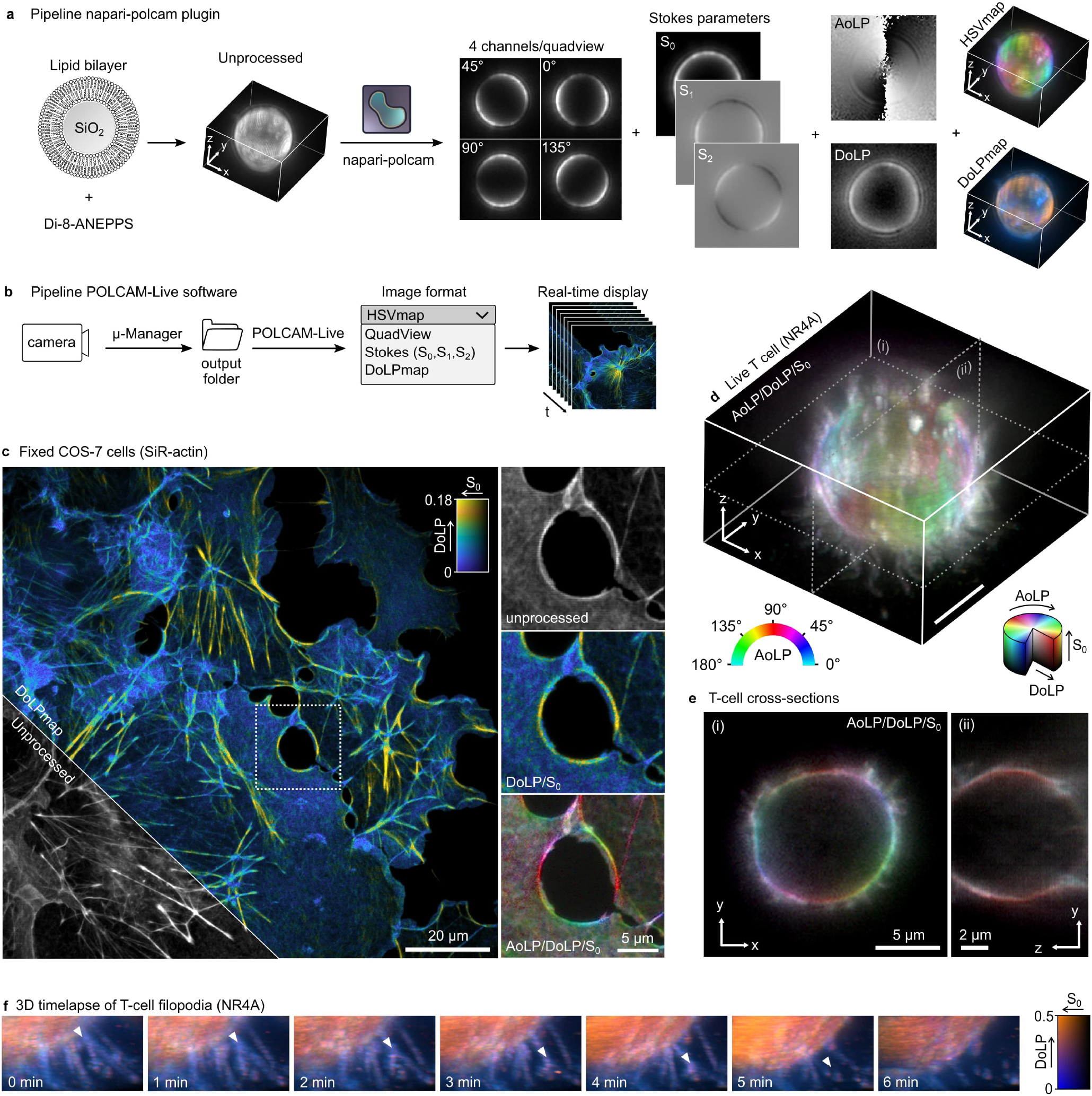
Diffraction-limited polarization microscopy using POLCAM. **a)** Image processing workflow of the napari plugin napari-polcam, demonstrated on an example 3D dataset of a lipid bilayer-coated silica microsphere labeled with the membrane dye Di-8-ANEPPS. **b)** Workflow of the POLCAM-Live software for real-time processing and rendering of polarization camera images during image acquisition. **c)** Diffraction-limited polarization camera image of fixed COS-7 cells labeled with SiR-actin, rendered using a DoLP colormap. An inset marked by a dashed square is shown in unprocessed, DoLP colormap and HSV polarisation colormap. **d)** A 3D image of the plasma membrane (using the dye NR4A) of a live T cell. **e)** Two cross-sections of the T cell shown in panel d. **f)** A 3D time-lapse of the movement of the filopodia of a live T cell rendered using a DoLP colormap. Randomly polarised regions appear blue (filopodia) and more structured and therefor polarised areas (larger, more smooth sections of the plasma membrane surface) appear orange. A white triangle tracks the movement of what appears to be a branching point on a filopodium.

We use these software tools to illustrate diffraction-limited polarization imaging, demonstrating 3D time-lapse imaging of the plasma membrane in live human T cells using the new probe NR4A [68]. The 3D images are acquired by axially scanning the objective. The filopodia of the T cells appear more unpolarized (Fig. 6d-f), allowing their simple identification from cell bodies, which is the subject of intense research interest with regard to surface receptor organisation [69]. The unpolarized appearance of filopodia is likely due to their small diameter. The 3D motion of the filopodia can be tracked over time (*∼*1 min/volume, limited by the speed of our z-stage). Furthermore, POLCAM can be used in a simplified mode to discriminate ordered vs non-ordered actin structures based on the DoLP. Using a simple DoLP-colormap, we can distinguish highly ordered regions (yellow) from more disordered regions (blue) in the actin network in COS-7 cells labeled with SiR-actin (Fig. 6c).

## Discussion

In this work, we present POLCAM, a new method for molecular orientation-resolved fluorescence microscopy that makes use of a polarization camera to dramatically simplify the experimental setup. The method can be used in two modes —single-molecule orientation localization microscopy (SMOLM) or diffraction-limited polarization fluorescence microscopy— and can be implemented by simply replacing the conventional detector on a widefield fluorescence microscope with a polarization camera. The polarization camera used in this work is supported by the popular image acquisition software Micro-Manager [57]. Furthermore, we provide software to improve widespread user adaption, as well as a comprehensive installation guide, and a software user manual.

We present two SMOLM analysis algorithms for POLCAM; 1) a fast algorithm based on Stokes parameter estimation and simple intensity measurements for robust in-plane angle estimation compatible with high-throughput data collection, and 2) an unbiased 3D orientation estimation algorithm that fits a DSF model. Due to the computational cost of the DSF-fitting algorithm, it is more suited to the analysis of small datasets.

In conventional four-channel polarized detection where beam splitters and polarization optics are used to separate the fluorescence into four channels, it is more likely that channel-dependent aberrations will occur. Due to the simplicity of the experimental setup of POLCAM, this is avoided, simplifying the use of a DSF fitting algorithm. Moreover, there is no need for channel registration and localization grouping because single-molecule localization is performed on the estimated incident intensity (*S*_0_) on the micropolarizer array, meaning that all in-plane orientations are equally detectable and no localizations are therefore missed.

A diffraction-limited polarization colormap (HSVmap) can be generated and displayed in real-time during experiments, for which we provide the standalone software POLCAM-Live. In theory, the intensity-only SMOLM algorithm is also fast enough to be compatible with real-time processing, as a 200*×*200 pixel image (10*×*10 μm) with *∼*15 localizations takes about 10 ms to process, which is less than typical camera exposure times. This could enable molecular orientation event-triggered microscopy [70, 71], something that would be extremely challenging if DSF-fitting is required.

Because a polarization camera uses polarizers to achieve polarized detection, on average the system is 50% efficient, as half of the photons that are captured by the objective will be absorbed by polarizers. Nevertheless, we demonstrate in this work that this approach is compatible with a wide variety of fluorophores (*e*.*g*., Alexa Fluor 488, Nile Red, SiR, Cy5, SYTOX Orange). The current generation of polarization cameras lacks the sophistication of modern sCMOS cameras in areas such as the quantum efficiency of the detector (70% *vs*. 95%) and the presence of onboard denoising algorithms. We expect this to improve over time. Further improvements to the analysis approach can be made by performing a pixel-dependent characterization of any defects of the micropolarizer array or deviation from an ideal micropolarizer array [72].

The success of a molecular-orientation resolved experiment stands and falls with the labeling approach. It would thus be meaningless to attempt to measure molecular orientation if the relative orientation of a target and the probe is random. Therefore, the use of antibodies with multiple dyes conjugated at random orientations (*e*.*g*., random lysine labeling), or staining using secondary antibodies are likely not suitable. Currently, the number of labeling protocols that restrict or control the orientation and rotational mobility of a fluorophore with respect to their target is still limited and there is a strong need for the development of more labeling approaches and the discovery of suitable probes. Examples are the use of bifunctional Rhodamines [16], and the genetically engineered rigid protein linker POLArIS [73]. It is likely that many common fluorescent probes are suitable for molecular orientation-resolved microscopy, but they have simply never been evaluated for this purpose. We anticipate that because of the accessibility and compatibility with high-throughput data acquisition, POLCAM will accelerate this much-needed development and discovery of new probes and expand the current toolkit to cover more biological systems.

We note that polarization cameras can also be used for label-free microscopy [74–77], leaving the possibility for multiplexing of fluorescence and label-free techniques. Overall, we envisage that the combination of POLCAM’s simple implementation and ease of use, computational speed, and open-source software will lead to new biological insight across diverse systems.

## Acknowledgements

This research was funded in part by Aligning Science Across Parkinson’s ASAP-000509 through the Michael J. Fox Foundation for Parkinson’s Research (MJFF). For the purpose of open access, the author has applied a CC BY public copyright license to all Author Accepted Manuscripts arising from this submission. R.B. was funded by the ERC (European Research Council Consolidator Grant 820188-NanoMechShape to E.K.P.). The authors thank the Cambridge Advanced Imaging Centre (CAIC, University of Cambridge) for the use of a microscope for the dSTORM experiments, and the mechanical workshop (Yusuf Hamied Department of Chemistry, University of Cambridge) for building custom parts of optical setups used in this work. We thank Dr. Aleks Ponjavic and Dr. Joseph Beckwith for critical reading of the manuscript. We thank Laila Elfeky for testing the POLCAM-SR software.

## Author Contributions Statement

SFL conceived the project. SFL, KOH and MDL supervised the research. EB built the optical setups. EB acquired the data. EB programmed and performed the simulations. EB and OZ analysed data. EB programmed the software POLCAM-SR, POLCAM-Live and napari-polcam. OZ adapted the RoSE-O algorithm for polarization cameras. LMN prepared the alpha-synuclein fibrils and dye in polymer samples. RP prepared the dSTORM actin samples. MK prepared the live T-cell samples. EB, MC and SD prepared the lipid bilayer-coated microsphere samples. TW provided the protocol for preparing lipid bilayer-coated microspheres. ASK provided the dye NR4A. SJD provided the J8 LFA-1 cell line and OKT3 antibodies. EB and SFL wrote the manuscript with input from all authors.

## Methods

### Optical setups

Experiments were performed on three very similar widefield fluorescence microscopes. The SYTOX orange, AF647 immobilized in polymer, TAB-PAINT of alpha-synuclein fibrils, and diffraction-limited polarisation imaging of the actin network in COS-7 cells were performed on ‘Microscope 1’. The dSTORM experiments were performed on ‘Microscope 2’. The lipid bilayer-coated silica microsphere and live T cell experiments were performed on ‘Microscope 3’.

‘Microscope 1’ is a widefield fluorescence microscope (Eclipse Ti-U, Nikon), with the illumination entering the microscope body through the back illumination port. The beams from three free space lasers (515 nm, 150 mW, Spectra Physics; 532 nm, 120 mW, Odic Force Lasers; and 638 nm, 350 mW, Odic Force Lasers) were expanded, spectrally and spatially filtered, combined with dichroics, and focused to a spot in the back focal plane of an oil immersion objective (Plan Apo, 60*×*A/1.40 oil, DIC H, *∞*/0.17, WD 0.21, Nikon) using an achromatic doublet lens (AC254-300-A, Thorlabs). This lens and a periscope were mounted on a linear translation stage to allow manual adjustment of the beam emerging from the objective and switch between EPI, HILO and TIRF illumination. Approximately circular polarization at the sample plane was achieved using quarter waveplates (WPQ10M-514, WPQ10M-532, and WPQ10M-633, Thorlabs). For diffraction-limited imaging of the actin network in COS-7 cells, a different laser source was used with a multimode fiber (LDI-7 Laser Diode Illuminator, 89North). Fluorescence was filtered by a dichroic beamsplitter (Di03-R405/488/532/635-t1 for AF647, and Di03-R532-t1 for SYTOX Orange and Nile Red, Semrock) and emission filters (BLP01-532R for SYTOX Orange; BLP01-635R for AF647; BLP01-532R and FF01-650/200 for Nile red, Semrock). The fluorescence was focused on a polarization camera (CS505MUP, Thorlabs) that was placed directly at the microscope body camera port. The pixel size of the camera is 3.45μm *×* 3.45μm, resulting in a virtual pixel size of 57.5 *×* 57.5 nm. The microscope PC was a Dell OptiPlex 7070 Mini Tower running on Windows 10 (64 bit), with an Intel® Core™ i9-9900 processor and 32 GB RAM.

‘Microscope 2’ is functionally similar to ‘Microscope 1’, with a different microscope body (Eclipse Ti-E, Nikon), laser source (Omicron LightHUB with 405, 488, 561 and 638 nm lasers, single-mode fiber, collimator RC08APC-P01, Thorlabs) and a 60*×* 1.42 NA oil immersion objective (Olympus, PlanApo N). A quadband imaging dichroic (Di03-R405/488/532/635-t1, Semrock) and emission filters were used (BLP01-488R and FF01-582/64 for AF488; BLP01-635R for AF647, Semrock). The fluorescence was focused on a polarization camera (CS505MUP, Thorlabs) that was placed directly at the microscope body camera port, resulting in a virtual pixel size of 51.7 *×* 51.7 nm. Note that because an Olympus objective (assumes use of an Olympus body with a 180.0 mm focal length tube lens) was used with a Nikon body (200.0 mm focal length tube lens).

‘Microscope 3’ is also functionally similar to ‘Microscope 1’, other than the microscope body (Eclipse Ti-E, Nikon), lasers (Cobolt C-FLEX combiner with 405, 488, 515, 561 and two 638 nm lasers, free space) coupled into a square-core multi-mode fiber (M97L02, Thorlabs) with a custom vibration-motor based mode scrambler, and a 100*×* 1.49 NA oil immersion objective. A 4f system consisting of two achromatic lenses (AC254-050-A-ML and AC254-100-A-ML, Thorlabs) was included in the emission path to demagnify the image 2*×*, resulting in a total system magnification of 50*×* and thus virtual pixel size of 69 *×* 69 nm. Imaging dichroics (and Di03-R515-t1 for NR4A and Nile red, and Di03-R405/488/532/635-t1 for SiR-actin; Semrock) and emission filters (BLP01-532R and FF01-650/200 for NR4A, BLP01-635R for SiR-actin; Semrock) were used. A multi-mode fiber was used for all diffraction-limited imaging experiments as this allowed us to achieve highly randomized polarization at the sample plane (compared to using a quarter-wave plate), resulting in negligible photoselection.

### Simulations

The emission of a fluorescent molecule was modeled as the far-field of an oscillating electric dipole as previously described in [1]. The transmission of the emission through the micropolarizer array on the camera sensor was modeled for each camera pixel using a Jones matrix *J*_*LP*_ for a linear polarizer with an axis of transmission at an angle *η* from the x-axis [2]

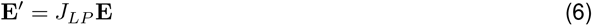

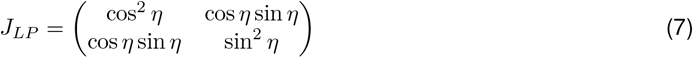

Unless otherwise specified, the following system parameters were used: 630 nm emission wavelength, 60*×* oil-immersion objective with a numerical aperture of 1.4, a tube lens with a focal length of 200.0 mm, physical camera pixel size of 3.45 μm. The molecules are placed on a glass-water refractive index interface (*𝓃*_glass_ = *𝓃*_oil_ = 1.518, *𝓃*_water_ = 1.33) and in focus. A CMOS camera noise model was used (Supplementary Note S1.7).

### Image acquisition

On all instruments, image acquisition was performed using Micro-Manager [3] (Micro-Manager v2.0.0, http://micro-manager.org, RRID:SCR 000415), or Thorcam (Thorcam v3.6.0, https://www.thorlabs.com/software_pages/ViewSoftwarePage.cfm?Code=ThorCam). Data was always recorded in unprocessed format. All experiments on all three microscopes were performed using epi-illumination (with the exception of the dSTORM experiment where a steep HILO angle was used) to easily control the polarization state at the sample plane. Perfectly random or circular polarization at the sample plane is very challenging to achieve experimentally, so to some degree there will always be a dominant axis of polarization. Transitioning from EPI to HILO and TIRF will change the amount of photoselection at the sample plane as the dominant axis and degree of randomness/ellipticity of the polarization will change in a way that is challenging to quantify and reproduce. Before each experiment and/or change of the inclination of the excitation beam, the polarization of the excitation beam at the sample is tuned by rotating a quarter wave plate. Alignment is deemed optimal when the DoLP of the background is minimized (or the appearance of gridding distinctive of polarized background disappears). This step is not necessary if lasers are coupled into a multi-mode fiber. The imaging parameters for the data shown in all main figures and supplementary figures is summarised in Supplementary Table 3.

### Image analysis

Single-molecule data analysis was performed using the MATLAB (MATLAB R2022a, Mathworks, http://www.mathworks.com/products/matlab/, RRID:SCR 001622) application POLCAM-SR, which includes tools for localization, filtering, drift correction, and data visualization. The source code and installer are available on GitHub at https://github.com/ezrabru/POLCAM-SR. Diffraction-limited data analysis was performed using POLCAM-SR and a napari [4] plugin called napari-polcam for processing and visualization of multi-dimensional polarization camera datasets. The source code and installation instructions for napari-polcam are available on GitHub at https://github.com/ezrabru/napari-polcam. Refer to Supplementary Note S7 for a detailed description of the image analysis pipeline.

### SYTOX orange on cover glass

Glass coverslips (VWR collection, 631-0124) were Argon plasma cleaned for 30 min (Expanded Plasma Cleaner, PDC-002, Harrick Plasma). An imaging chamber was created on the coverslips using frame-seal slide chambers (9×9 mm, SLF0201, Bio-rad). The glass in the chamber was coated with 70 μl of poly-L-lysine (PLL, 0.01 % w/v, P4707, Sigma-Aldrich) for 15 min. After removing excess PLL and washing 3 times with filtered PBS (0.02 μm syringe filter, Whatman, 6809-1102), 50 μl of 1 nM SYTOX Orange (S11368, Invitrogen) was added gently. The sample was imaged straight away.

This protocol is available on Protocols.io as *Imaging single SYTOX Orange molecules on a PLL-coated cover glass* [5].

### PAINT imaging of lipid bilayer-coated silica microspheres

To prepare the lipid bilayer-coated silica microspheres, a slightly modified version of the protocol in [6] was used. First, lipid vesicles of a certain composition are prepared as follows: DPPC (850355C, Avanti Polar Lipids) and cholesterol (C8667-5G, Sigma-Aldrich) were dissolved in chloroform (366927, Sigma-Aldrich) to respectively 25 mg/ml and 10 mg/ml. A DPPC/40%chol mixture was prepared by combining 23 μl DPPC and 20 μl cholesterol. The solvent was evaporated overnight under vacuum. The lipid/cholesterol mixture was rehydrated using 1 ml of Tris-Ca^2+^ buffer (100 mM NaCl, 3 mM CaCl_2_, 10 mM Tris base, pH 7.4) and vortexed for 30 seconds. The solution was sonicated using a tip sonicator (cycles of 45 s on, 15 s off, 60% amplitude) for 40 minutes until the solution ran clear. The sonicated solution was centrifuged for 90 s at 14.000 rcrf to remove titanium residue from the sonicator probe.

Next, silica microspheres with a diameter of 5μm (44054-5ML-F, Sigma-Aldrich) were diluted to approximately 2.8 mg/ml, and cleaned by centrifuging and replacing the stock buffer with Tris-Ca^2+^. The micro-spheres and lipid vesicle solutions were heated to 65°C using a heated water bath and mixed together in a 1:1 ratio. After 30 minutes at 65°C, the mixtures were slowly cooled down to room temperature (but turning the heating bath off). The buffer was gradually replaced by Tris (100 mM NaCl, 10 mM Tris base, pH 7.4) by centrifugation (5 minutes at 0.3 rcf) and replacement of two thirds of the supernatant with Tris, repeated 6 times. The lipid-coated microspheres were stored at 4°C and used within less than 2 weeks of preparation. For imaging, the lipid-coated microspheres were added to an Argon plasma-cleaned, PLL-coated cover glass (VWR collection, 631-0124), and 1 nM of NR4A or Nile red was added for PAINT imaging of the lipid bilayer. The Nile red derivative NR4A was provided by Andrey S. Klymchenko at Universitéde Strasbourg. For diffraction-limited imaging of lipid-coated microspheres, 100 nM dye in PBS was used (for Nile red, NR4A, and Di-8-ANEPPS). All buffers (PBS, Tris, Tris-Ca^2+^) were filtered before use (0.02 μm syringe filter, Whatman, 6809-1102).

This protocol is available on Protocols.io as *Preparation and imaging of lipid bilayer-coated silica micro-spheres* [7].

### AF647 immobilized in PVA

A polyvinyl alcohol solution (PVA) (1%, 1 g in 100 ml) was prepared by slow addition of solid PVA into filtered (0.02 μm syringe filter, Whatman, 6809-1102) milli-Q water with stirring. The solution was then heated to 90°C and stirred for 30 minutes, then removed from the heat with continued stirring for 12 hours. The PVA solution was then filtered (0.02 μm syringe filter, Whatman, 6809-1102) and stored at 4°C. Alexa Fluor 647 (500 pM) was diluted in the 1% PVA solution, 10 μl was then spin-cast (3000 rpm, 45 s) onto an Ar plasma-cleaned glass coverslip (ODC-002, Harrick Plasma) and sealed prior to imaging.

This protocol is available on Protocols.io as *Imaging single AF647 molecules immobilised in PVA on a cover glass* [8].

### TAB-PAINT of alpha-synuclein fibrils

To prepare alpha-synuclein fibrils, alpha-synuclein monomer was diluted to 70 μM concentration in PBS (with 0.01% NaN_3_) and incubated at 37°C in a shaker (200 rpm) to aggregate for ¿24 h. To prepare fibrils for imaging, glass coverslips (VWR collection, 631-0124) were plasma cleaned for 1 h (argon plasma cleaner, PDC-002, Harrick Plasma). An imaging chamber was created on the coverslips using frame-seal slide chambers (9×9 mm, SLF0201, Bio-rad). The glass in the chamber was coated with 70 μl of poly-L-lysine (PLL, 0.01 % w/v, P4707, Sigma-Aldrich) for 30 min. After removing excess PLL and washing 3 times with filtered PBS (20 nm pore filters), 50 μl Tetraspeck beads (0.1 μM stock, 10x diluted) were added for lateral drift correction. Samples were washed again 3x with filtered PBS and 50 μl of alpha-synuclein fibrils (diluted to 35 mM monomer concentration, from a 70 mM monomer concentration stock that was stored at 4 degrees). Fibrils were stuck to the PLL by pipetting up and down a couple of times in the four corners of the chamber. Before imaging, excess solution is removed, followed by a gentle wash with filtered PBS. 50 μl of the imaging buffer (1 nM Nile Red in PBS, diluted from a 1 mM aliquot in DMSO, stored at −20°C) is added and the sample is imaged straight away.

This protocol is available on Protocols.io as *TAB-PAINT imaging of alpha-synuclein fibrils using Nile Red* [9].

### dSTORM of actin in fixed HeLa cells

HeLa TDS cells (RRID:CVCL 0030) were cultured in DMEM (Gibco, Invitrogen) supplemented with 10% Fetal Bovine Serum (FBS, Life Technologies), 1% penicillin/streptomycin (Life Technologies), and 1% glutamine (Life Technologies) at 37°C + 5% CO_2_. Cells were periodically tested for mycoplasma contamination and passaged 3 times per week. Cells were plated at low density on high-precision glass coverslips (Mat-Tek, P35G-0.170-14-C) 1 day prior to fixation for dSTORM experiments.

Cells were simultaneously fixed and permeabilized in cytoskeleton buffer (CBS, 10 mM MES, 138 mM KCl, 3 mM MgCl_2_, 2 mM EGTA, 4.5% sucrose w/v, pH 7.4), + 4% paraformaldehyde (PFA) and 0.2% Triton for 6 minutes at 37°C, and further fixed in CBS + 4% PFA for 14 minutes at 37°C. Post-fixation, cells were washed x3 in PBST (PBS supplemented with 0.1% Tween) and permeabilized a second time in PBS + 0.5% Triton for 5 minutes at room temperature. The samples were then washed 3 times in PBST and blocked for 30 minutes in 5% BSA. Cells were washed x3 in PBST and then incubated with Alexa Fluor™ 488 Phalloidin (A12379, Invitrogen, 1:50 in PBS) for 1 h in the dark, followed by x2 washes in PBS. Prior to dSTORM imaging, PBS was replaced with dSTORM imaging buffer (base buffer consisting of 0.56 M glucose, 50 mM Tris (pH 8.5), and 10 mM NaCl supplemented with 5 U/mL pyranose oxidase (Sigma, P4234), 10 mM cysteamine (Sigma, 30070), 40 μg/mL catalase (Sigma, C100) and 2 mM cyclooctatetraene (Sigma, 138924).

This protocol is available on Protocols.io as *dSTORM of actin in fixed HeLa cells* [10].

### Fixed COS-7 cells labeled with SiR-actin

A commercial slide (GATTA-Cells 4C, Gattaquant) was used, containing fixed COS-7 cells (RRID:CVCL 0224) labeled with SiR-actin (and DAPI, Tom20 with Alexa Fluor 488, and *α*-Tubulin with Alexa Fluor 555.)

### Live-cell imaging of the plasma membrane of Jurkat T cells

J8 LFA-1 cells were incubated overnight (*∼*18 h) in complete-RPMI (StableCell RPMI-1640 media, Sigma) supplemented with 10% (v/v) fetal calf serum (FCS), 1% (v/v) HEPES buffer, and 1% (v/v) pen/strep antibiotics. 1 mL of cells were collected by centrifugation and resuspended in phenol-red free RPMI supplemented with 1% HEPES.

Round coverslips were rinsed with IPA, MilliQ, dried, and Ar-plasma cleaned for 20 minutes. Grace Bio-Labs CultureWells were attached, and the slide was incubated with OKT3 antibody (provided by the Human Immunology Unit, WIMM, Oxford) for 30 minutes. The slide was washed 5 times with phenol-red free RPMI supplemented with 1% HEPES and a final wash with phenol-red free RPMI supplemented with 1% HEPES and 200 nM NR4A (MemGlow™ NR4A Membrane Polarity Probe, Cat. #MG06, Cytoskeleton Inc.) before imaging.

This protocol is available on Protocols.io as *Live-cell imaging of the plasma membrane of Jurkat T cells* [11].

## Data Availability

The datasets generated as part of this study were uploaded to Zenodo: pixel-dependent camera calibration results (https://doi.org/10.5281/zenodo.10578307), single SYTOX Orange on a cover glass (https://doi.org/10.5281/zenodo.10469322), PAINT data of single Nile red dyes binding to lipid bilayer-coated silica microspheres (https://doi.org/10.5281/zenodo.10469444), TAB-PAINT data (https://doi.org/10.5281/zenodo.10470795), dSTORM phalloidin-AF488 (https://doi.org/10.5281/zenodo.10470982), dSTORM phalloidin-AF647 (https://doi.org/10.5281/zenodo.10732697) and T cells (https://doi.org/10.5281/zenodo.10471496). The datasets generated as part of this study are available from the corresponding author upon request.

## Code Availability

The installers and source code of all custom code and software used in this work are available on GitHub and Zenodo: POLCAM-SR (https://github.com/ezrabru/POLCAM-SR, https://zenodo.org/doi/10.5281/ zenodo.10732422), POLCAM-Live (https://github.com/ezrabru/POLCAM-Live, https://zenodo.org/doi/10.5281/zenodo.10732437), napari-polcam (https://github.com/ezrabru/napari-polcam, https://zenodo.org/doi/10.5281/zenodo.10732441), RoSE-O-POLCAM (https://github.com/Lew-Lab/RoSE-O_polCam), and the camera calibration software (https://github.com/TheLeeLab/cameraCalibrationCMOS, https://zenodo.org/doi/10.5281/zenodo.10732469).

## References

1. Betzig, E. et al. Imaging intracellular fluorescent proteins at nanometer resolution. Science 313(5793), 1642–1645 (2006).

2. Rust, M., Bates, M. & Zhuang, X. Sub-diffraction-limit imaging by stochastic optical reconstruction microscopy (STORM). Nature Methods 3(10), 793–795 (2006).

3. Hess, S., Girirajan, T. & Mason, M. Ultra-high resolution imaging by fluorescence photoactivation localization microscopy. Biophysical Journal 91(11), 4258–4272 (2006).

4. Sharonov, A. & Hochstrasser, R. Wide-field subdiffraction imaging by accumulated binding of diffusing probes. PNAS 103(50), 18911–18916 (2006).

5. Xu, K., Zhong, G. & Zhuang, X. Actin, spectrin, and associated proteins form a periodic cytoskeletal structure in axons. Science 339, 452–456 (2013).

6. Szymborska, A. et al. Nuclear pore scaffold structure analyzed by super-resolution microscopy and particle averaging. Science 341, 655–658 (2013).

7. Mund, M. e. a. Systematic nanoscale analysis of endocytosis links efficient vesicle formation to patterned actin nucleation. Cell 174, 884–896 (2018).

8. Shroder, D., Lippert, L. & Goldman, Y. Single molecule optical measurements of orientation and rotations of biological macromolecules. Methods Appl. Fluoresc. 4, 042004 (2016).

9. Sosa, H., Peterman, E. J., Moerner, W. E. & Goldstein, L. S. ADP-induced rocking of the kinesin motor domain revealed by single-molecule fluorescence polarization microscopy. Nat. Struct. Biol. 8, 540– 544 (2001).

10. Lippert, L. et al. Angular measurements of the dynein ring reveal a stepping mechanism dependent on a flexible stalk. PNAS 114(23), E4564–E4573 (2017).

11. Backer, A. S. et al. Single-molecule polarization microscopy of DNA intercalators sheds light on the structure of S-DNA. Sci. Adv. 5, eaav1083 (2019).

12. Ding, T., Wu, T., Mazidi, H., Zhang, O. & Lew, M. Single-molecule orientation localization microscopy for resolving structural heterogeneities within amyloid fibrils. Optica 7(6), 602–607 (2020).

13. Lu, J., Mazidi, H., Ding, T., Zhang, O. & Lew, M. Single-Molecule 3D Orientation Imaging Reveals Nanoscale Compositional Heterogeneity in Lipid Membranes. Angew. Chem. Int. Ed. 59, 17572 (2020).

14. Corrie, J. E. T. et al. Dynamic measurement of myosin light-chain-domain tilt and twist in muscle contraction. en. Nature 400, 425–430. https://www.nature.com/articles/22704 (1999).

15. Forkey, J. N., Quinlan, M. E., Alexander Shaw, M., Corrie, J. E. T. & Goldman, Y. E. Three-dimensional structural dynamics of myosin V by single-molecule fluorescence polarization. en. Nature 422, 399–404. https://www.nature.com/articles/nature01529 (2023) (Mar. 2003).

16. Beausang, J. F., Sun, Y., Quinlan, M. E., Forkey, J. N. & Goldman, Y. E. Fluorescent Labeling of Calmodulin with Bifunctional Rhodamine. Cold Spring Harbor protocols 2012, 10.1101/pdb.prot069351pdb.prot069351. ISSN: 1940-3402. https://www.ncbi.nlm.nih.gov/pmc/articles/PMC3852408/ (2023) (May 2012).

17. Chen, C. et al. Elongation factor G initiates translocation through a power stroke. Proceedings of the National Academy of Sciences 113, 7515–7520. https://www.pnas.org/doi/10.1073/pnas.1602668113 (2023) (July 2016).

18. Novotny, L. & Hecht, B. Principles of Nano-Optics (Cambridge University Press, New York, 2007).

19. Backer, A., Backlund, M., Lew, M. & Moerner, W. Single-molecule orientation measurements with a quadrated pupil. Optics letters 38, 1521 (2013).

20. Backer, A. & Moerner, W. Extending single-molecule microscopy using optical fourier processing. J Phys Chem B 118, 8313–8329 (2014).

21. Hafi, N. et al. Fluorescence nanoscopy by polarization modulation and polarization angle narrowing. Nature Methods 11, 579–584. (2024) (May 2014).

22. Zhanghao, K. et al. Super-resolution dipole orientation mapping via polarization demodulation. Light: Science & Applications 5, e16166–e16166. (2024) (Oct. 2016).

23. Guan, M. et al. Polarization modulation with optical lock-in detection reveals universal fluorescence anisotropy of subcellular structures in live cells. en. Light: Science & Applications 11, 4. (2024) (Jan. 2022).

24. Aguet, F., Geissbühler, S., Märki, I., Lasser, T. & Unser, M. Super-resolution orientation estimation and localization of fluorescent dipoles using 3-D steerable filters. Opt. Express 17, 6829–6848 (2009).

25. Mortensen, K., Churchman, L., Spudich, J. & Flyvbjerg, H. Optimized localization-analysis for single-molecule tracking and super-resolution microscopy. Nature Methods 7(5), 377–381 (2010).

26. Backer, A. S. & Moerner, W. E. Determining the rotational mobility of a single molecule from a single image: a numerical study. Optics Express 23(4), 4255–4276 (2015).

27. Böhmer, M. & Enderlein, J. Orientation imaging of single molecules by wide-field epifluorescence microscopy. J Opt Soc Am B 20(3), 554–559 (2003).

28. patra, D., Gregor, I. & Enderlein, J. Image analysis of defocused single-molecule images for three-dimensional molecule orientation studies. J Phys Chem A 108, 6836–6841 (2004).

29. Toprak, E. et al. Defocused orientation and position imaging (DOPI) of myosin V. PNAS 103(17), 6495–6499 (2006).

30. Backlund, M. et al. Simultaneous, accurate measurement of the 3D position and orientation of single molecules. PNAS 109(47), 19087–19092 (2012).

31. Backer, A., Backlund, M., von Diezmann, A., Sahl, S. & Moerner, W. A bisected pupil for studying single-molecule orientational dynamics and its application to three-dimensional super-resolution microscopy. Applied Physics Letters 104, 193701 (2014).

32. Zhang, O., Lu, J., Ding, T. & Lew, M. Imaging the three-dimensional orientation and rotational mobility of fluorescent emitters using the tri-spot point spread function. Appl. Phys. Lett. 113, 031103 (2018).

33. Wu, T., Lu, J. & Lew, M. D. Dipole-spread-function engineering for simultaneously measuring the 3D orientations and 3D positions of fluorescent molecules. Optica 9, 505–511 (2022).

34. Hulleman, C. N. et al. Simultaneous orientation and 3D localization microscopy with a Vortex point spread function. Nature Communications 12, 5934. 10.1038/s41467-021-26228-5 (Oct. 2021).

35. Ding, T. & Lew, M. D. Single-Molecule Localization Microscopy of 3D Orientation and Anisotropic Wobble Using a Polarized Vortex Point Spread Function. Journal of Physical Chemistry B 125, 12718–12729 (2021).

36. Curcio, V., Alemán-Castañeda, L. A., Brown, T. G., Brasselet, S. & Alonso, M. A. Birefringent Fourier filtering for single molecule coordinate and height super-resolution imaging with dithering and orientation. Nature Communications 11, 5307 (2020).

37. Zhang, O. et al. Six-dimensional single-molecule imaging with isotropic resolution using a multi-view reflector microscope. Nature Photonics. ISSN: 1749-4893 (Dec. 2022).

38. Gould, T. J. et al. Nanoscale imaging of molecular positions and anisotropies. Nature Methods 5(12), 1027–1030 (2008).

39. Cruz, C. A. V. et al. Quantitative nanoscale imaging of orientational order in biological filaments by polarized superresolution microscopy. PNAS 7(13544), E820–E828 (2016).

40. Lewis, J. & Lu, Z. Resolution of ångström-scale protein conformational changes by analyzing fluorescence anisotropy. Nature structural and molecular biology 26, 802–807 (2019).

41. Mehta, S. et al. Dissection of molecular assembly dynamics by tracking orientation and position of single molecules in live cells. PNAS, E6352–E6361 (2016).

42. Ohmachi M, e. a. Fluorescence microscopy for simultaneous observation of 3D orientation and movement and its application to quantum rod-tagged myosin V. PNAS 109(14), 5294–5298 (2012).

43. Chung, I., Shimizu, K. & Bawendi, M. Room temperature measurements of the 3D orientation of single CdSe quantum dots using polarization microscopy. PNAS 100(2), 405–408 (2003).

44. Rimoli, C. V., Valades-Cruz, C. A., Curcio, V., Mavrakis, M. & Brasselet, S. 4polar-STORM polarized super-resolution imaging of actin filament organization in cells. Nature Communications 13, 301 (Jan. 2022).

45. Beckwith, J. S. & Yang, H. Sub-millisecond Translational and Orientational Dynamics of a Freely Moving Single Nanoprobe. The Journal of Physical Chemistry B 125, 13436–13443 (Dec. 2021).

46. Monaghan, J. W. et al. Calcite-Assisted Localization and Kinetics (CLocK) Microscopy. The Journal of Physical Chemistry Letters 13. Publisher: American Chemical Society, 10527–10533 (Nov. 2022).

47. Foreman, M. R. & Török, P. Fundamental limits in single-molecule orientation measurements. en. New Journal of Physics 13, 093013. ISSN: 1367-2630 (Sept. 2011).

48. Lewis, J. H., Beausang, J. F., Sweeney, H. L. & Goldman, Y. E. The azimuthal path of myosin V and its dependence on lever-arm length. Journal of General Physiology 139, 101–120. ISSN: 0022-1295. 10.1085/jgp.201110715 (2023) (Jan. 2012).

49. Beausang, J. F., Shroder, D. Y., Nelson, P. C. & Goldman, Y. E. Tilting and Wobble of Myosin V by High-Speed Single-Molecule Polarized Fluorescence Microscopy. English. Biophysical Journal 104. Publisher: Elsevier, 1263–1273. ISSN: 0006-3495. https://www.cell.com/biophysj/abstract/S0006-3495(13)00199-9 (2023) (Mar. 2013).

50. Chun, C. S. L., Fleming, D. L. & Torok, E. J. Polarization-sensitive thermal imaging. Proc. SPIE, Automatic Object Recognition IV 2234, 275–286 (1994).

51. Gruev, V., Perkins, R. & York, T. CCD polarization imaging sensor with aluminum nanowire optical filters. Optics Express 18(18), 19087–19094 (2010).

52. Gruev, V., Van der Spiegel, J. & Engheta, N. Dual-tier thin film polymer polarization imaging sensor. Optics Express 18(18), 19292–19303 (2010).

53. Kulkarni, M. & Gruev, V. Integrated spectral-polarization imaging sensor with aluminum nanowire polarization filters. Optics Express 20(21), 22997–23012 (2012).

54. Ratliff, B., LaCasse, C. & Tyo, J. Interpolation strategies for reducing IFOV artifacts in microgrid polarimeter imagery. Optics Express 17(11) (2009).

55. Tyo, J., LaCasse, C. & Ratliff, B. Total elimination of sampling errors in polarization imagery obtained with integrated microgrid polarimeters. Optics Letters 34(20) (2009).

56. Gao, S. & Gruev, V. Bilinear and bicubic interpolation methods for division of focal plane polarimeters. Opt. Express 19, 26161–26173 (2011).

57. Edelstein, A., Amodaj, N., Hoover, K., Vale, R. & Stuurman, N. Computer Control of Microscopes Using μManager. Current Protocols in Molecular Biology 92, 14.20.1–14.20.17. eprint: https://currentprotocols.onlinelibrary.wiley.com/doi/pdf/10.1002/0471142727.mb1420s92. https://currentprotocols.onlinelibrary.wiley.com/doi/abs/10.1002/0471142727.mb1420s92 (2010).

58. Sofroniew, N. et al. napari: a multi-dimensional image viewer for Python Nov. 2022. https://zenodo.org/record/7276432 (2023).

59. Axelrod, D. Carbocyanine dye orientation in red cell membrane studied by microscopic fluorescence polarization. Biophys J 26, 557–574 (1979).

60. Fourkas, J. T. Rapid determination of the three-dimensional orientation of single molecules. Opt. Lett. 26, 211–213. http://opg.optica.org/ol/abstract.cfm?URI=ol-26-4-211 (2001).

61. Ströhl, F., Bruggeman, E., Rowlands, C. J., Wolfson, D. L. & Ahluwalia, B. S. Quantification of the NA dependent change of shape in the image formation of a z-polarized fluorescent molecule using vectorial diffraction simulations. Microscopy Research and Technique 85, 2016–2022. https://onlinelibrary.wiley.com/doi/abs/10.1002/jemt.24060 (x2022) (2022).

62. Born, M. & Wolf, E. Principles of Optics 7th (Cambridge University Press, 2019).

63. Gunturk, B., Glotzbach, J., Altunbasak, Y., Schafer, R. & Mersereau, R. Demosaicking: color filter array interpolation. IEEE Signal Processing Magazine 22. Conference Name: IEEE Signal Processing Magazine, 44–54. ISSN: 1558-0792 (Jan. 2005).

64. Li, N., Zhao, Y., Pan, Q. & Kong, S. G. Removal of reflections in LWIR image with polarization characteristics. Opt. Express 26, 16488–16504. https://opg.optica.org/oe/abstract.cfm?URI=oe-26-3-16488 (2018).

65. Spehar, K. et al. Super-resolution imaging of amyloid structures over extended times by using transient binding of single Thioflavin T molecules. Chem. Bio. Chem. 19, 1944–1948 (2018).

66. Nieuwenhuizen, R. P. J. et al. Measuring image resolution in optical nanoscopy. Nature Methods 10, 557–562 (June 2013).

67. Mazidi, H., King, E. S., Zhang, O., Nehorai, A. & Lew, M. D. Dense super-resolution imaging of molecular orientation via joint sparse basis deconvolution and spatial pooling. IEEE 16th International Symposium on Biomedical Imaging (ISBI 2019), 325–329 (2019).

68. Danylchuk, D. I., Moon, S., Xu, K. & Klymchenko, A. S. Switchable Solvatochromic Probes for Live-Cell Super-resolution Imaging of Plasma Membrane Organization. Angewandte Chemie International Edition 58, 14920–14924 (2019).

69. Göhring, J., Schrangl, L., Schütz, G. J. & Huppa, J. B. Mechanosurveillance: Tiptoeing T Cells. Frontiers in Immunology 13. ISSN: 1664-3224. https://www.frontiersin.org/articles/10.3389/fimmu.2022.886328 (x2023) (2022).

70. Mahecic, D. et al. Event-driven acquisition for content-enriched microscopy. Nature Methods 19, 1262–1267 (Oct. 2022).

71. Alvelid, J., Damenti, M., Sgattoni, C. & Testa, I. Event-triggered STED imaging. Nature Methods 19, 1268–1275 (Oct. 2022).

72. Liberatore, A. et al. PolarCam micropolarizer cameras characterization and usage in International Conference on Space Optics — ICSO 2020 11852 (SPIE, June 2021), 358–379. https://www.spiedigitallibrary.org/conference-proceedings-of-spie/11852/118520W/PolarCam-micropolarizer-cameras-characterization-and-usage/10.1117/12.2599180.full (2023).

73. Sugizaki, A. et al. POLArIS, a versatile probe for molecular orientation, revealed actin filaments associated with microtubule asters in early embryos. PNAS 118(11), e2019071118 (2021).

74. Shuhei Shibata, S., Takano, W., Hagen, N., Matsuda, M. & Otani, Y. Video-rate quantitative phase analysis by a DIC microscope using a polarization camera. Biomedical Optics Express 10(3), 1273– 1281 (2019).

75. Yeh, L.-H. et al. uPTI: uniaxial permittivity tensor imaging of intrinsic density and anisotropy in Biophotonics Congress 2021 (2021), paper NM3C.4 (Optica Publishing Group, Apr. 2021), NM3C.4. https://opg.optica.org/abstract.cfm?uri=NTM-2021-NM3C.4 (2023).

76. Kalita, R. et al. Single-shot phase contrast microscopy using polarisation-resolved differential phase contrast. Journal of Biophotonics 14, e202100144 (2021).

77. Song, S., Kim, J., Hur, S., Song, J. & Joo, C. Large-Area, High-Resolution Birefringence Imaging with Polarization-Sensitive Fourier Ptychographic Microscopy. ACS Photonics 8, 158–165 (Jan. 2021).

## Methods-only references

1. Backer, A. & Moerner, W. Extending single-molecule microscopy using optical fourier processing. J Phys Chem B 118, 8313–8329 (2014).

2. Hecht, E. Optics 4th (Addison Wesley, 2002).

3. Edelstein, A., Amodaj, N., Hoover, K., Vale, R. & Stuurman, N. Computer Control of Microscopes Using μManager. Current Protocols in Molecular Biology 92, 14.20.1–14.20.17. eprint: https://currentprotocols.onlinelibrary.wiley.com/doi/pdf/10.1002/0471142727.mb1420s92. https://currentprotocols.onlinelibrary.wiley.com/doi /abs/10.1002/0471142727.mb1420s92 (2010).

4. Sofroniew, N. et al. napari: a multi-dimensional image viewer for Python Nov. 2022. https://zenodo.org/record/7276432 (2023).

5. Bruggeman, E. Imaging single SYTOX Orange molecules on a PLL-coated cover glass. en. https://www.protocols.io/view/imaging-single-sytox-orange-molecules-on-a-pll-coa-c672zhqe (2024) (May 2024).

6. Wu, T., Lu, J. & Lew, M. D. Dipole-spread-function engineering for simultaneously measuring the 3D orientations and 3D positions of fluorescent molecules. Optica 9, 505–511 (2022).

7. Bruggeman, E. Preparation and imaging of lipid bilayer-coated silica microspheres. en. https://www.protocols.io/view/preparation-and-imaging-of-lipid-bilayer-coated-si-ddca22se (2024) (May 2024).

8. Bruggeman, E. Imaging single AF647 molecules immobilised in PVA on a cover glass. en. https://www.protocols.io/view/imaging-single-af647-molecules-immobilised-in-pva-c68tzhwn (2024) (May 2024).

9. Bruggeman, E. TAB-PAINT imaging of alpha-synuclein fibrils using Nile Red. en. https://www.protocols.io/view/tab-paint-imaging-of-alpha-synuclein-fibrils-using-c69bzh2n (2024) (May 2024).

10. Bruggeman, E. & Peters, R. dSTORM of actin in fixed HeLa cells. en. https://www.protocols.io/view/dstorm-of-actin-in-fixed-hela-cells-ddet23en (2024) (May 2024).

11. Bruggeman, E. & Körbel, M. Live-cell imaging of the plasma membrane of Jurkat T cells. en. https://www.protocols.io/view/live-cell-imaging-of-the-plasma-membrane-of-jurkat-ddeu23ew (2024) (May 2024).

