## Supplementary Information for "POLCAM: Instant molecular orientation microscopy for the life sciences"

### List of supplementary datasets

- **Data 1 SYTOX orange:** Representative unprocessed data set of SYTOX orange dispersed on a cover glass.
- **Data 2 PAINT a-syn fibrils:** Representative unprocessed data of TAB-PAINT data of Nile red reversibly binding to alpha-synuclein fibrils attached to a cover glass.
- **Data 3 dSTORM phalloidin-AF488:** Representative unprocessed dSTORM data of fixed HeLa cells labeled with phalloidin-AF488, showing highly linearly polarized emission (*i.e.*, rotationally restricted fluorophores).
- **Data 4 dSTORM phalloidin-AF647:** Representative unprocessed dSTORM data of fixed HeLa cells labeled with phalloidin-AF647, showing less linearly polarized emission (*i.e.*, less rotationally restricted fluorophores).

### List of supplementary videos

- **Video S1 SYTOX Orange:** Supporting movie showing single SYTOX Orange molecules dispersed on a cover glass. The movie shows raw image data from the polarization camera and highlights four example molecules at different orientations and rotational mobilities. The 4 example molecules are shown in unprocessed image format, $S_0$, and an HSV polarization colormap.
- **Video S2 TAB-PAINT fibrils:** Supporting movie showing data from a TAB-PAINT experiment, showing single Nile red molecules binding along the long axis of amyloid fibrils composed of the protein alpha-synuclein. The data is shown in an unprocessed format before and after background correction, an HSV polarization colormap, and finally, localizations are built up to generate a multi-dimensional super-resolution image (from 10 seconds worth of image data).
- **Video S3 dSTORM phalloidin-AF488:** Supporting movie showing data from a dSTORM experiment on fixed HeLa cells labeled with phalloidin-AF488. The reconstructed super-resolution image is shown on the left-hand side. A small region of interest is highlighted, for which the unprocessed dSTORM data is shown on the right-hand side.
- **Video S4 dSTORM phalloidin-AF647:** Supporting movie showing data from a dSTORM experiment on fixed HeLa cells labeled with phalloidin-AF647. Unprocessed image data is shown, with a magnified inset. The inset is also shown in the processed $S_0$ format, which is used to determine the position of single molecules during localization.
- **Video S5 COS-7 SiR-actin:** Supporting movie demonstrating diffraction-limited polarization camera imaging of fixed COS-7 cells labeled with SiR-actin in an HSV polarization colormap. During the acquisition, the stage was moved to explore different regions of the sample.
- **Video S6 Live T cell NR4A 3D:** Supporting movie showing a diffraction-limited 3D polarization camera image of the membrane (stained with NR4A) of a live human T cell. The volume was acquired by scanning the objective and is rendered in an HSV colormap. The polarized cell body is readily differentiated from the filopodia, which appear white.
- **Video S7 Live T cell NR4A 4D:** Supporting movie showing a diffraction-limited polarization camera 3D movie of the membrane (stained with NR4A) of a live human T cell. The filopodia can be seen interacting with the antibody-coated cover glass. The recording is replayed four times during the video.

### S1 Optical model

The emission of a fluorescent molecule was modeled as the far-field of an oscillating electric dipole as previously described in [1]. For completeness, we will summarize the model here. The detection by the objective lens is modeled using vectorial diffraction theory. Transmission through the micropolarizer array on the camera chip is modeled using Jones matrices for a linear polarizer on the calculated electric field at the image plane.

#### S1.1 Field emitted by a fluorescent molecule

We consider a fluorescent molecule at position $\mathbf{r}'$ in object space with an emission dipole moment $\vec{\mu}$

$$\vec{\mu} \propto \begin{pmatrix} \mu_x \\ \mu_y \\ \mu_z \end{pmatrix} = \begin{pmatrix} \sin\Theta\cos\Phi \\ \sin\Theta\sin\Phi \\ \cos\Theta \end{pmatrix} \tag{8}$$

where $\Theta$ is the polar angle and $\Phi$ the azimuthal angle we wish to determine in our measurements. If we treat the molecule as an oscillating electric dipole, the emitted electric field at a point $\mathbf{r}$ in object space will be [2]

$$\mathbf{E}(\mathbf{r}) \propto \mathbf{G}(\mathbf{r}, \mathbf{r}')\hat{\mu} \tag{9}$$

where $\mathbf{G}$ is the dyadic Green's function. In the far-field (i.e. for fields that appear a distance $r \gg \lambda$ from the molecule) the dyadic Green's function is [2]

$$\mathbf{G}_{\text{ff}}(\mathbf{r}, \mathbf{r}') \simeq \left(\mathbf{I} - \hat{r}\hat{r}\right) \frac{e^{ink_0\|\mathbf{r}-\mathbf{r}'\|}}{4\pi\|\mathbf{r}-\mathbf{r}'\|} \tag{10}$$

where $\mathbf{I}$ is an identity matrix, $\hat{r} = \mathbf{r}/\|\mathbf{r}\|$ is a unit vector, $k_0 = 2\pi/\lambda$ the wavenumber in free space and $\lambda$ the wavelength, and $n$ is the refractive index of the sample medium. Using spherical coordinates $\hat{r} = (\sin\theta\cos\phi, \sin\theta\sin\phi, \cos\theta)$ and placing the molecule in the origin $\mathbf{r}' = (0, 0, 0)$ of object space we get

$$\mathbf{G}_{\text{ff}}(\phi, \theta, r) \simeq \begin{pmatrix} 1 - \sin^2\theta\cos^2\phi & -\sin^2\theta\sin\phi\cos\phi & -\sin\theta\cos\theta\cos\phi \\ -\sin^2\theta\cos\phi\sin\phi & 1 - \sin^2\theta\sin^2\phi & -\sin\theta\cos\theta\sin\phi \\ -\sin\theta\cos\theta\cos\phi & -\cos\theta\sin\theta\sin\phi & 1 - \cos^2\theta \end{pmatrix} \frac{e^{ink_0 r}}{4\pi r} \tag{11}$$

where $r = \|\mathbf{r}\|$. We can later place the molecule at different positions in object space by adding a tip, tilt and defocus phase term to the field at the back focal plane. Considering the small size of our fields of view and assuming spatial shift invariance, this is reasonable. The Green's function can be modified for a dipole emitter near a planar refractive index interface [1], *e.g.* the interface between the cover glass and an aqueous sample medium when imaging using an oil-immersion objective.

#### S1.2 Modelling refraction by the objective lens

The objective lens will capture a cone with half angle $\theta_{max} = \sin^{-1}(\text{NA}/n)$ of the emitted field and refract it. This can be modeled using a rotation matrix $\mathbf{R}_{\text{obj}}$ which is a combination of a rotation matrix $\mathbf{R}$ and refraction matrix $\mathbf{L}$ [3]

$$\mathbf{R}_{\text{obj}} = \mathbf{R}^{-1}\mathbf{L}\mathbf{R}$$

$$\mathbf{R} = \begin{pmatrix} \cos\phi & \sin\phi & 0 \\ -\sin\phi & \cos\phi & 0 \\ 0 & 0 & 1 \end{pmatrix} \quad ; \quad \mathbf{L} = \frac{1}{A(\theta)} \begin{pmatrix} \cos\theta & 0 & \sin\theta \\ 0 & 1 & 0 \\ -\sin\theta & 0 & \cos\theta \end{pmatrix}$$

where $A(\theta) = \sqrt{(n_1/n_2)|\cos\theta|}$ is an apodization factor for aplanatic collimation and $n_1$ and $n_2$ the refractive indices before and after the lens. We neglect the apodization factor for simplicity. The electric field in the back focal plane of the objective (at $r = f_{\text{obj}}$) is then

$$\begin{aligned} \mathbf{E}_{\text{bfp}}(\phi, \theta; \boldsymbol{\mu}) &= \mathbf{R}_{\text{obj}}\mathbf{G}_{\text{ff}}(\phi, \theta)\hat{\mu} \\ &= \frac{e^{ink_0 f_{\text{obj}}}}{4\pi f_{\text{obj}}} \begin{bmatrix} \sin^2\phi + \cos^2\phi\cos\theta & \sin(2\phi)(\cos\theta - 1)/2 & -\sin\theta\cos\phi \\ \sin(2\phi)(\cos\theta - 1)/2 & \cos^2\phi + \sin^2\phi\cos\theta & -\sin\theta\sin\phi \\ 0 & 0 & 0 \end{bmatrix} \begin{bmatrix} \mu_x \\ \mu_y \\ \mu_z \end{bmatrix} \\ &= \frac{e^{ink_0 f_{\text{obj}}}}{4\pi f_{\text{obj}}} \begin{bmatrix} G_{xx} & G_{xy} & G_{xz} \\ G_{yx} & G_{yy} & G_{yz} \\ G_{zx} & G_{zy} & G_{zz} \end{bmatrix} \begin{bmatrix} \mu_x \\ \mu_y \\ \mu_z \end{bmatrix} \end{aligned} \tag{12}$$

#### S1.3 Positioning the molecule in object space

To place the molecule at a position $\mathbf{r}' = (x, y, z)$ in object space, we add a phase term to the field in the back focal plane

$$\mathbf{E}_{\text{bfp}}(\phi, \rho; x, y, z) = \mathbf{E}_{\text{bfp}}(\phi, \rho)e^{i\Psi(x,y,z)} \tag{13}$$

$$\Psi(x, y, z) = nk_0(x\rho\cos\phi + y\rho\sin\phi + z\sqrt{1-\rho^2}) \tag{14}$$

#### S1.4 Modeling refraction by the tube lens

The tube lenses used in widefield microscopy have a low NA, so the field at the image plane can be accurately calculated using a scaled Fourier transform of the field in the back focal plane. The pixel size in the image plane $d_{\text{img}}$ is then $\lambda f_{\text{tube}}/\o$, where $\lambda$ is the wavelength, $f_{\text{tube}}$ the focal length of the tube lens, $\o$ the diameter of the field being transformed. The pixel size in simulations was adjusted by padding the field in the back focal plane with zeros before taking a Fourier transform, i.e. $\o = (\o_{\text{bfp}} + \text{padding})$. The diameter of the back focal plane is $\o_{\text{bfp}} = 2f_{\text{obj}}\text{NA}$. For simplicity, we neglected any effects that the dichroic might have on the field.

#### S1.5 Transmission of the field through the micropolarizer array

The transmission of the emission through the micropolarizer array on the camera sensor was modeled for each camera pixel using a Jones matrix $J_{LP}$ for a linear polarizer with an axis of transmission at an angle $\eta$ from the x-axis [4]

$$\mathbf{E}' = J_{LP}\mathbf{E} \tag{15}$$

$$J_{LP} = \begin{pmatrix} \cos^2\eta & \cos\eta\sin\eta \\ \cos\eta\sin\eta & \sin^2\eta \end{pmatrix} \tag{16}$$

Considering the low NA of the tube lens and the fact that the polarizer microarray is deposited directly onto the chip, we applied the Jones matrices to the field in the image plane. The z-component of the field has been omitted because its contribution to the final intensity distribution is negligible. The intensity at the image plane is calculated from the electric field as $I = |E'_x|^2 + |E'_y|^2$. Figure **??** shows a simulated image of immobilized fluorescent molecules at different orientations spaced in a grid.

#### S1.6 Rotational mobility

Simulations that include rotational mobility assuming a uniform rotation in a cone model were implemented using weighted combinations of basis functions [1]

$$I_{\text{img}} \propto \begin{pmatrix} \mu_x^2\gamma + \frac{1-\gamma}{3} \\ \mu_y^2\gamma + \frac{1-\gamma}{3} \\ \mu_z^2\gamma + \frac{1-\gamma}{3} \\ \mu_x\mu_y\gamma \\ \mu_x\mu_z\gamma \\ \mu_y\mu_z\gamma \end{pmatrix}^T \begin{pmatrix} XX^x + XX^y \\ YY^x + YY^y \\ ZZ^x + ZZ^y \\ XY^x + XY^y \\ XZ^x + XZ^y \\ ZY^x + ZY^y \end{pmatrix} \tag{17}$$

where the basis functions $\{\text{XX}^{x,y}, \text{YY}^{x,y}, \text{ZZ}^{x,y}, \text{XY}^{x,y}, \text{XZ}^{x,y}, \text{YZ}^{x,y}\}$ are defined as in [1], and $\gamma \in [0, 1]$ is a rotational mobility parameter [5] that relates to the size of the cone in which an emitter is free to rotate. In the case of complete rotational freedom $\gamma = 0$, and for complete rotational immobilization, $\gamma = 1$.

#### S1.7 Detection noise model

The following detection model for a CMOS camera was used

$$\begin{aligned} n_{e^-} &= \mathcal{P}\{\text{QE} \cdot I(x, y)\} + n_{\text{read}} \\ n_{\text{ADU}} &= \text{floor}(n_{e^-}/e_{\text{ADU}}) + o \end{aligned} \tag{18}$$

where $I(x, y)$ the detected signal in photons, QE is the quantum efficiency of the detector at the average wavelength, $\mathcal{P}$ refers to a Poisson distribution, $n_{\text{read}} \sim \mathcal{N}(0, \sigma_{\text{RMS}})$ Gaussian read noise with variance $\sigma^2_{\text{RMS}}$ and $o$ the camera offset in ADU (analog-to-digital) counts. The final camera image is $n_{\text{ADU}}$ after taking into account camera saturation. CMOS cameras have a readout circuitry for each individual pixel causing the offset, gain, and variance to vary between pixels, but for simplicity, we used the same parameters for all pixels in simulations. The polarization camera noise parameters used in simulations are summarized in Table 1.

| | CS505MUP (Thorlabs) | hypothetical state-of-the-art | ideal camera |
|---|---|---|---|
| QE (e-/photon) | 0.70 | 0.95 | 1 |
| $e_{ADU}$ (ADU/e-) | 0.424 | 1 | 1 |
| RMS read noise (e-) | 2.082 | 1 | 0 |
| offset (ADU) | 100 | 100 | 0 |
| bit | 12 | 16 | - |

**Table 1:** Summary of polarisation camera noise parameters used in simulations. The first column (CS505MUP, Thorlabs) contains values that have been measured using a camera calibration as described in section S1.8.

#### S1.8 Camera calibration

The camera that was used in this work was calibrated using the method described in [6]. The code used is available on GitHub at `https://github.com/TheLeeLab/cameraCalibrationCMOS`. The result was validated on a smaller calibration dataset using the camera calibration (Mean-Variance Test) included in the ImageJ [7] plugin GDSC SMLM [8]. Full sensor images (2448 x 2048 pixels) were recorded at 7 different intensities (including dark frames), with 20.000 frames per intensity level. The measured pixel-dependent offset, gain and read noise are shown in figure S1 and summarized in Table 2. The results of the full-sensor camera calibration (the tif files of the pixel-dependent offset, gain, variance and readnoise) were uploaded to Zenodo at `https://zenodo.org/doi/10.5281/zenodo.10578307`.

| | cameraCalibrationCMOS | GDSC SMLM |
|---|---|---|
| Offset (ADU) | 100.4 $\pm$ 0.3 | 100.4 $\pm$ 0.897 |
| Gain (ADU/e-) | 0.424 $\pm$ 0.004 | 0.424 |
| Read noise (e-) | 2.1 $\pm$ 0.3 | 2.115 |
| Read noise RMS (e-) | 2.082 | - |
| Read noise median (e-) | 2.133 | - |

**Table 2:** Validation of the results of the CMOS camera calibration code used in this work (`https://github.com/TheLeeLab/cameraCalibrationCMOS`) through comparison with results from the ImageJ [7] plugin GDSC SMLM [8].

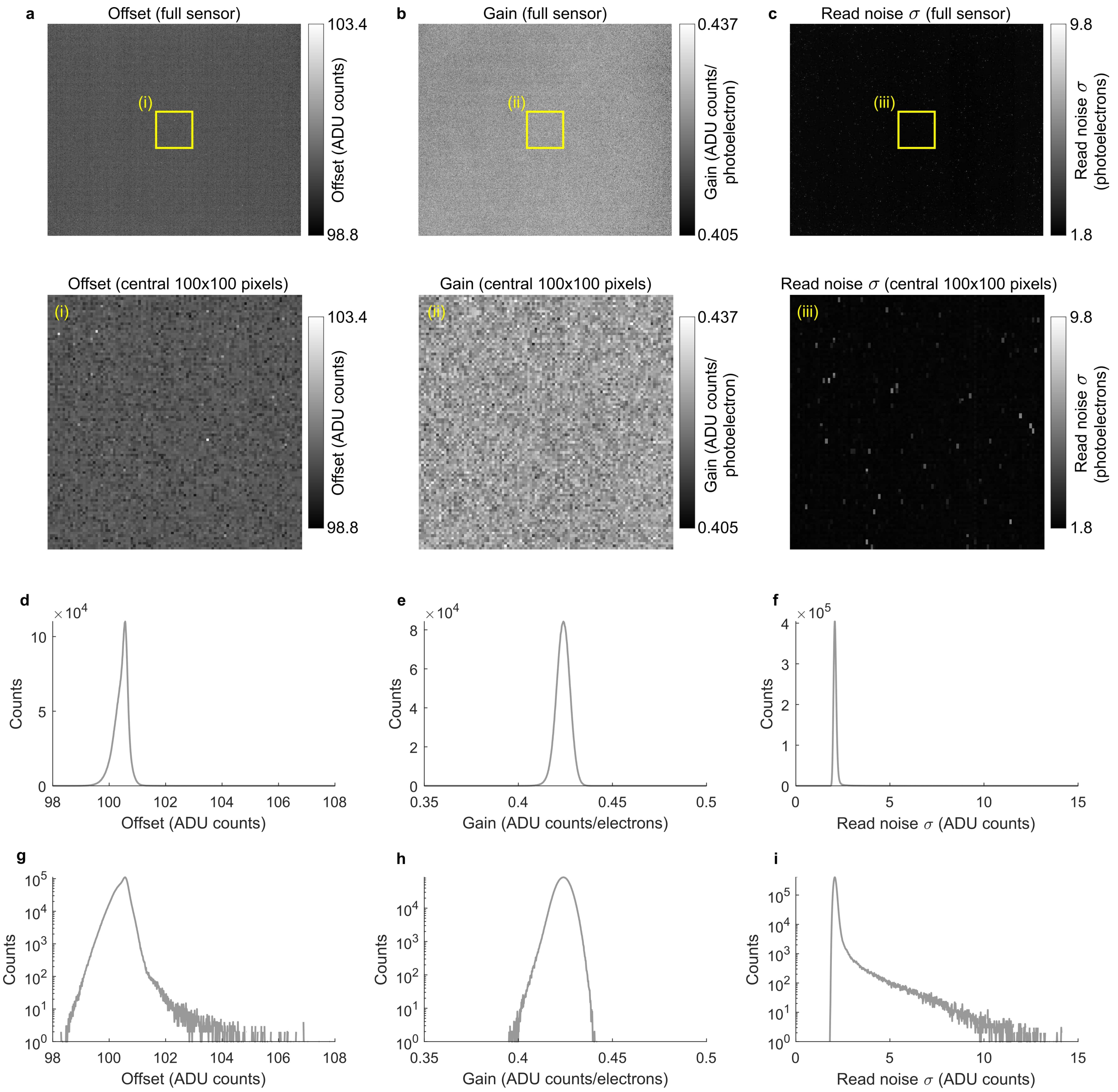

**Figure S1: Measured pixel-dependent gain, offset, and read noise. a** Top, pixel-dependent measured offset for the full camera sensor (2448 x 2048 pixels). Bottom, the 100 x 100 central pixels off the offset map (as marked by the yellow box). **b)** The same as panel a for the pixel-dependent gain. **c)** The same as panels a and b, but for the pixel-dependent read noise standard deviation. **d** Histogram of the offset values from the whole sensor in panel a. **e)** Histogram of the gain values from the whole sensor in panel b. **f)** Histogram of the read noise standard deviation values from the whole sensor in panel c. **g-i** The same histograms as shown in panels d,e and f but with a logarithmic y-axis to show outliers.

#### S2 Comparison with a state-of-the-art sCMOS camera

To compare the performance of the polarization camera with a state-of-the-art sCMOS camera, we imaged single fluorescent beads and molecules simultaneously on the polarization camera and the Prime 95B from Teledyne Photometrics using a 50:50 non-polarizing beam splitter setup as depicted in Fig. S2a. The relay lenses on the setup (L1, L2, and L3) were chosen such that the virtual pixel size of the Prime 95B camera was 110 nm, and 69 nm for the polarization camera.

Fig. S2b shows that we can register the images from both cameras onto each other using a similarity transform (only translation, rotation, and global scaling). Before registration, the polarization camera image was computationally resized (using the MATLAB function *imresize* and bilinear interpolation) to have a pixel size of 110 nm, so both images have the same magnification. The image registration was performed using the Registration Estimator app in MATLAB, using a multimodal intensity-based registration (similarity transform with 1.000 iterations).

Fig. S2c shows scatter plots of the estimated position of fluorescent beads (TetraSpeck™ Microspheres, 0.1 µm, fluorescent blue/green/orange/dark red, T7279, Invitrogen) on both cameras. Image data similar to the data shown in Fig. S2b was taken, for 100 frames. The same beads were identified in the images from both cameras and localized in all 100 frames. The localizations of some representative beads are shown as scatter plots. Above each scatter plot, the mean number of photons detected per frame and the standard deviation on the localization coordinates are shown. The six examples shown are for the same 6 beads on each camera. As expected, the polarization camera detects about 50% of the photons compared to the Prime 95B, as the polarizers absorb light. Secondly, the localization precision gets better (standard deviation smaller) when more photons are detected. As a result, the localization precision on the sCMOS camera is better than on the polarization camera, but not dramatically. This might be because at high photon numbers, having a smaller pixel size improves the localization precision.

Fig. S2d shows that the polarization camera detects half as many photons as the Prime 95B. Each dot on the graph represents the average number of photons detected per frame from one single bead, on both cameras. The average is calculated from 100 repeats. Photon numbers are calculated by taking into account the camera gain and wavelength-dependent quantum efficiency. The gray line is the function $y(x) = x/2$. We do indeed expect the data to follow this trend, as the emission of beads will be approximately randomly polarized (a bead is a collection of randomly oriented dyes), and according to Malus' Law, randomly polarized light will be attenuated by 50% after passing through a linear polarizer.

Fig. S2e shows that we can detect a single molecule on both cameras simultaneously. Single Cy5 molecules embedded and immobilized in polymer (PMMA) were imaged with an exposure time of 200 ms. The example shown is in the top 5% brightest emitters in the collected dataset. The number of detected photons is correlated between the two cameras; we observe one blinking event, followed by permanent photobleaching in a single step, characteristic of single fluorescent molecules:

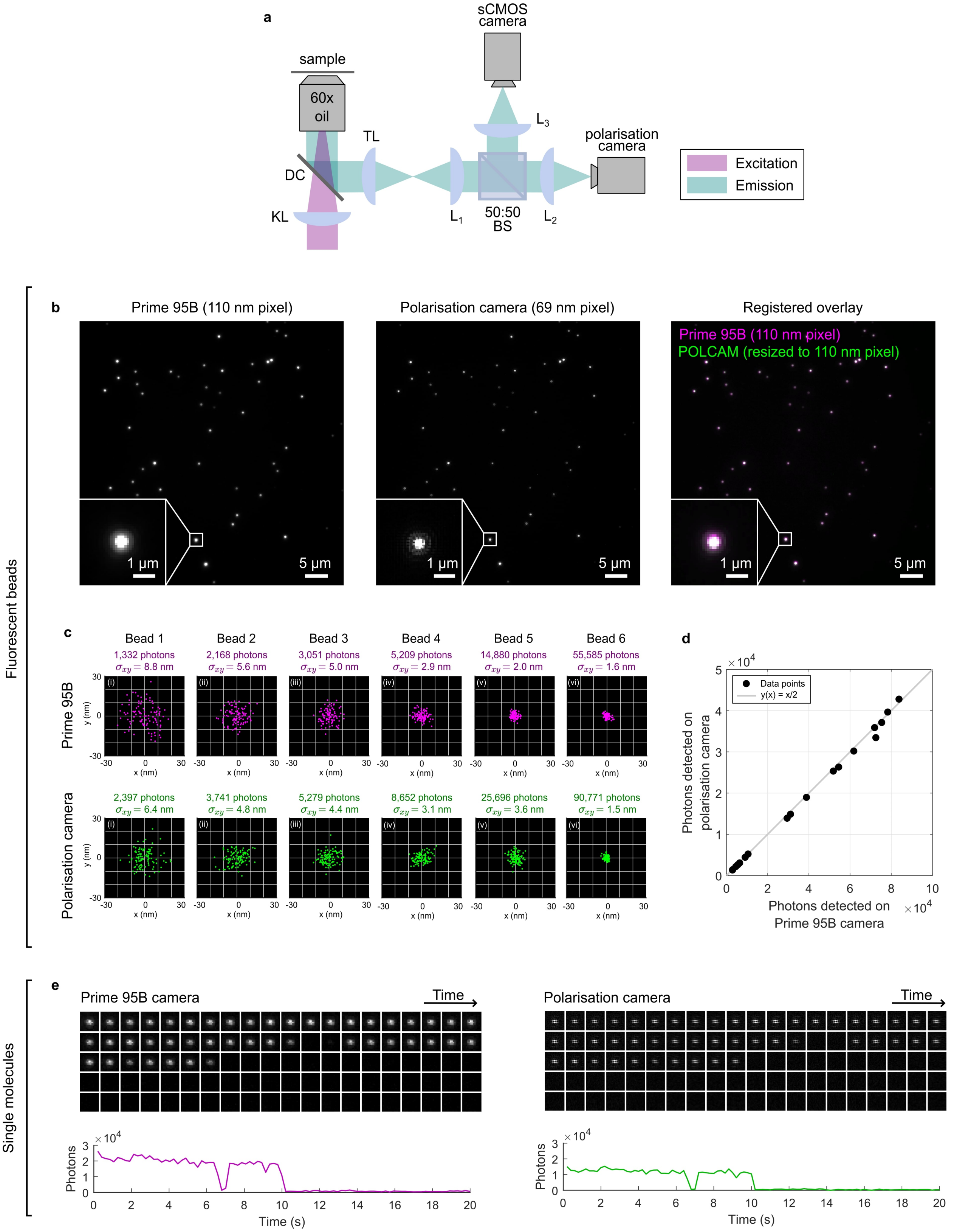

**Figure S2: Comparison with a state-of-the-art sCMOS camera.** **a)** Diagram of the optical setup. **b)** Image of 100 nm diameter fluorescent beads recorded simultaneously on the Prime 95B and the polarization camera. **c)** Localizations of 6 representative beads that were recorded simultaneously on the Prime 95B (top row, in magenta) and the polarization camera (bottom row, in green). **d)** The average intensity of fluorescent beads detected simultaneously on the Prime 95B (x-axis) and the polarization camera (y-axis). **e)** A single Cy5 molecule immobilised in polymer (PMMA) imaged on both cameras simultaneously using a 200 ms exposure time. The montages show the raw image data. The photon traces are the photon values over time, showing clear correlation, a blinking event and single-step permanent photobleaching.

### S3 Example simulations

This section contains simulated single-molecule images, at different 3D orientations, rotational mobilities, and signal-to-noise ratio. The following parameters were used for the simulations presented in this section (unless otherwise specified in the figure caption): 650 nm emission wavelength, 60x oil-immersion objective with a numerical aperture of 1.42, a tube lens with a focal length of 200 mm, physical camera pixel size of 3.45 µm. All molecules are in focus ($z = 0$) and in the presence of a refractive index interface, they are positioned on the interface. The refractive index of the cover glass and immersion medium are assumed to be the same ($n_{\text{glass}} = n_{\text{immersion}} = n_{\text{medium}} = 1.518$), and the refractive index of water if $n_{\text{medium}} = 1.33$. No noise was added and the molecule is perfectly rotationally immobilized ($\gamma = 1$).

#### S3.1 Intensity and polarization at the back focal plane

Figures S3 and S4 show the simulated back focal plane for a range of dipole orientations.

#### S3.2 Intensity and polarization at the image plane

Figure S5 shows the simulated back focal plane intensity distribution and image plane intensity distribution for 4 example orientations, in the absence of a refractive index mismatch. The back focal plane and image plane are shown in gray scale, and in a polarization colormap. Figure S6 shows the same, but for a molecule that is positioned on a planar refractive interface between cover glass (n = 1.518) and an aqueous medium (n = 1.33).

Figure S7 shows the intensity at the image plane for a *regular* camera and a polarization camera. Figures S8 and S9 show the corresponding Stokes parameter images ($S_0$, $S_1$ and $S_2$) and polarization colormap rendering.

#### S3.3 Image plane for varying rotational mobility, $\gamma$

Figures S12 and S13 show simulations of the Stokes parameter images ($S_0$, $S_1$ and $S_2$) at the image plane for different rotational mobility parameters $\gamma$, respectively for (i) different out-of-plane angles $\theta$ ($\phi = 0°$), and (ii) different in-plane angles ($\theta = 90°$).

Figures S14 and S15 show respectively the DoLP colormap and polarization colormap representation of the same rotational mobilities and orientations as shown in figures S12 and S13.

#### S3.4 Image plane for varying signal-to-noise ratios (SNR)

To generate figures S16- S19, noisy polarization camera images were simulated and processed using cubic spline interpolation. To generate the noisy images, the detection noise model described in section S1.7 was used, using the experimentally measured camera calibration parameters in Table 2. A fixed background of 10 photons per pixel was also added to all images.

Figures S16 and S17 show simulations of the Stokes parameter images ($S_0$, $S_1$ and $S_2$) at the image plane for different signal-to-noise ratios, respectively for (i) different out-of-plane angles $\theta$ ($\phi = 0°$), and (ii) different in-plane angles ($\theta = 90°$).

Figures S18 and S19 show respectively the DoLP colormap and polarization colormap representation of the same rotational mobilities and orientations as shown in figures S16 and S17.

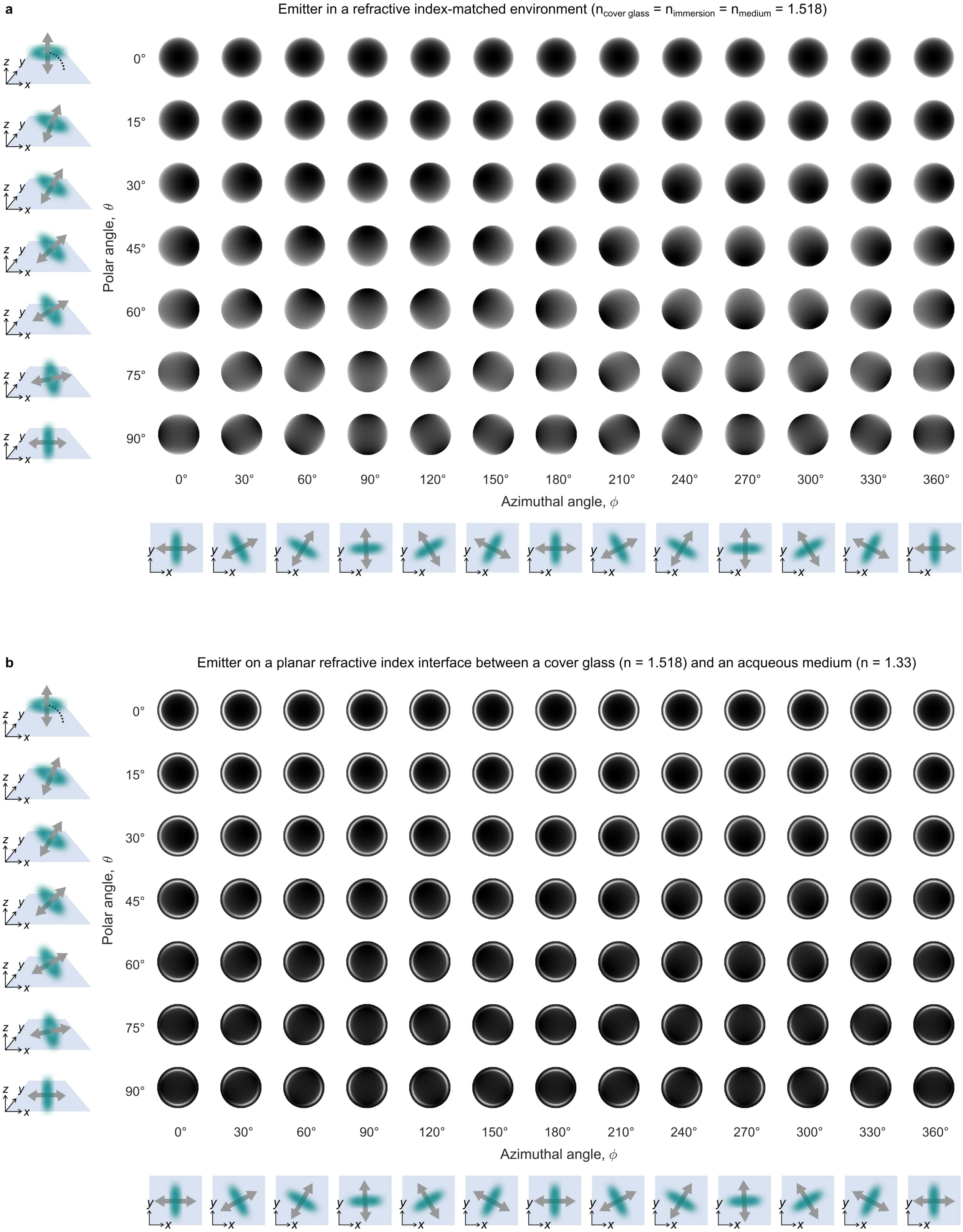

**Figure S3: Simulated back focal plane for different dipole orientations: Intensity.** The simulation parameters used to generate this figure are described in section S3. **a)** These simulations assume there is no refractive index mismatch ($n_{\text{glass}} = n_{\text{immersion}} = n_{\text{medium}} = 1.518$). **b)** These simulations assume there is a planar refractive index mismatch between the cover glass ($n_{\text{glass}} = n_{\text{immersion}} = 1.518$) and the sample medium ($n_{\text{sample}} = 1.33$).

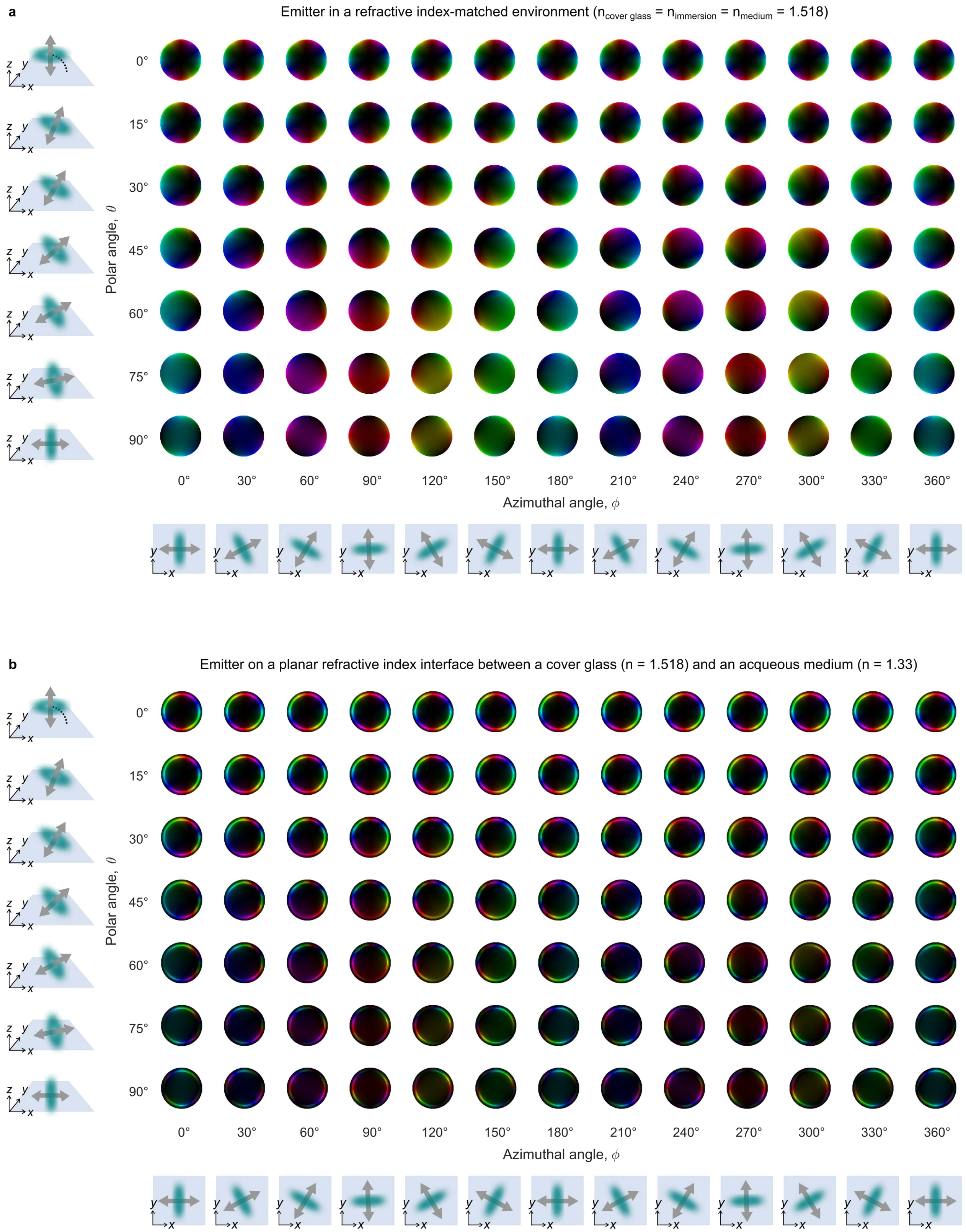

**Figure S4: Simulated back focal plane for different dipole orientations: Polarization.** The simulation parameters used to generate this figure are described in section S3. **a)** These simulations assume there is no refractive index mismatch ($n_{\text{glass}} = n_{\text{immersion}} = n_{\text{medium}} = 1.518$). **b)** These simulations assume there is a planar refractive index mismatch between the cover glass ($n_{\text{glass}} = n_{\text{immersion}} = 1.518$) and the sample medium ($n_{\text{sample}} = 1.33$).

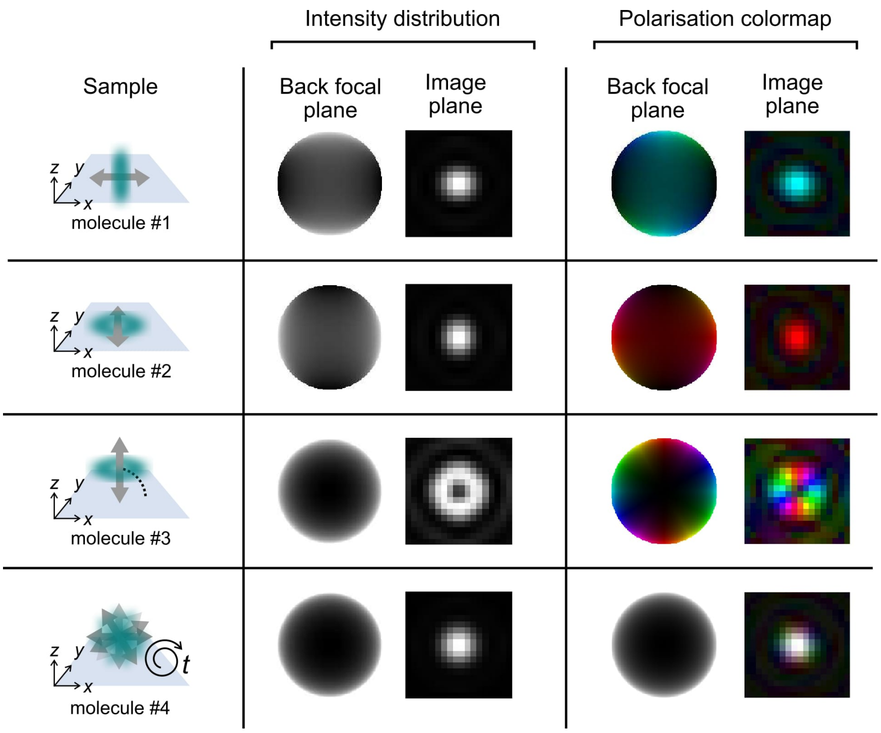

**Figure S5: Simulated back focal plane and image plane of four example emitters: no RI mismatch** The simulation parameters used to generate this figure are described in section S3.

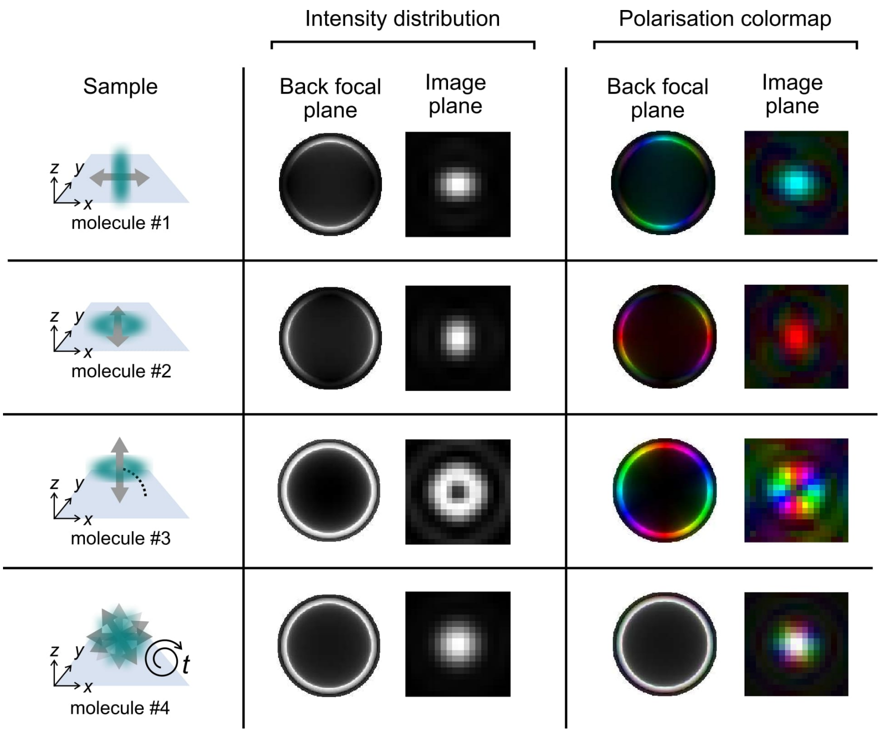

**Figure S6: Simulated images of immobilized single molecules: RI mismatch** The simulation parameters used to generate this figure are described in section S3.

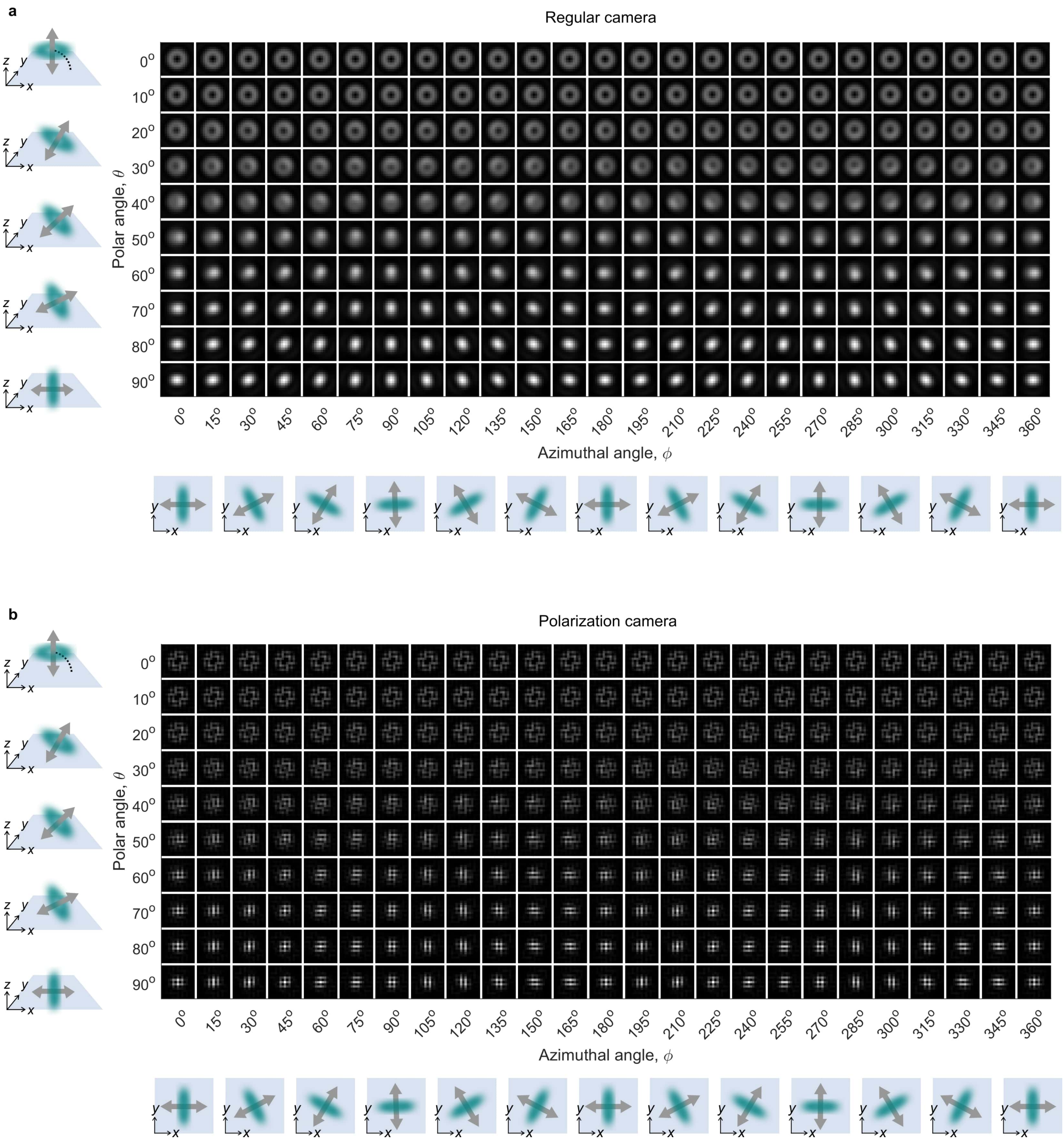

**Figure S7: Simulated image plane for different dipole orientations: Intensity. a)** Simulated images of an immobilized single fluorescent molecule at different 3D orientations, assuming a regular camera. **b)** The same images as shown in panel a, but for a polarization camera. The simulation parameters used to generate this figure are described in section S3. The molecules are positioned on a planar refractive index interface between glass (n=1.158) and water (n=1.33).

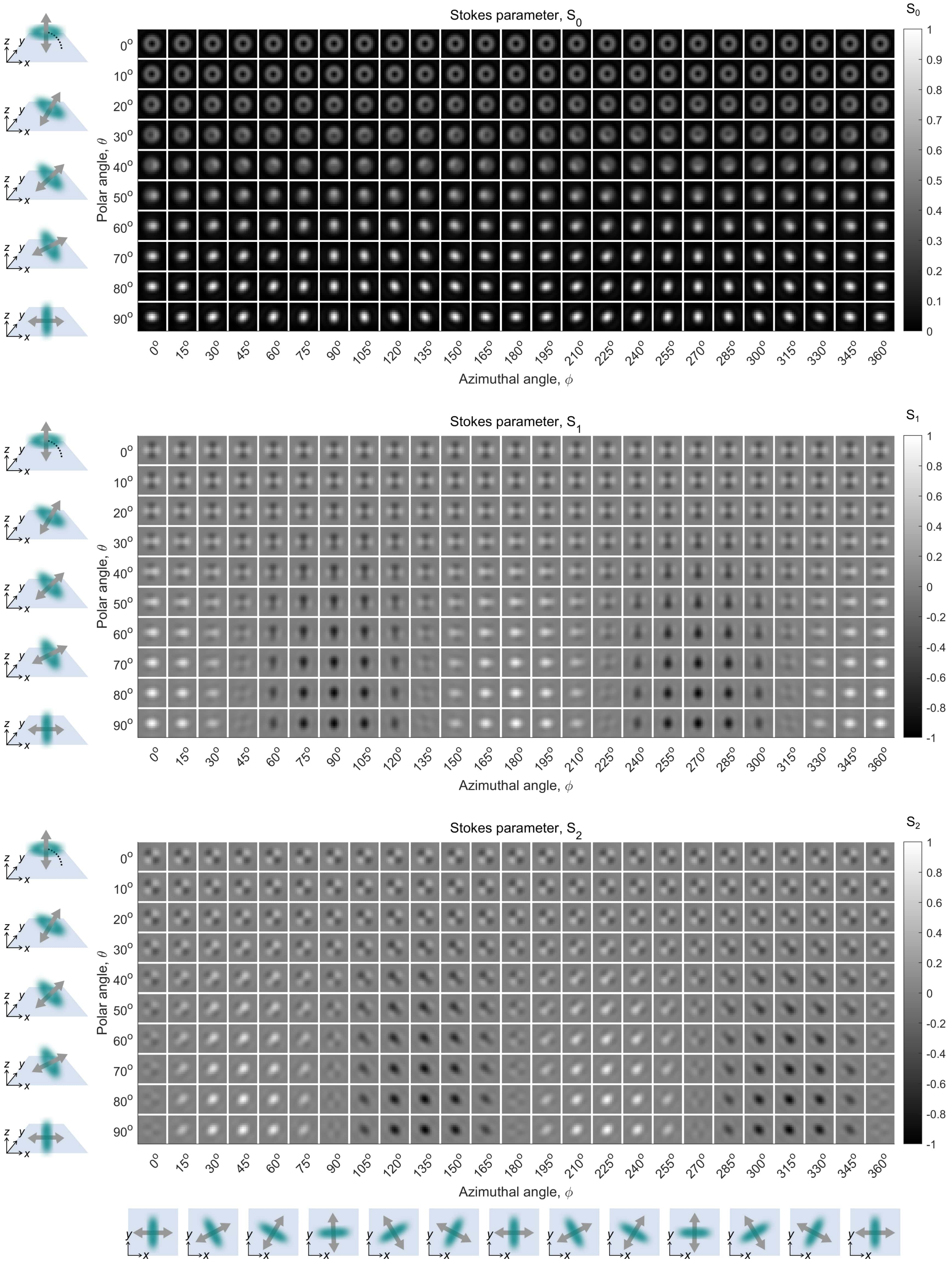

**Figure S8: Simulated image plane for different dipole orientations: Stokes parameters.** Simulated Stokes parameter images at the image plane for an immobilized single fluorescent molecule at different 3D orientations. The simulation parameters used to generate this figure are described in section S3. The molecules are positioned on a planar refractive index interface between glass (n=1.158) and water (n=1.33).

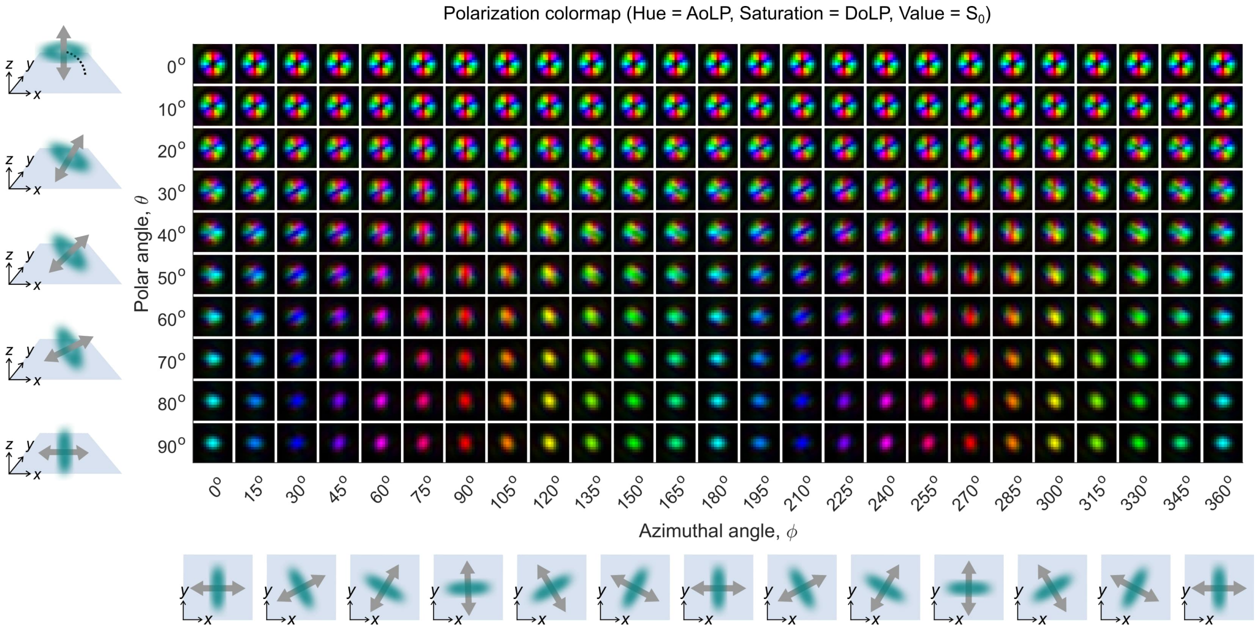

**Figure S9: Simulated image plane for different dipole orientations: Polarization.** Simulated images at the image plane for an immobilized single fluorescent molecule at different 3D orientations, visualized using a polarization colormap that combines AoLP, DoLP, and $S_0$ in HSV colorspace. The simulation parameters used to generate this figure are described in section S3. The molecules are positioned on a planar refractive index interface between glass (n=1.158) and water (n=1.33).

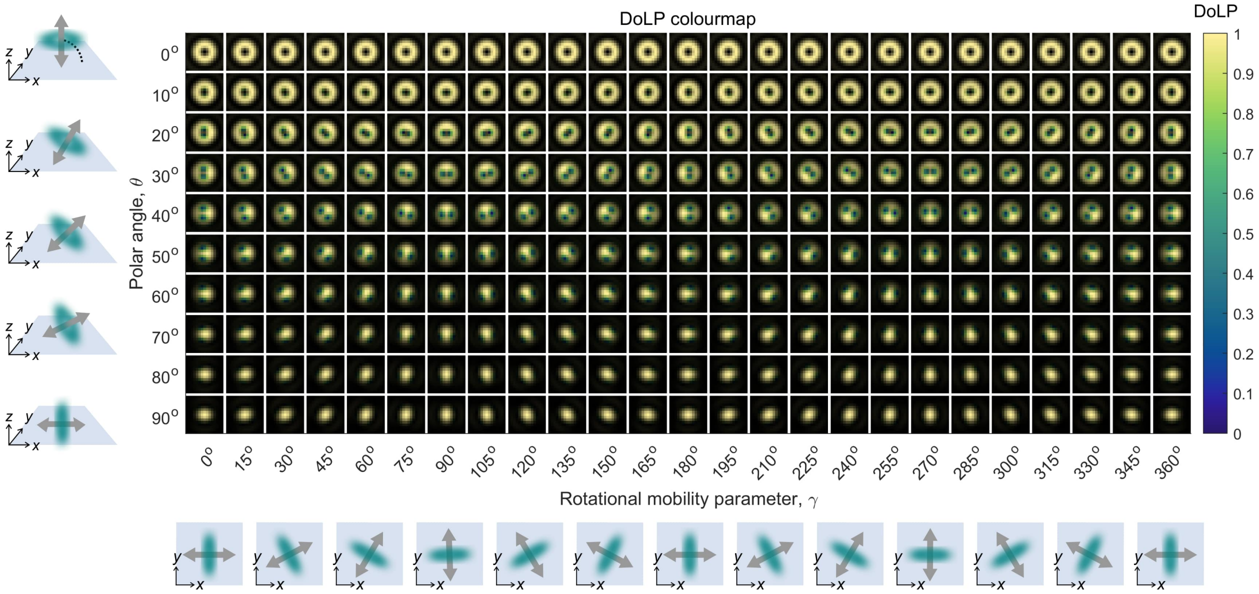

**Figure S10: Simulated polarization camera images for different dipole orientations: Degree of Linear Polarization.** Simulated images at the image plane for an immobilized single fluorescent molecule at different 3D orientations, visualized using a DoLP colormap that combines DoLP and $S_0$. The simulation parameters used to generate this figure are described in section S3. The molecules are positioned on a planar refractive index interface between glass (n=1.158) and water (n=1.33).

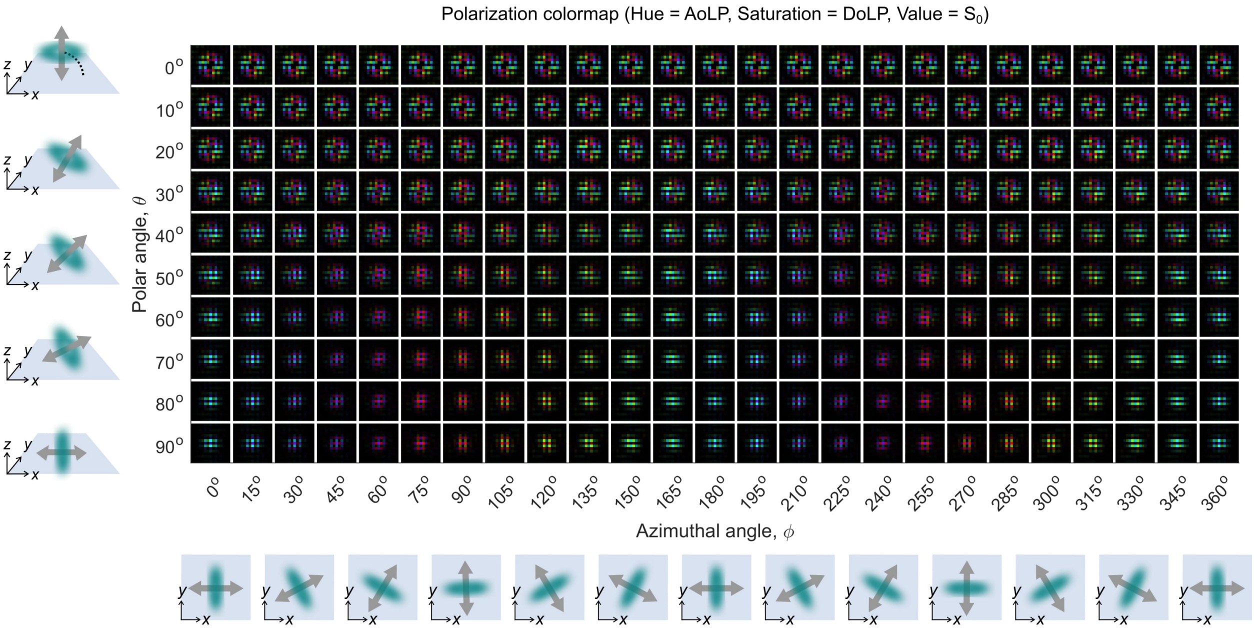

**Figure S11: Simulated polarization camera images for different dipole orientations: Polarization.** Simulated polarisation camera images at the image plane for an immobilized single fluorescent molecule at different 3D orientations, visualized using a polarization colormap that combines AoLP, DoLP, and $S_0$ in HSV colorspace. Pixels with a micropolarizer oriented at $0°$, $45°$, $90°$ and $-45°$ relative to the x-axis will respectively appear cyan, purple, red, and green. The simulation parameters used to generate this figure are described in section S3. The molecules are positioned on a planar refractive index interface between glass (n=1.158) and water (n=1.33).

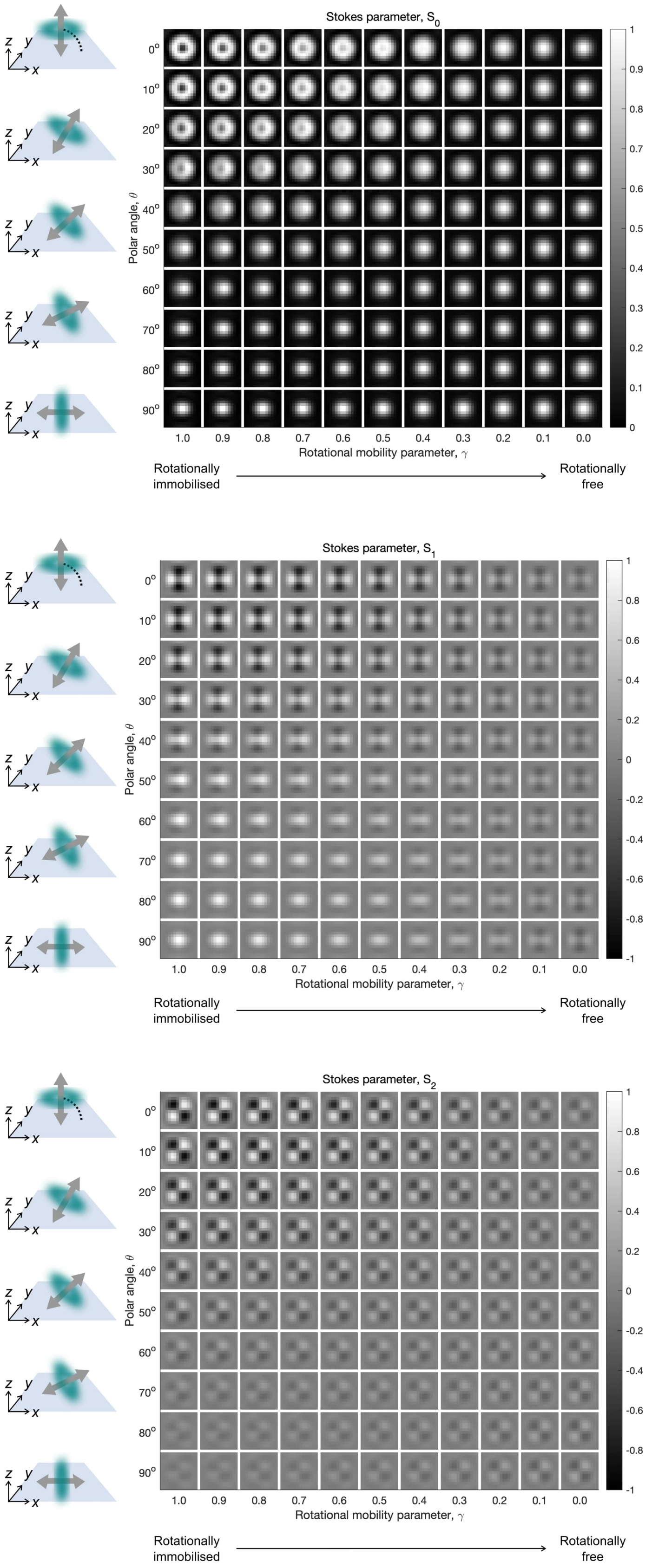

**Figure S12: Simulated image plane for different rotational mobilities, $\gamma$ ($\phi = 0°$).** Simulated Stokes parameter images ($S_0$, $S_1$, and $S_2$) at the image plane for a single fluorescent molecule at different 3D orientations, and rotational mobility parameters $\gamma$. The in-plane angle $\phi$ is fixed to $0°$. The simulation parameters used to generate this figure are described in section S3. The molecules are positioned on a planar refractive index interface between glass (n=1.158) and water (n=1.33). No noise was added.

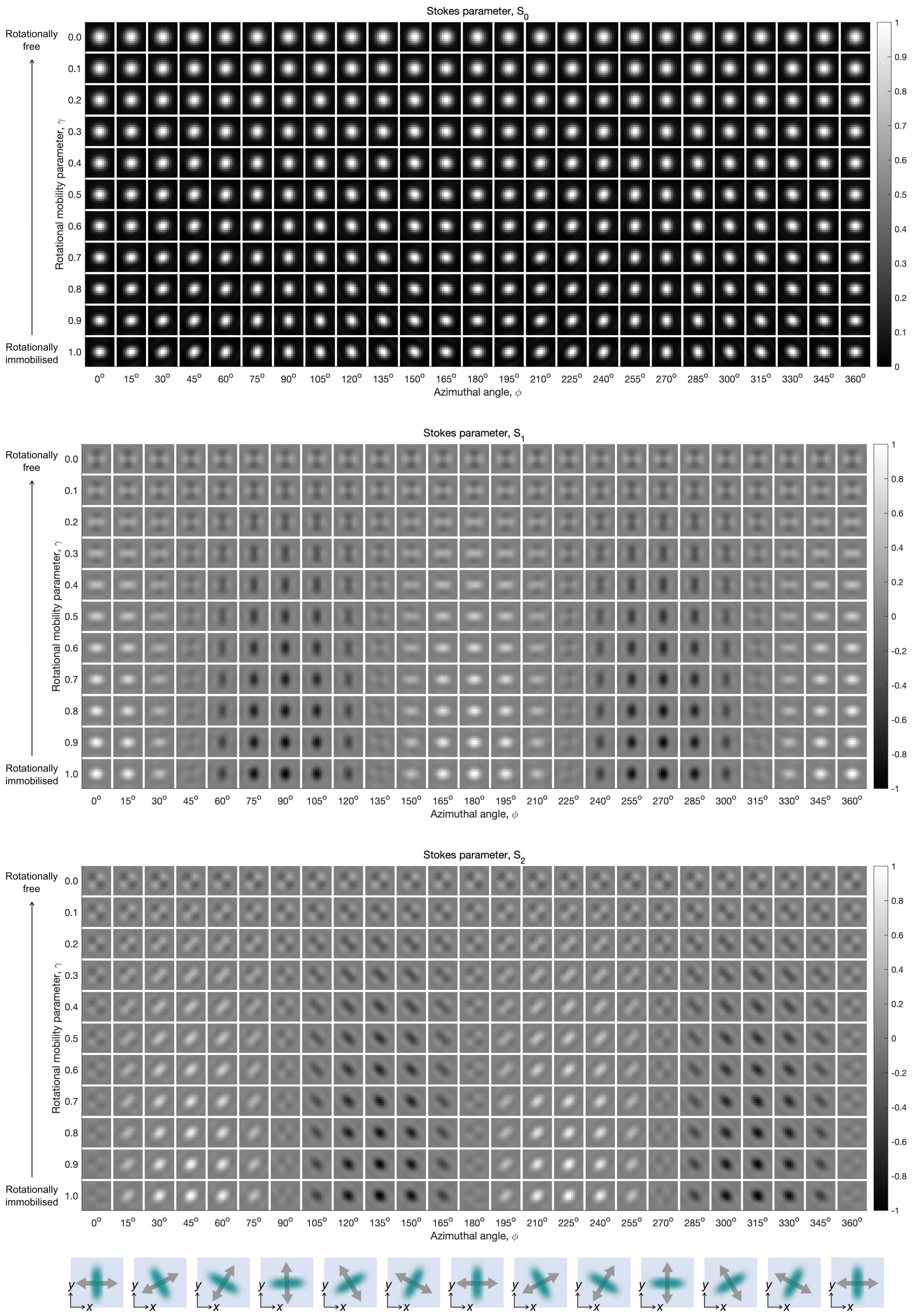

**Figure S13: Simulated image plane for different rotational mobilities, $\gamma$ ($\theta = 90°$).** Simulated Stokes parameter images ($S_0$, $S_1$, and $S_2$) at the image plane for a single fluorescent molecule at different 3D orientations, and rotational mobility parameters $\gamma$. The out-of-plane angle $\theta$ is fixed to $90°$. The simulation parameters used to generate this figure are described in section S3. The molecules are positioned on a planar refractive index interface between glass (n=1.158) and water (n=1.33). No noise was added.

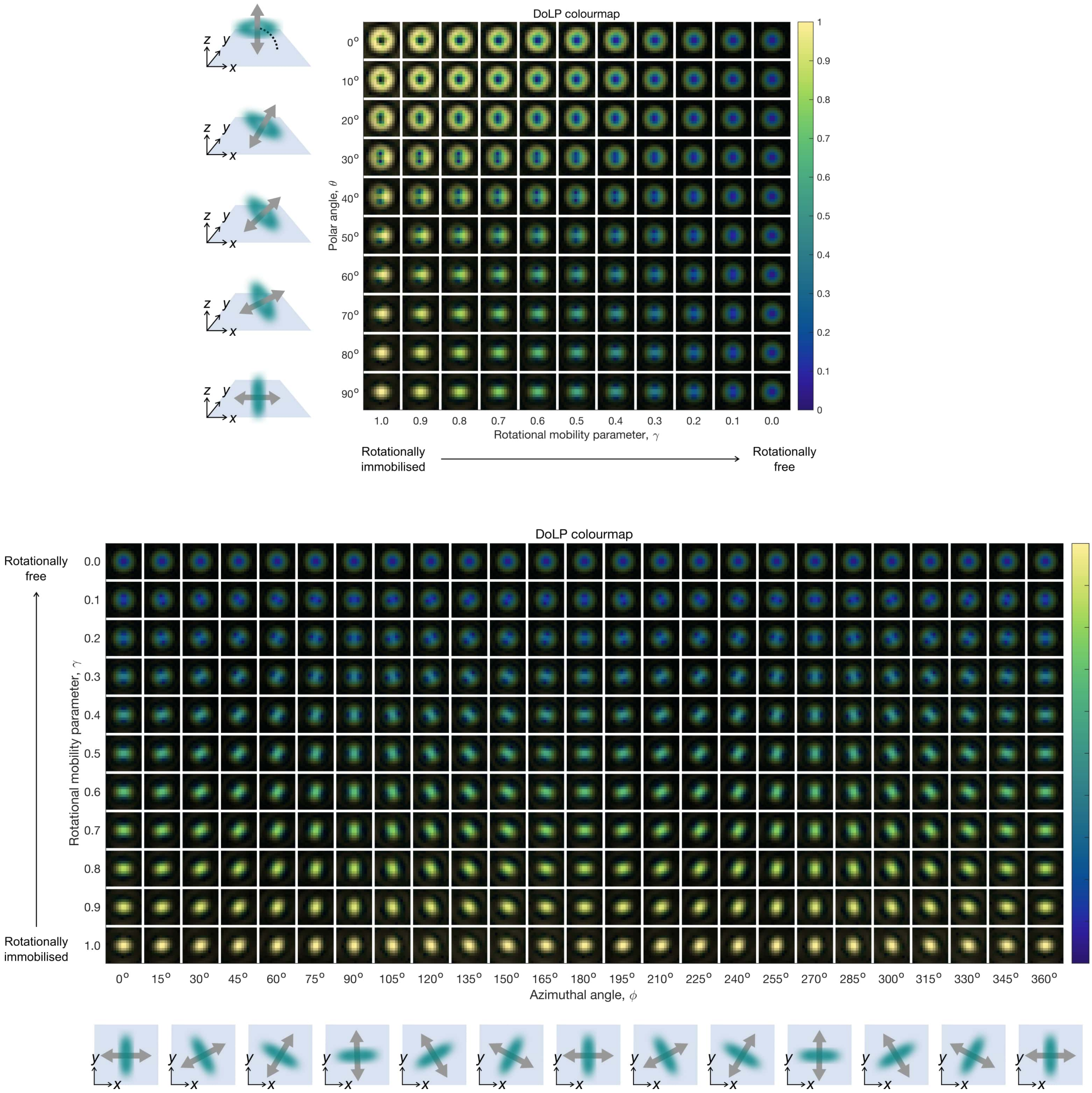

**Figure S14: Simulated image plane for different rotational mobilities, $\gamma$: Degree of Linear Polarization.** Simulated image plane for a single fluorescent molecule at different 3D orientations, and rotational mobility parameters $\gamma$, shown using a DoLP colormap representation that combines DoLP and $S_0$. In the top figure, the in-plane angle $\phi$ is fixed to $0°$. In the bottom figure, the out-of-plane angle $\theta$ is fixed to $90°$.The simulation parameters used to generate this figure are described in section S3. The molecules are positioned on a planar refractive index interface between glass (n=1.158) and water (n=1.33). No noise was added.

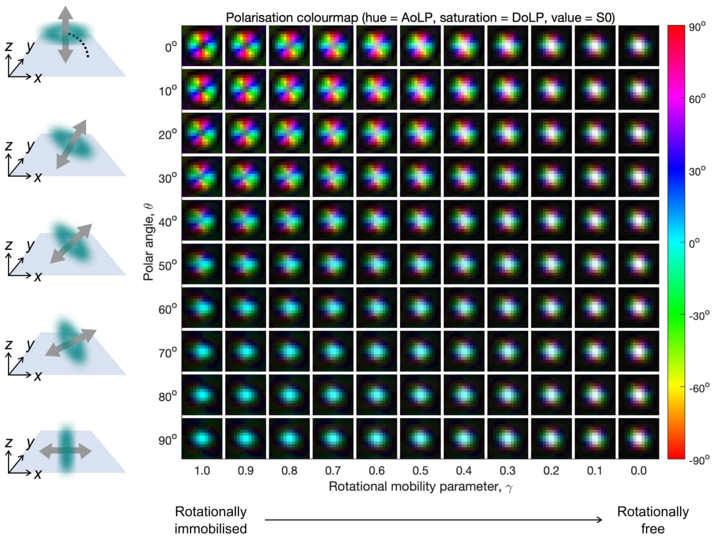

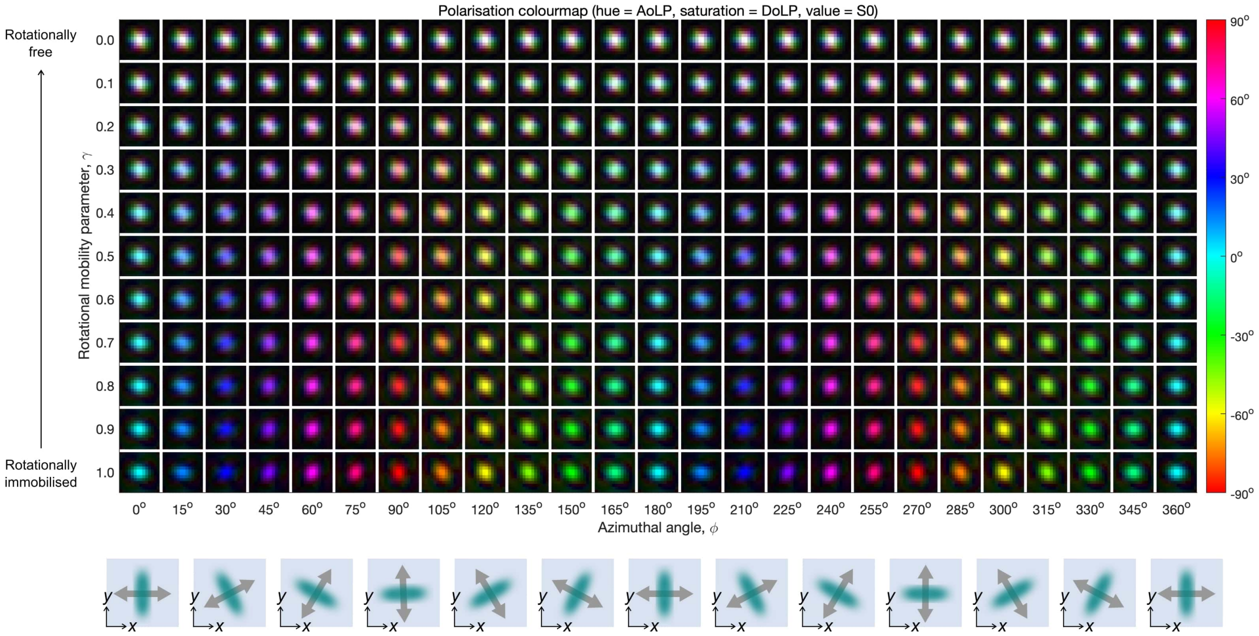

**Figure S15: Simulated image plane for different rotational mobilities, $\gamma$: Polarization.** Simulated image plane for a single fluorescent molecule at different 3D orientations, and rotational mobility parameters $\gamma$, shown using a polarization colormap representation that combines AoLP, DoLP, and $S_0$ in HSV colorspace. In the top figure, the in-plane angle $\phi$ is fixed to $0°$. In the bottom figure, the out-of-plane angle $\theta$ is fixed to $90°$.The simulation parameters used to generate this figure are described in section S3. The molecules are positioned on a planar refractive index interface between glass (n=1.158) and water (n=1.33). No noise was added.

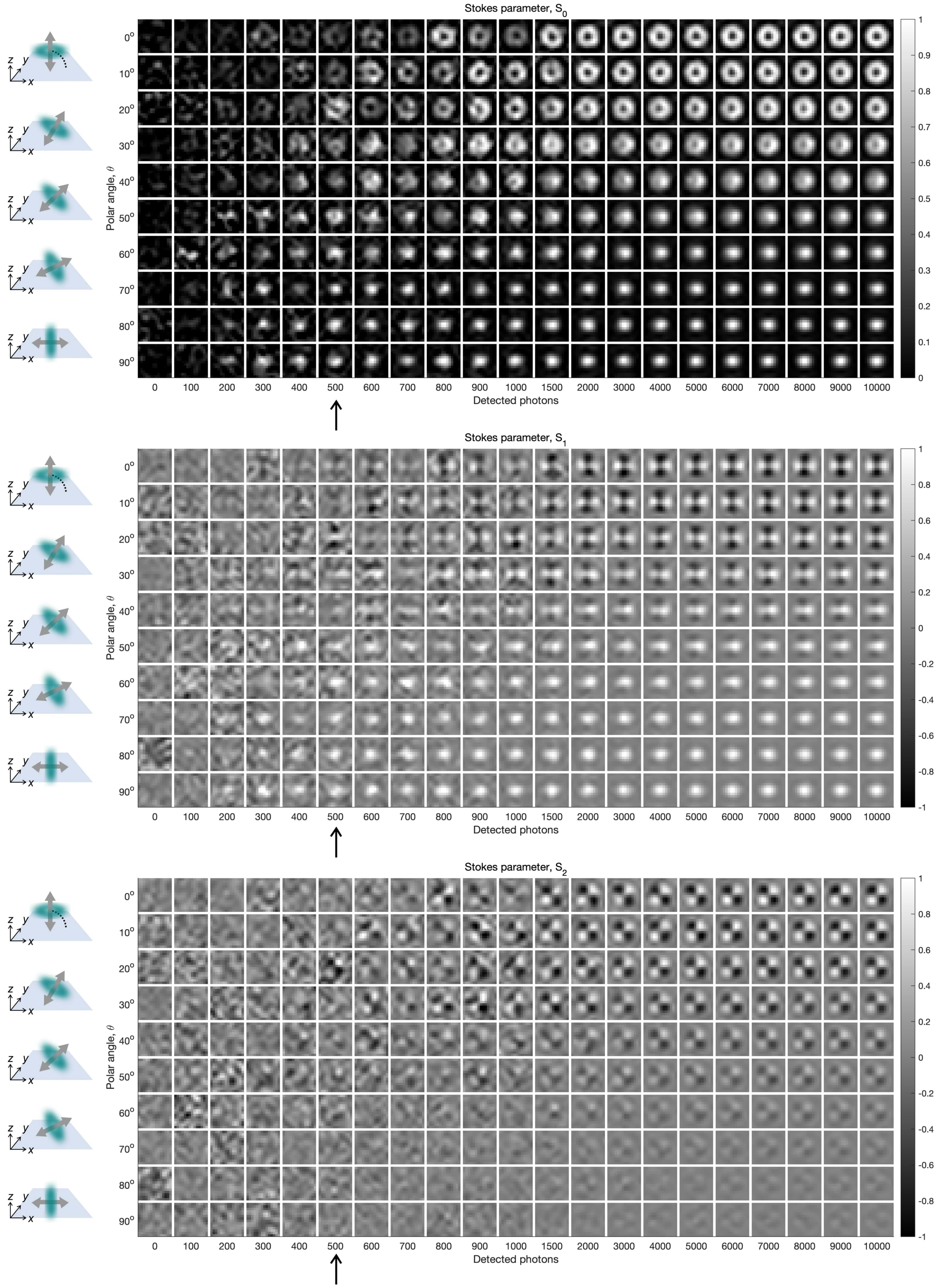

**Figure S16: Simulated image plane at different SNR ($\phi = 0°$).** Stokes parameter images ($S_0$, $S_1$, and $S_2$) at the image plane, estimated from simulated noisy polarization camera images of a single fluorescent molecule at different 3D orientations and signal-to-noise ratios. The in-plane angle $\phi$ is fixed to $0°$. The simulation parameters used to generate this figure are described in section S3. The molecules are positioned on a planar refractive index interface between glass (n=1.158) and water (n=1.33). To generate the noisy images, the detection noise model described in section S1.7 was used, using the experimentally measured camera calibration parameters in Table 2. A fixed background of 10 photons per pixel was also added to all images. Channel interpolation was performed using cubic spline interpolation.

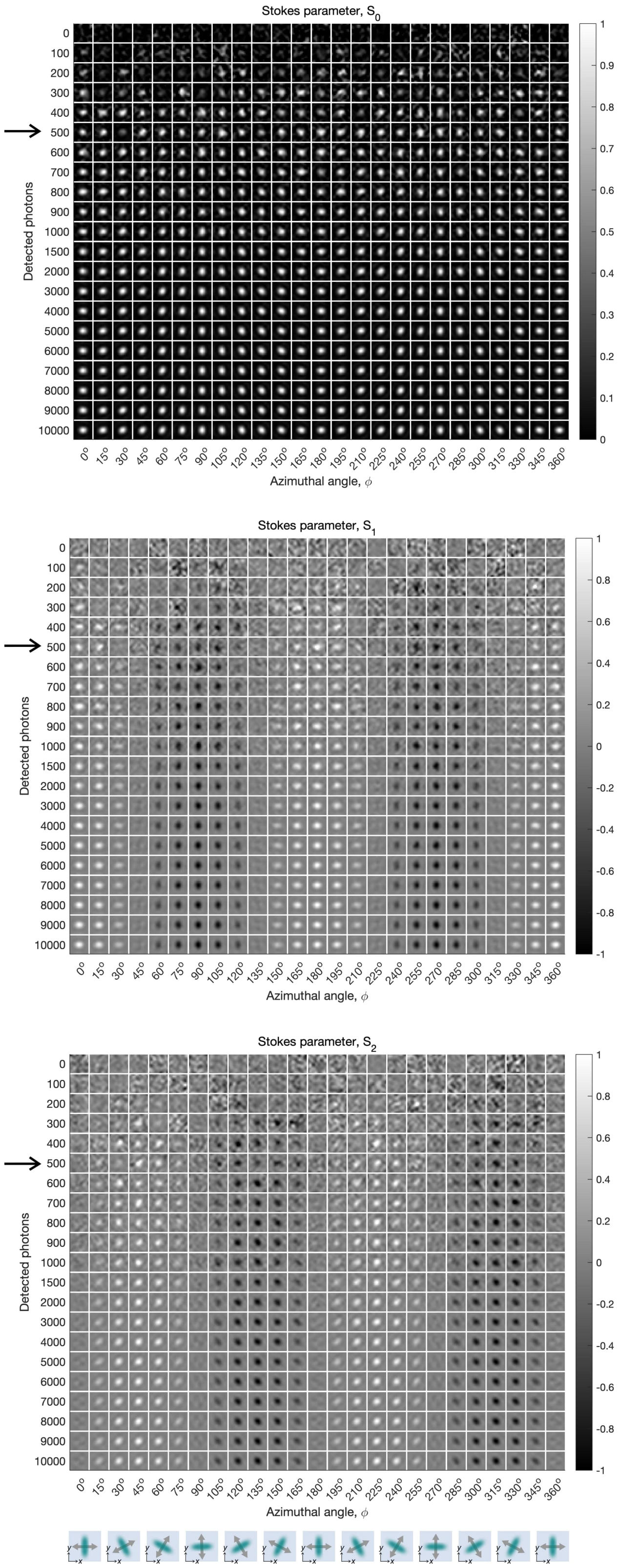

**Figure S17: Simulated image plane at different SNR ($\theta = 90°$).** Stokes parameter images ($S_0$, $S_1$, and $S_2$) at the image plane, estimated from simulated noisy polarization camera images of a single fluorescent molecule at different 3D orientations and signal-to-noise ratios. The out-of-plane angle $\theta$ is fixed to $90°$. The simulation parameters used to generate this figure are described in section S3. The molecules are positioned on a planar refractive index interface between glass (n=1.158) and water (n=1.33). To generate the noisy images, the detection noise model described in section S1.7 was used, using the experimentally measured camera calibration parameters in Table 2. A fixed background of 10 photons per pixel was also added to all images. Channel interpolation was performed using cubic spline interpolation.

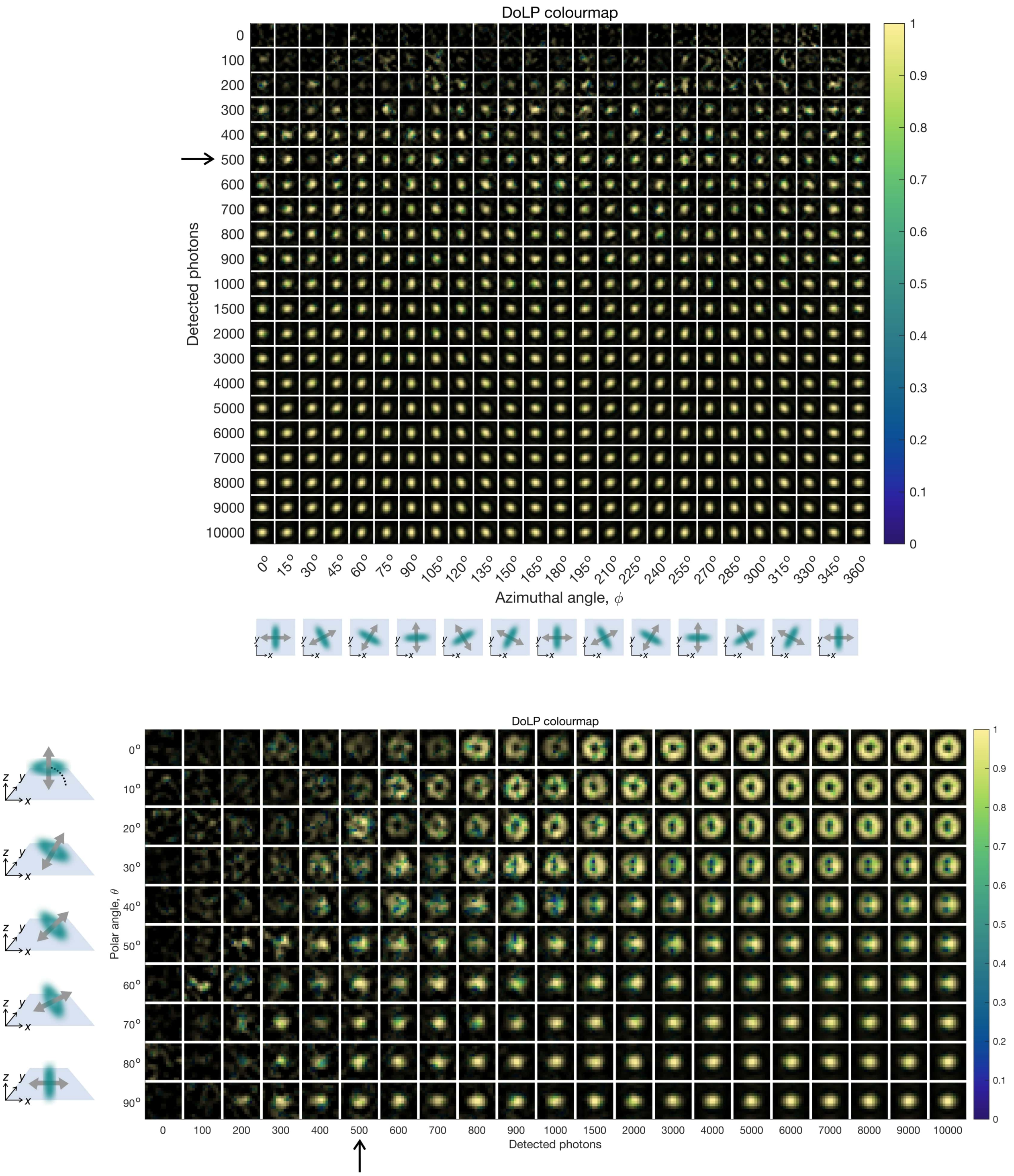

**Figure S18: Simulated image at different SNR: Degree of Linear Polarization.** DoLP colormap representation that combines DoLP and $S_0$, estimated from simulated noisy polarization camera images of a single fluorescent molecule at different 3D orientations and signal-to-noise ratios. The simulation parameters used to generate this figure are described in section S3. The molecules are positioned on a planar refractive index interface between glass (n=1.158) and water (n=1.33). To generate the noisy images, the detection noise model described in section S1.7 was used, using the experimentally measured camera calibration parameters in Table 2. A fixed background of 10 photons per pixel was also added to all images. Channel interpolation was performed using cubic spline interpolation. In the top figure, the out-of-plane angle $\theta$ is fixed to $90°$. In the bottom figure, the in-plane angle $\phi$ is fixed to $0°$.

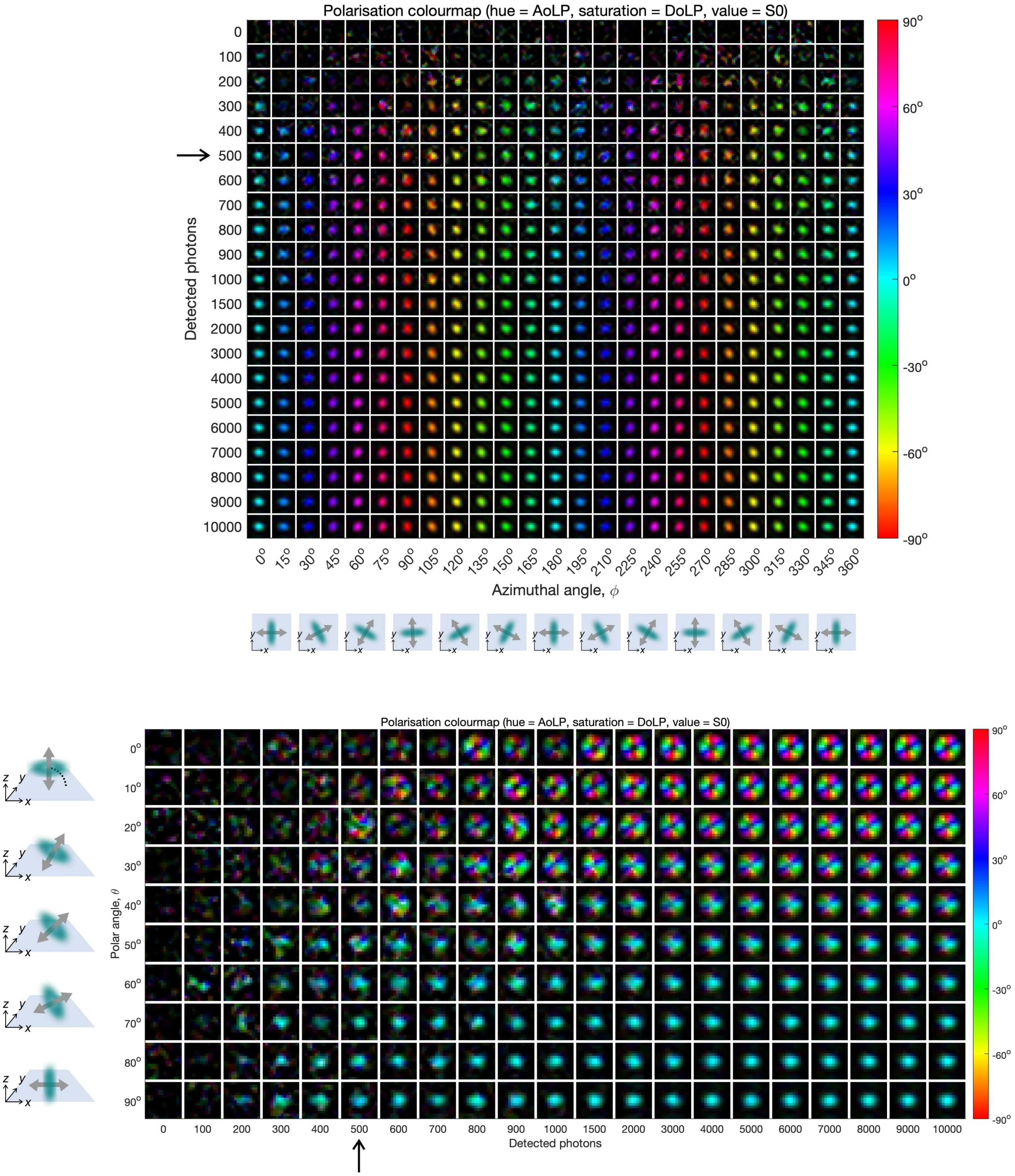

**Figure S19: Simulated image plane at different SNR: Polarization.** Polarisation colormap representation that combines AoLP, DoLP, and $S_0$ in HSV colorspace, estimated from simulated noisy polarization camera images of a single fluorescent molecule at different 3D orientations and signal-to-noise ratios. The simulation parameters used to generate this figure are described in section S3. The molecules are positioned on a planar refractive index interface between glass (n=1.158) and water (n=1.33). To generate the noisy images, the detection noise model described in section S1.7 was used, using the experimentally measured camera calibration parameters in Table 2. A fixed background of 10 photons per pixel was also added to all images. Channel interpolation was performed using cubic spline interpolation. In the top figure, the out-of-plane angle $\theta$ is fixed to $90°$. In the bottom figure, the in-plane angle $\phi$ is fixed to $0°$.

### S4 Expressions for phi and theta

#### S4.1 Stokes parameters, AoLP and DoLP

Stokes parameters describe the polarization state of an optical beam and are expressed in terms of intensities. There are four Stokes parameter $\mathbf{S} = (S_0, S_1, S_2, S_3)$ for which

$$S_0^2 \geqslant S_1^2 + S_2^2 + S_3^2 \tag{19}$$

The equality holds for completely polarized light. The parameter $S_0$ describes the total intensity of the optical field, $S_1$ describes the preponderance of linearly horizontally polarized light over linearly vertically polarized light, $S_2$ describes the preponderance of linear +45° polarized light over linear -45° polarized light and $S_3$ describes the preponderance of right circularly polarized light over left circularly polarized light [4].

The polarization camera used in this work has pixels with polarizers with transmission axis orientations at $0°$, $45°$, $90°$ and $-45°$ relative to a horizontal axis (specified on the camera body). The intensity measured in these four polarized detection channels can be used to calculate Stokes parameters $S_0$, $S_1$, and $S_2$, which in turn can be used to calculate the angle of linear polarization (AoLP) and degree of linear polarization (DoLP) as follows:

$$\text{AoLP} = \frac{1}{2}\tan^{-1}\left(\frac{S_2}{S_1}\right) \tag{20}$$

$$\text{DoLP} = \sqrt{\frac{S_1^2 + S_2^2}{S_0^2}} \tag{21}$$

#### S4.2 Calculating Stokes parameters from polarized measurements

The relationship between the measured intensities in the four channels and the Stokes parameters is well known, but we include the derivation here for completeness. Mueller matrices can be used to calculate the Stokes parameters of light after passing through an optical element, e.g. a linear polarizer [4]. The Mueller matrix for an ideal linear polarizer with its transmission axis at an angle $\eta$ to the x-axis is

$$\mathbf{M}_{LP}(\eta) = \frac{1}{2}\begin{bmatrix} 1 & \cos(2\eta) & \sin(2\eta) & 0 \\ \cos(2\eta) & \cos^2(2\eta) & \sin(2\eta)\cos(2\eta) & 0 \\ \sin(2\eta) & \sin(2\eta)\cos(2\eta) & \sin^2(2\eta) & 0 \\ 0 & 0 & 0 & 0 \end{bmatrix} \tag{22}$$

The factor $1/2$ is true for an ideal linear polarizer. We consider a beam with a Stokes vector $\mathbf{S} = (S_0, S_1, S_2, S_3)^T$ incident on the linear polarizer. The emerging beam will have a modified Stokes vector $\mathbf{S}'$

$$\mathbf{S}' = \mathbf{M}_{LP}(\theta)\mathbf{S} \tag{23}$$

$$= \begin{bmatrix} 1 & \cos(2\eta) & \sin(2\eta) & 0 \\ \cos(2\eta) & \cos^2(2\eta) & \sin(2\eta)\cos(2\eta) & 0 \\ \sin(2\eta) & \sin(2\eta)\cos(2\eta) & \sin^2(2\eta) & 0 \\ 0 & 0 & 0 & 0 \end{bmatrix}\begin{bmatrix} S_0 \\ S_1 \\ S_2 \\ S_3 \end{bmatrix} \tag{24}$$

$$= \begin{bmatrix} S_0 + S_1\cos(2\eta) + S_2\sin(2\eta) \\ S_0\cos(2\eta) + S_1\cos^2(2\eta) + S_2\sin(2\eta)\cos(2\eta) \\ S_0\sin(2\eta) + S_1\sin(2\eta)\cos(2\eta) + S_2\sin^2(2\eta) \\ 0 \end{bmatrix} \tag{25}$$

The intensity of the emerging beam is its first Stokes parameter

$$I(\eta) = S_0'(\eta) = S_0 + S_1\cos(2\eta) + S_2\sin(2\eta) \tag{26}$$

Plugging the orientations of the polarizer elements of the polarizer microarray $\eta = \{0°, 45°, 90°, -45°\}$ into Eq. (26) and solving for the Stokes parameters yields [box=]align $\text{S}_0 = I(0°) + I(90°) = I(45°) + I(-45°)$
$S_1 = I(0°) - I(90°)$
$S_2 = I(45°) - I(-45°)$ We can only measure the first three Stokes parameters using our polarization camera and setup [9–11].

#### S4.3 Rewriting expressions for $\phi$ and $\theta$ as a function of Stokes parameters

As derived by John T. Fourkas in [12], the in-plane and out-of-plane orientation of the emission dipole moment of a fluorescent molecule can be determined from polarized detection measurements using simple equations. The following equations for $\phi$, $\theta$ and $I_{tot}$ are taken directly from reference [12]:

$$\phi = \frac{1}{2}\tan^{-1}\left[\left(I_{45} - \frac{I_0 + I_{90}}{2}\right) \Big/ \left(\frac{I_0 - I_{90}}{2}\right)\right] \tag{28}$$

$$\theta = \sin^{-1}\left[\left(\frac{I_0 - I_{90}}{2CI_{tot}\cos(2\phi)}\right)^{1/2}\right] \tag{29}$$

$$I_{tot} = \frac{1}{2A}\left[\left(1 - \frac{B}{C\cos(2\phi)}\right)I_0 + \left(1 + \frac{B}{C\cos(2\phi)}\right)I_{90}\right] \tag{30}$$

where the coefficients $A, B$ and $C$ are functions of the maximum collection angle $\alpha = \sin^{-1}(\mathrm{NA}/n)$ of the objective:

$$\begin{aligned} A &= \frac{1}{6} - \frac{1}{4}\cos\alpha + \frac{1}{12}\cos^3\alpha \\ B &= \frac{1}{8}\cos\alpha - \frac{1}{8}\cos^3\alpha \\ C &= \frac{7}{48} - \frac{1}{16}\cos\alpha - \frac{1}{16}\cos^2\alpha - \frac{1}{48}\cos^3\alpha \end{aligned} \tag{32}$$

In this work, we rewrite these equations in terms of Stokes parameters by substituting Eq. (27) into equations (28), (29) and (30), resulting in:

[box=]align $\phi = \frac{1}{2}\tan^{-1}\left(\frac{S_2}{S_1}\right)$

$$\theta = \sin^{-1}\left(\sqrt{\frac{A \cdot \text{netDoLP}}{C - B \cdot \text{netDoLP}}}\right)$$

where the coefficients $A, B$ and $C$ are defined as in Eq. (31) and the net degree of linear polarization (netDoLP) is defined as

$$\boxed{\text{netDoLP} = \sqrt{\frac{\langle S_1\rangle^2 + \langle S_2\rangle^2}{\langle S_0\rangle^2}}} \tag{34}$$

The brackets $\langle ... \rangle$ represent spatial averaging. We also use another DoLP-based metric which we call avgDoLP and define as

$$\boxed{\text{avgDoLP} = \langle \text{DoLP} \rangle = \left\langle \sqrt{\frac{S_1^2 + S_2^2}{S_0^2}} \right\rangle} \tag{35}$$

We use avgDoLP as a proxy for rotational mobility.

### S5 Derivation of expressions for $\phi$ and $\theta$

To derive equations for the in-plane angle $\phi$ and the out-of-plane angle $\theta$ of the emission dipole moment of an immobilised emitter, we will use an approach similar to the one used in a publication by Fourkas [12]. Fourkas used Eq. (12) to describe the electric field emitted by a single immobilised fluorescent molecule at the back focal plane of the objective lens. They subsequently used Jones matrices to derive expressions for the electric field after a polariser in the back focal plane, with a transmission axis respectively at $0°$, $45°$, $90°$ and $-45°$ degrees from the x-axis, measured anti-clockwise. For each polarised channel, they then square the electric field to get the intensity of the field after the polariser, and integrate over the collection cone of the objective lens. The result is four equations for the measured intensity in the four polarised channels as a function of the orientation of the emission dipole moment; $I_0(\phi,\theta)$, $I_{45}(\phi,\theta)$, $I_{90}(\phi,\theta)$ and $I_{-45}(\phi,\theta)$. These expressions can then be rearranged to arrive at formulas for the in-plane angle $\phi$ and the out-of-plane angle $\theta$ as a function of the measured intensities in the four polarised channels.

#### S5.1 Intensity after a linear polariser as a function of dipole orientation

In this section, the same derivation will be performed as presented in section ??, but including a linear polariser in the detection path. We describe the effect of a linear polariser on the measured intensity by multiplying the electric field from Eq. (12) with a Jones matrix **J**:

$$\mathbf{E}_{\text{bfp}}(\phi',\theta') \propto \mathbf{J}\mathbf{G}_{\text{bfp}}(\phi',\theta')\hat{\mu} \tag{36}$$

The Jones matrices for a linear polariser with a transmission axis oriented at respectively $0°, 45°, 90°$ and $-45°$ degrees (anti-clockwise with respect to the x-axis) are [4]

$$\mathbf{J}_{0°} = \begin{bmatrix} 1 & 0 & 0 \\ 0 & 0 & 0 \\ 0 & 0 & 0 \end{bmatrix}, \quad \mathbf{J}_{90°} = \begin{bmatrix} 0 & 0 & 0 \\ 0 & 1 & 0 \\ 0 & 0 & 0 \end{bmatrix}, \quad \mathbf{J}_{45°} = \frac{1}{2}\begin{bmatrix} 1 & 1 & 0 \\ 1 & 1 & 0 \\ 0 & 0 & 0 \end{bmatrix}, \quad \mathbf{J}_{-45°} = \frac{1}{2}\begin{bmatrix} 1 & -1 & 0 \\ -1 & 1 & 0 \\ 0 & 0 & 0 \end{bmatrix} \tag{37}$$

The measured intensity will be the square of the electric field integrated over the collection cone of the objective:

$$I = \int_0^{2\pi}\int_0^{\alpha} |\mathbf{E}|^2 \sin\theta' d\theta' d\phi' = \int_0^{2\pi}\int_0^{\alpha} \left(E_x^2 + E_y^2 + E_z^2\right)\sin\theta' d\theta' d\phi' \tag{38}$$

##### S5.1.1 Linear polariser with transmission axis at $0°$

We first get the electric field after a linear polariser with transmission axis at $0°$ as described in Eq. (36) and then calculate the total measured intensity after the polariser using Eq. (38):

$$\begin{aligned}
\mathbf{E}_{\text{bfp},0°}(\phi',\theta') &= \mathbf{J}_{0°}\mathbf{G}_{\text{bfp}}(\phi',\theta')\hat{\mu} \\
&= \begin{bmatrix} 1 & 0 & 0 \\ 0 & 0 & 0 \\ 0 & 0 & 0 \end{bmatrix} \begin{bmatrix} G_{xx} & G_{xy} & G_{xz} \\ G_{yx} & G_{yy} & G_{yz} \\ G_{zx} & G_{zy} & G_{zz} \end{bmatrix} \begin{bmatrix} \mu_x \\ \mu_y \\ \mu_z \end{bmatrix} \\
&= \begin{bmatrix} G_{xx} & G_{xy} & G_{xz} \\ 0 & 0 & 0 \\ 0 & 0 & 0 \end{bmatrix} \begin{bmatrix} \mu_x \\ \mu_y \\ \mu_z \end{bmatrix} \\
&= \begin{bmatrix} G_{xx}\mu_x + G_{xy}\mu_y + G_{xz}\mu_z \\ 0 \\ 0 \end{bmatrix}
\end{aligned}$$

$$\begin{aligned}
I_{0°}(\boldsymbol{\mu}) &= \int_0^{2\pi}\int_0^{\alpha} |\mathbf{E}_{\text{bfp},0°}(\phi',\theta')|^2 \sin\theta' d\theta' d\phi' \\
&= \int_0^{2\pi}\int_0^{\alpha} (G_{xx}\mu_x + G_{xy}\mu_y + G_{xz}\mu_z)^2 \sin\theta' d\theta' d\phi' \\
&= C_1\mu_x^2 + C_2\mu_y^2 + C_3\mu_z^2
\end{aligned}$$

where the value of $C_1$, $C_2$ and $C_3$ can be found in section S5.4.

##### S5.1.2 Linear polariser with transmission axis at $90°$

We first get the electric field after a linear polariser with transmission axis at $90°$ as described in Eq. (36) and then calculate the total measured intensity after the polariser using Eq. (38):

$$
\begin{aligned}
\mathbf{E}_{\text{bfp},90°}(\phi', \theta') &= \mathbf{J}_{90°}\mathbf{G}_{\text{bfp}}(\phi', \theta')\hat{\mu} \\
&= \begin{bmatrix} 0 & 0 & 0 \\ 0 & 1 & 0 \\ 0 & 0 & 0 \end{bmatrix} \begin{bmatrix} G_{xx} & G_{xy} & G_{xz} \\ G_{yx} & G_{yy} & G_{yz} \\ G_{zx} & G_{zy} & G_{zz} \end{bmatrix} \begin{bmatrix} \mu_x \\ \mu_y \\ \mu_z \end{bmatrix} \\
&= \begin{bmatrix} 0 & 0 & 0 \\ G_{yx} & G_{yy} & G_{yz} \\ 0 & 0 & 0 \end{bmatrix} \begin{bmatrix} \mu_x \\ \mu_y \\ \mu_z \end{bmatrix} \\
&= \begin{bmatrix} 0 \\ G_{yx}\mu_x + G_{yy}\mu_y + G_{yz}\mu_z \\ 0 \end{bmatrix}
\end{aligned}
$$

$$
\begin{aligned}
I_{90°}(\boldsymbol{\mu}) &= \int_0^{2\pi}\int_0^{\alpha} |\mathbf{E}_{\text{bfp},90°}(\phi', \theta')|^2 \sin\theta' d\theta' d\phi' \\
&= \int_0^{2\pi}\int_0^{\alpha} (G_{yx}\mu_x + G_{yy}\mu_y + G_{yz}\mu_z)^2 \sin\theta' d\theta' d\phi' \\
&= C_2\mu_x^2 + C_1\mu_y^2 + C_3\mu_z^2
\end{aligned}
$$

where the value of $C_1$, $C_2$ and $C_3$ can be found in section S5.4.

##### S5.1.3 Linear polariser with transmission axis at $45°$

We first get the electric field after a linear polariser with transmission axis at $45°$ as described in Eq. (36) and then calculate the total measured intensity after the polariser using Eq. (38):

$$
\begin{aligned}
\mathbf{E}_{\text{bfp},45°}(\phi', \theta') &= \mathbf{J}_{45°}\mathbf{G}_{\text{bfp}}(\phi', \theta')\hat{\mu} \\
&= \frac{1}{2}\begin{bmatrix} 1 & 1 & 0 \\ 1 & 1 & 0 \\ 0 & 0 & 0 \end{bmatrix} \begin{bmatrix} G_{xx} & G_{xy} & G_{xz} \\ G_{yx} & G_{yy} & G_{yz} \\ G_{zx} & G_{zy} & G_{zz} \end{bmatrix} \begin{bmatrix} \mu_x \\ \mu_y \\ \mu_z \end{bmatrix} \\
&= \frac{1}{2}\begin{bmatrix} G_{xx} + G_{yx} & G_{xy} + G_{yy} & G_{xz} + G_{yx} \\ G_{xx} + G_{yx} & G_{xy} + G_{yy} & G_{xz} + G_{yx} \\ 0 & 0 & 0 \end{bmatrix} \begin{bmatrix} \mu_x \\ \mu_y \\ \mu_z \end{bmatrix} \\
&= \frac{1}{2}\begin{bmatrix} (G_{xx} + G_{yx})\mu_x + (G_{xy} + G_{yy})\mu_y + (G_{xz} + G_{yz})\mu_z \\ (G_{xx} + G_{yx})\mu_x + (G_{xy} + G_{yy})\mu_y + (G_{xz} + G_{yz})\mu_z \\ 0 \end{bmatrix}
\end{aligned}
$$

$$
\begin{aligned}
I_{45°}(\boldsymbol{\mu}) &= \int_0^{2\pi}\int_0^{\alpha} |\mathbf{E}_{\text{bfp},45°}(\phi', \theta')|^2 \sin\theta' d\theta' d\phi' \\
&= \frac{1}{4}\int_0^{2\pi}\int_0^{\alpha} \Big( \big[(G_{xx} + G_{yx})\mu_x + (G_{xy} + G_{yy})\mu_y + (G_{xz} + G_{yz})\mu_z\big]^2 \\
&\qquad + \big[(G_{xx} + G_{yx})\mu_x + (G_{xy} + G_{yy})\mu_y + (G_{xz} + G_{yz})\mu_z\big]^2 \Big) \sin\theta' d\theta' d\phi' \\
&= \frac{(C_1 + C_2)}{2}(\mu_x^2 + \mu_y^2) + C_3\mu_z^2 + (C_2 + C_4)\mu_x\mu_y
\end{aligned}
$$

where the value of $C_1$, $C_2$, $C_3$ and $C_4$ can be found in section S5.4.

##### S5.1.4 Linear polariser with transmission axis at $-45°$

We first get the electric field after a linear polariser with transmission axis at $-45°$ as described in Eq. (36) and then calculate the total measured intensity after the polariser using Eq. (38):

$$\begin{aligned}
\mathbf{E}_{\text{bfp},-45°}(\phi',\theta') &= \mathbf{J}_{-45°}\mathbf{G}_{\text{bfp}}(\phi',\theta')\hat{\mu} \\
&= \frac{1}{2}\begin{bmatrix} 1 & -1 & 0 \\ -1 & 1 & 0 \\ 0 & 0 & 0 \end{bmatrix}\begin{bmatrix} G_{xx} & G_{xy} & G_{xz} \\ G_{yx} & G_{yy} & G_{yz} \\ G_{zx} & G_{zy} & G_{zz} \end{bmatrix}\begin{bmatrix} \mu_x \\ \mu_y \\ \mu_z \end{bmatrix} \\
&= \frac{1}{2}\begin{bmatrix} G_{xx}-G_{yx} & G_{xy}-G_{yy} & G_{xz}-G_{yz} \\ G_{yx}-G_{xx} & G_{yy}-G_{xy} & G_{yz}-G_{xz} \\ 0 & 0 & 0 \end{bmatrix}\begin{bmatrix} \mu_x \\ \mu_y \\ \mu_z \end{bmatrix} \\
&= \frac{1}{2}\begin{bmatrix} (G_{xx}-G_{yx})\mu_x + (G_{xy}-G_{yy})\mu_y + (G_{xz}-G_{yz})\mu_z \\ (G_{yx}-G_{xx})\mu_x + (G_{yy}-G_{xy})\mu_y + (G_{yz}-G_{xz})\mu_z \\ 0 \end{bmatrix}
\end{aligned}$$

$$\begin{aligned}
I_{-45°}(\boldsymbol{\mu}) &= \int_0^{2\pi}\int_0^{\alpha} |\mathbf{E}_{\text{bfp},-45°}(\phi',\theta')|^2 \sin\theta' d\theta' d\phi' \\
&= \frac{1}{4}\int_0^{2\pi}\int_0^{\alpha}\Big(\big[(G_{xx}-G_{yx})\mu_x + (G_{xy}-G_{yy})\mu_y + (G_{xz}-G_{yz})\mu_z\big]^2 \\
&\qquad + \big[(G_{yx}-G_{xx})\mu_x + (G_{yy}-G_{xy})\mu_y + (G_{yz}-G_{xz})\mu_z\big]^2\Big)\sin\theta' d\theta' d\phi' \\
&= \frac{(C_1+C_2)}{2}(\mu_x^2+\mu_y^2) + C_3\mu_z^2 - (C_2+C_4)\mu_x\mu_y
\end{aligned}$$

where the value of $C_1$, $C_2$, $C_3$ and $C_4$ can be found in section S5.4.

##### S5.1.5 Linear polariser: Summary of equations

We can rewrite the final expressions for the measured intensity derived in sections S5.1.1-S5.1.4 to be expressed in terms of the angles $\phi$ and $\theta$ using Eq. (8):

$$I_{0°}(\phi,\theta) = (C_1\cos^2\phi + C_2\sin^2\phi)\sin^2\theta + C_3\cos^2\theta \tag{39a}$$

$$I_{90°}(\phi,\theta) = (C_2\cos^2\phi + C_1\sin^2\phi)\sin^2\theta + C_3\cos^2\theta \tag{39b}$$

$$I_{45°}(\phi,\theta) = \frac{(C_1+C_2)}{2}\sin^2\theta + (C_2+C_4)\sin^2\theta\sin\phi\cos\phi + C_3\cos^2\theta \tag{39c}$$

$$I_{-45°}(\phi,\theta) = \frac{(C_1+C_2)}{2}\sin^2\theta - (C_2+C_4)\sin^2\theta\sin\phi\cos\phi + C_3\cos^2\theta \tag{39d}$$

#### S5.2 Stokes parameters as a function of dipole orientation

From the intensities measured in the four polarised channels, we can calculate the Stokes parameters. The first three Stokes parameters are linear combinations of Eq. (39a), (39b), (39c) and (39d):

$$S_0(\phi,\theta) = I_{0°}(\phi,\theta) + I_{90°}(\phi,\theta) = (C_1+C_2)\sin^2\theta + 2C_3\cos^2\theta \tag{40a}$$

$$S_1(\phi,\theta) = I_{0°}(\phi,\theta) - I_{90°}(\phi,\theta) = (C_1-C_2)\sin^2\theta\cos(2\phi) \tag{40b}$$

$$S_2(\phi,\theta) = I_{45°}(\phi,\theta) - I_{-45°}(\phi,\theta) = (C_2+C_4)\sin^2\theta\sin(2\phi) \tag{40c}$$

#### S5.3 Derivation of expressions for $\phi$ and $\theta$

We have derived expressions for the integrated Stokes parameters for a dipole emitter with an emission dipole moment $\boldsymbol{\mu}(\phi, \theta)$, detected using an objective with half maximum collection angle $\alpha = \sin^{-1}(\text{NA}/n)$. We rearrange Eq. (40a), (40b) and (40c), to solve for $\phi$ and $\theta$.

##### S5.3.1 In-plane angle $\phi$

We divide Stokes parameter $S_2$ by $S_1$ and simplify to get an expression for the in-plane angle $\phi$ of the emission dipole moment:

$$\frac{S_2}{S_1} = \frac{(C_1 - C_2)\sin^2\theta \sin(2\phi)}{(C_2 + C_4)\sin^2\theta\cos(2\phi)} = \frac{\sin(2\phi)}{\cos(2\phi)} = \tan(2\phi) \Leftrightarrow \boxed{\phi = \frac{1}{2}\tan^{-1}\left(\frac{S_2}{S_1}\right)} \tag{41}$$

This is also the expression for the angle of linear polarisation (AoLP), an expression commonly used in polarimetry.

##### S5.3.2 Out-of-plane angle $\theta$

To derive an expression for the out-of-plane angle, we start from the definition of the net degree of linear polarisation (netDoLP):

$$\begin{aligned}
\text{netDoLP}^2 &= \frac{S_1^2 + S_2^2}{S_0^2} \\
&= \frac{\left[(C_1 - C_2)\sin^2\theta\cos(2\phi)\right]^2 + \left[(C_2 + C_4)\sin^2\theta\sin 2\phi\right]^2}{[(C_1 + C_2)\sin^2\theta + 2C_3\cos^2\theta]^2} \\
&= \frac{\left[C_7\sin^2\theta\cos(2\phi)\right]^2 + \left[C_7\sin^2\theta\sin 2\phi\right]^2}{\left[2C_6\sin^2\theta + 2C_3\cos^2\theta\right]^2} \\
&= \frac{C_7^2\sin^4\theta}{\left[2C_6\sin^2\theta + 2C_3(1 - \sin^2\theta)\right]^2} \\
&= \frac{C_7^2\sin^4\theta}{\left[2(C_6 - C_3)\sin^2\theta + 2C_3\right]^2} \\
&= \frac{C_7^2\sin^4\theta}{4(C_6 - C_3)^2\sin^4\theta + 8C_3(C_6 - C_3)\sin^2\theta + 4C_3^2}
\end{aligned}$$

This can be written as a quadratic equation of the form $ax^2 + bx + c = 0$, where $x = \sin^2\theta$ is the unknown:

$$\left[4(C_6 - C_3)^2\text{netDoLP}^2 - C_7^2\right]\sin^4\theta + \left[8C_3(C_6 - C_3)\text{netDoLP}^2\right]\sin^2\theta + 4C_3^2\text{netDoLP}^2 = 0$$

We find, after some simplification, that:

$$\sin^2\theta = \frac{2C_3\text{netDoLP}}{2(C_3 - C_6)\text{netDoLP} \pm C_7} \equiv \frac{A \cdot \text{netDoLP}}{B \cdot \text{netDoLP} \pm C}$$

$$\theta = \sin^{-1}\left(\sqrt{\frac{A \cdot \text{netDoLP}}{B \cdot \text{netDoLP} \pm C}}\right)$$

where we defined $A = 2C_3$, $B = 2(C_3 - C_6)$ and $C = C_7$.

#### S5.4 Integrals of products of Green's tensor elements

The following is a list of integrals of all the combinations of products of elements of the Green's tensor (see Eq. (12)) of the form $G_{ij}G_{kl}$ where $i, j, k, l = \{x, y, z\}$. We integrate over $\phi$ from 0 to $2\pi$, and over $\theta$ from 0 to $\alpha$, where $\alpha$ is half the collection angle of the objective lens:

$$\begin{aligned}\int_0^{2\pi}\int_0^{\alpha} G_{xx}^2 \sin\theta d\theta d\phi &= \int_0^{2\pi}\int_0^{\alpha} \left[\sin^2\phi + \cos^2\phi\cos\theta\right]^2 \sin\theta d\theta d\phi \\ &= \frac{\pi}{8}(1-\cos\alpha)\left(4\cos\alpha + \cos 2\alpha + 11\right) \equiv C_1\end{aligned}$$

$$\begin{aligned}\int_0^{2\pi}\int_0^{\alpha} G_{xy}^2 \sin\theta d\theta d\phi &= \int_0^{2\pi}\int_0^{\alpha} \left[\sin(2\phi)(\cos\theta - 1)/2\right]^2 \sin\theta d\theta d\phi \\ &= \frac{\pi}{12}(1-\cos\alpha)^3 \equiv C_2\end{aligned}$$

$$\begin{aligned}\int_0^{2\pi}\int_0^{\alpha} G_{xz}^2 \sin\theta d\theta d\phi &= \int_0^{2\pi}\int_0^{\alpha} \left[-\sin\theta\cos\phi\right]^2 \sin\theta d\theta d\phi \\ &= \frac{\pi}{3}(1-\cos\alpha)^2\left(\cos\alpha + 2\right) \equiv C_3\end{aligned}$$

$$\begin{aligned}\int_0^{2\pi}\int_0^{\alpha} G_{yx}^2 \sin\theta d\theta d\phi &= \int_0^{2\pi}\int_0^{\alpha} \left[\sin(2\phi)(\cos\theta - 1)/2\right]^2 \sin\theta d\theta d\phi \\ &= \frac{\pi}{12}(1-\cos\alpha)^3 \equiv C_2\end{aligned}$$

$$\begin{aligned}\int_0^{2\pi}\int_0^{\alpha} G_{yy}^2 \sin\theta d\theta d\phi &= \int_0^{2\pi}\int_0^{\alpha} \left[\cos^2\phi + \sin^2\phi\cos\theta\right]^2 \sin\theta d\theta d\phi \\ &= \frac{\pi}{8}(1-\cos\alpha)\left(4\cos\alpha + \cos 2\alpha + 11\right) \equiv C_1\end{aligned}$$

$$\begin{aligned}\int_0^{2\pi}\int_0^{\alpha} G_{yz}^2 \sin\theta d\theta d\phi &= \int_0^{2\pi}\int_0^{\alpha} \left[-\sin\theta\sin\phi\right]^2 \sin\theta d\theta d\phi \\ &= \frac{\pi}{3}(1-\cos\alpha)^2\left(\cos\alpha + 2\right) \equiv C_3\end{aligned}$$

$$\begin{aligned}\int_0^{2\pi}\int_0^{\alpha} G_{xy}G_{yx} \sin\theta d\theta d\phi &= \int_0^{2\pi}\int_0^{\alpha} \left[\sin(2\phi)(\cos\theta - 1)/2\right]^2 \sin\theta d\theta d\phi \\ &= \frac{\pi}{12}(1-\cos\alpha)^3 \equiv C_2\end{aligned}$$

$$\begin{aligned}\int_0^{2\pi}\int_0^{\alpha} G_{xx}G_{yy} \sin\theta d\theta d\phi &= \int_0^{2\pi}\int_0^{\alpha} \left[\sin^2\phi + \cos^2\phi\cos\theta\right]\left[\cos^2\phi + \sin^2\phi\cos\theta\right] \sin\theta d\theta d\phi \\ &= \frac{\pi}{24}(1-\cos\alpha)\left(20\cos\alpha + \cos 2\alpha + 27\right) \equiv C_4\end{aligned}$$

All other products of the form $G_{ij}G_{kl}$ where $i, j, k, l = \{x, y, z\}$ that are not mentioned in this list integrate to zero.

### S6 Avoiding instantaneous field-of-view (IFOV) errors

#### S6.1 Efficient sampling

To determine efficient sampling we used an approach described by Tyo *et al* [13] for interpreting the spatial sampling effect of micropolarizer arrays. The method is based on the fact that the Fourier transform of a polarization camera image has three components in the Fourier domain; $S_0$ in the center, and $S_1 + S_2$ and $S_1 - S_2$ centered at the Nyquist frequency along the x- and y-direction respectively.

The method can be used for Stokes parameter estimation as illustrated in figure S20a. In order for this method to work, the three components should not overlap in Fourier space. Figure S20b shows the effect of virtual pixel size on the overlap of the different components. The components will overlap when the pixel size is too large (undersampling).

Figures S20c-e show the optimal virtual pixel size as a function of emission wavelength for different NA oil-immersion objectives and sample media. To generate these curves, we defined the optimal pixel size as the pixel size where the components in Fourier space nearly touch. To generate 1 point on the curves in figures S20c-e, polarization camera images of a single immobilized emitter at different virtual pixel sizes (from 30 nm to 80 nm, in 0.1 nm steps) were simulated (for a given NA and sample medium). The Fourier transform of each image was taken. Then, a fixed threshold was used on the central cross-section of the magnitude of the Fourier transform to determine whether the components overlap. The largest pixel size before overlap occurs is then the calculated optimal pixel size. The fixed parameters used in the simulations are magnification = 60, tube lens focal length = 200 mm, objective focal length = 3.333 mm, refractive index immersion medium = 1.518, and field-of-view of 51-by-51 pixels (and 101-by-101 pixels for the back focal plane simulation). The refractive index for the different sample media is 1.518 for oil, 1.33 for water, and 1.0 for air.

#### S6.2 Comparison of algorithms for Stokes parameter estimation

The performance of different demosaicking algorithms for Stokes parameter estimation was compared using simulated polarization camera images of single emitters. All of the compared methods (except the Fourier-based method) use Eq. (27) to calculate the Stokes parameter images from the estimated intensity channels.
The six algorithms that were compared are:

- *1) None (no interpolation)*: The most straightforward demosaicking method is to split the unprocessed polarization camera image into four smaller images $I_{0^\circ}, I_{90^\circ}, I_{45^\circ}$ and $I_{-45^\circ}$, each consisting only of pixels covered by a polarizer with the transmission axis at the same orientation. As this results in four images that are 2-fold smaller in each dimension, they were upsampled 2-fold to compare the results with the ground truth images.

- *2-5) Interpolation*: A common demosaicking method is to split the unprocessed image into four images $I_{0^\circ}, I_{90^\circ}, I_{45^\circ}$ and $I_{-45^\circ}$ and use interpolation to fill in pixels that are unknown. The resulting estimates have the same size as the original unprocessed image. Here, we used the MATLAB function `interp2` using interpolation methods *linear*, *cubic*, *makima* (based on a modified Akima cubic Hermite interpolation), and *spline* (cubic spline using not-a-knot end conditions) using MATLAB version R2022a.

- *6) Fourier*: The Fourier domain-based method [13] described in figure S20a. The resulting estimates have the same size as the original unprocessed image.

As a ground truth, simulated conventional 4-channel images (*i.e.* without the micropolarizer array) were used. The following simulation parameters were used: magnification = 60, tube lens focal length = 200 mm, objective focal length = 3.333 mm, refractive index immersion medium = 1.518, refractive index sample medium = 1.33 (water), and field-of-view of 22-by-22 pixels (and 101-by-101 pixels for the back focal plane simulation). No noise was added (no Poisson noise, no detector noise).

The Fourier domain-based method [13] does an excellent job at retrieving the true polarization, followed by cubic spline interpolation (Fig. S21, S22, S23 and S24). The Fourier-based method does scale less favorably with image size (Fig. S25).

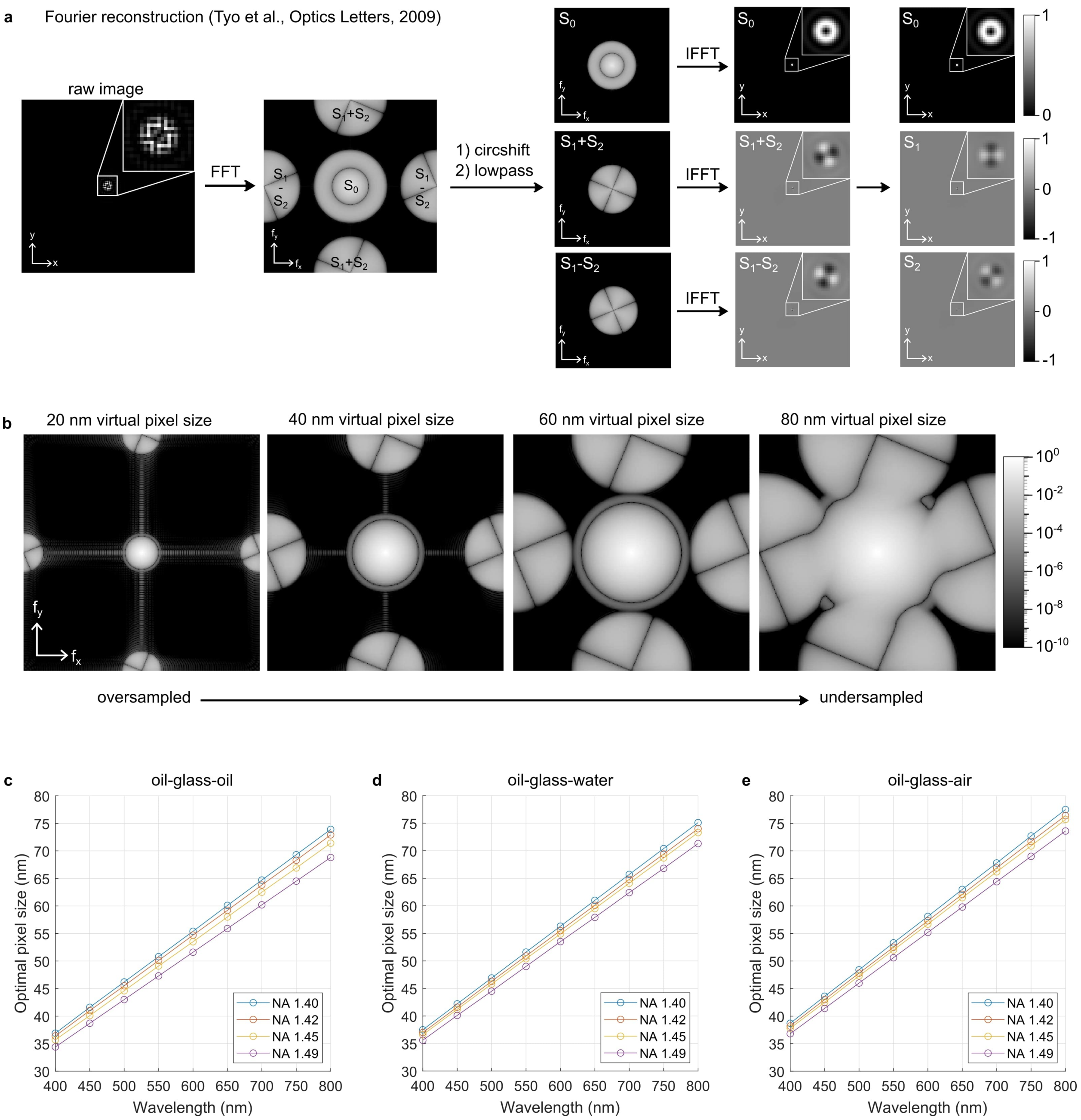

**Figure S20: Determining efficient sampling. a)** The Fourier reconstruction algorithm demonstrated on a simulated image of an immobilized single molecule ($\gamma = 1, \phi = 0, \theta = 0$). Fourier transforms are displayed as the magnitude in log scale. **b)** The magnitude of the Fourier transform of a rotationally free single emitter ($\gamma = 0$) at different virtual pixel sizes. **c)** The optimal pixel size (defined as the largest virtual pixel size that still allows accurate reconstruction using the Fourier-based reconstruction method) as a function of emission wavelength and objective NA for an oil immersion objective in the absence of a refractive index mismatch. **d)** Same as panel c, but for a sample in water. **e)** Same as panel c, but for a sample in air. The fixed parameters used in the simulations for panels c-e are magnification = 60, tube lens focal length = 200 mm, objective focal length = 3.333 mm, refractive index immersion medium = 1.518, and field-of-view of 51-by-51 pixels (and 101-by-101 pixels for the back focal plane simulation). The refractive index for the different sample media is 1.518 for oil, 1.33 for water, and 1.0 for air.

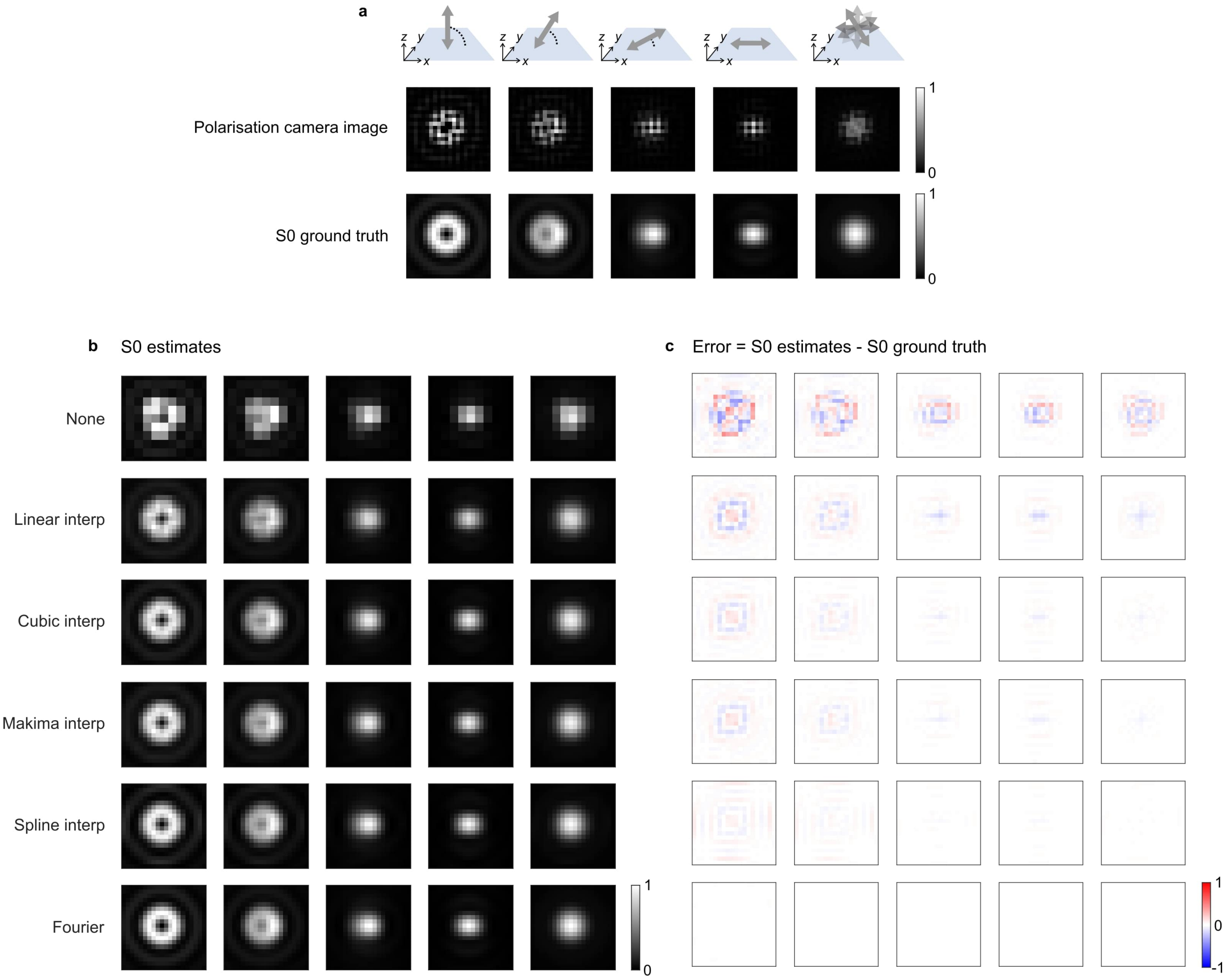

**Figure S21: Comparison of demosaicking methods on Stokes parameter $S_0$. a)** Top, diagrams of five different molecular orientations and rotational mobilities, from left to right: $(\phi, \theta, \gamma) = (0°, 0°, 1)$, $(0°, 30°, 1)$, $(0°, 60°, 1)$, $(0°, 90°, 1)$ and $(0°, 90°, 0)$. For each diagram, a simulated polarization camera image and a ground truth $S_0$ image are shown. **b)** The estimates of $S_0$ were generated using different algorithms (None, linear interpolation, cubic interpolation, makima interpolation, spline interpolation, and Fourier method). **c)** The error of the estimates generated by subtracting the estimates in panel b by the ground truth in a. The following simulation parameters were used: magnification = 60, tube lens focal length = 200 mm, objective focal length = 3.333 mm, refractive index immersion medium = 1.518, refractive index sample medium = 1.33 (water), and field-of-view of 22-by-22 pixels (and 101-by-101 pixels for the back focal plane simulation). No noise was added (no Poisson noise, no detector noise).

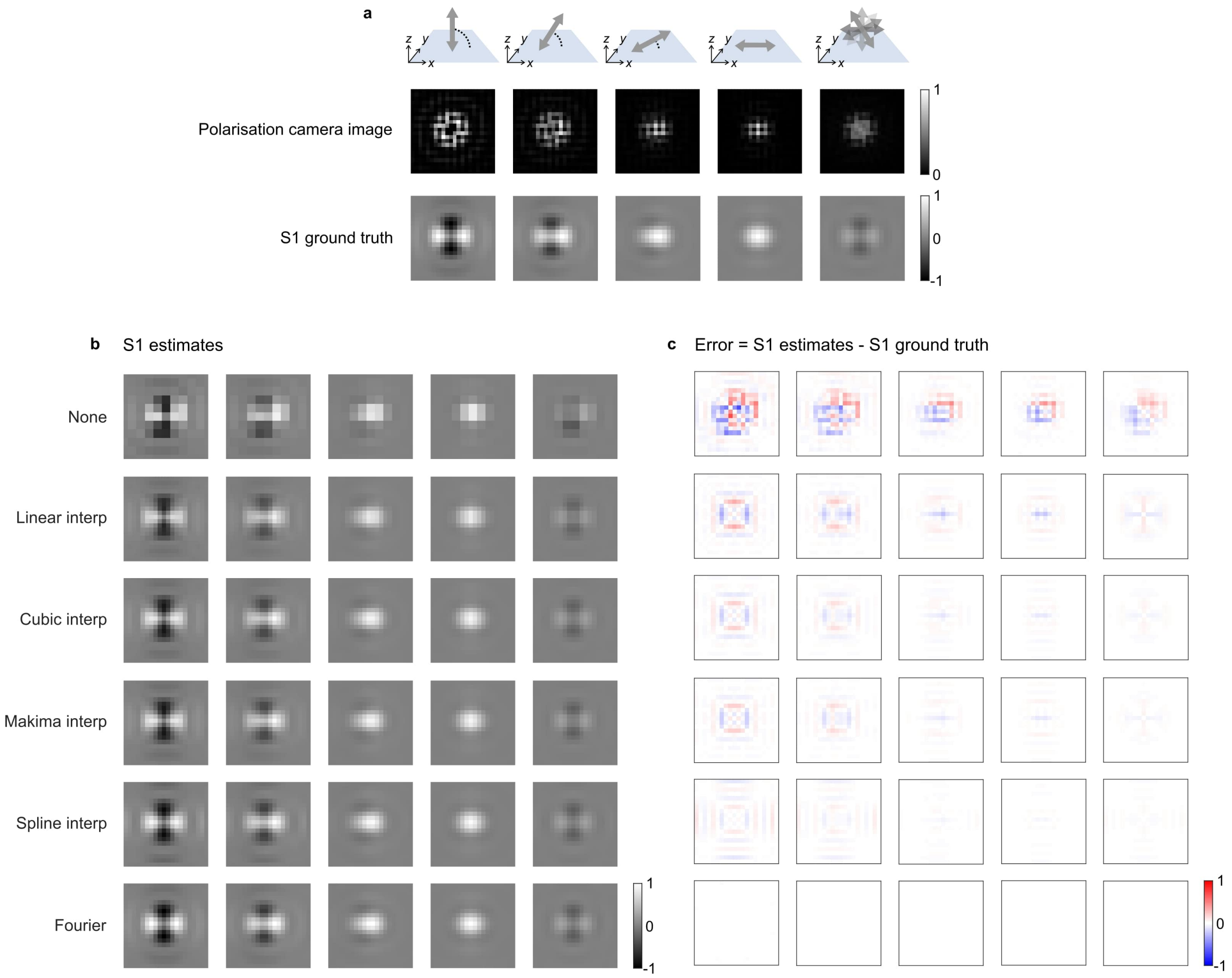

**Figure S22: Comparison of demosaicking methods on Stokes parameter $S_1$. a)** Top, diagrams of five different molecular orientations and rotational mobilities, from left to right: $(\phi, \theta, \gamma) = (0°, 0°, 1)$, $(0°, 30°, 1)$, $(0°, 60°, 1)$, $(0°, 90°, 1)$ and $(0°, 90°, 0)$. For each diagram, a simulated polarization camera image and a ground truth $S_1$ image are shown. **b)** The estimates of S1 were generated using different algorithms (None, linear interpolation, cubic interpolation, makima interpolation, spline interpolation, and Fourier method). **c)** The error of the estimates generated by subtracting the estimates in panel b by the ground truth in a. The following simulation parameters were used: magnification = 60, tube lens focal length = 200 mm, objective focal length = 3.333 mm, refractive index immersion medium = 1.518, refractive index sample medium = 1.33 (water), and field-of-view of 22-by-22 pixels (and 101-by-101 pixels for the back focal plane simulation). No noise was added (no Poisson noise, no detector noise).

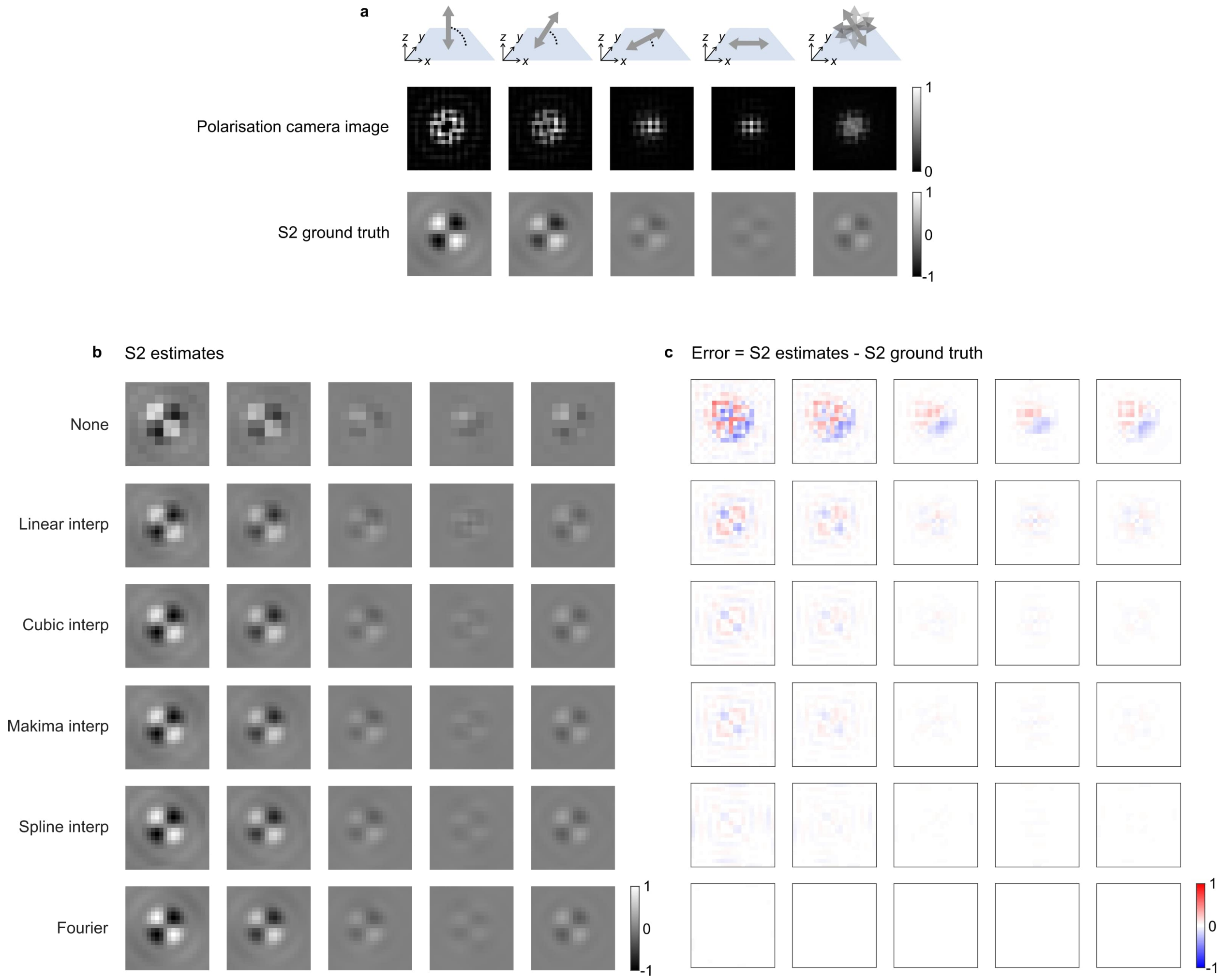

**Figure S23: Comparison of demosaicking methods on Stokes parameter $S_2$. a)** Top, diagrams of five different molecular orientations and rotational mobilities, from left to right: $(\phi, \theta, \gamma) = (0°, 0°, 1)$, $(0°, 30°, 1)$, $(0°, 60°, 1)$, $(0°, 90°, 1)$ and $(0°, 90°, 0)$. For each diagram, a simulated polarization camera image and a ground truth $S_2$ image are shown. **b)** The estimates of $S_2$ were generated using different algorithms (None, linear interpolation, cubic interpolation, makima interpolation, spline interpolation, and Fourier method). **c)** The error of the estimates generated by subtracting the estimates in panel b by the ground truth in a. The following simulation parameters were used: magnification = 60, tube lens focal length = 200 mm, objective focal length = 3.333 mm, refractive index immersion medium = 1.518, refractive index sample medium = 1.33 (water), and field-of-view of 22-by-22 pixels (and 101-by-101 pixels for the back focal plane simulation). No noise was added (no Poisson noise, no detector noise).

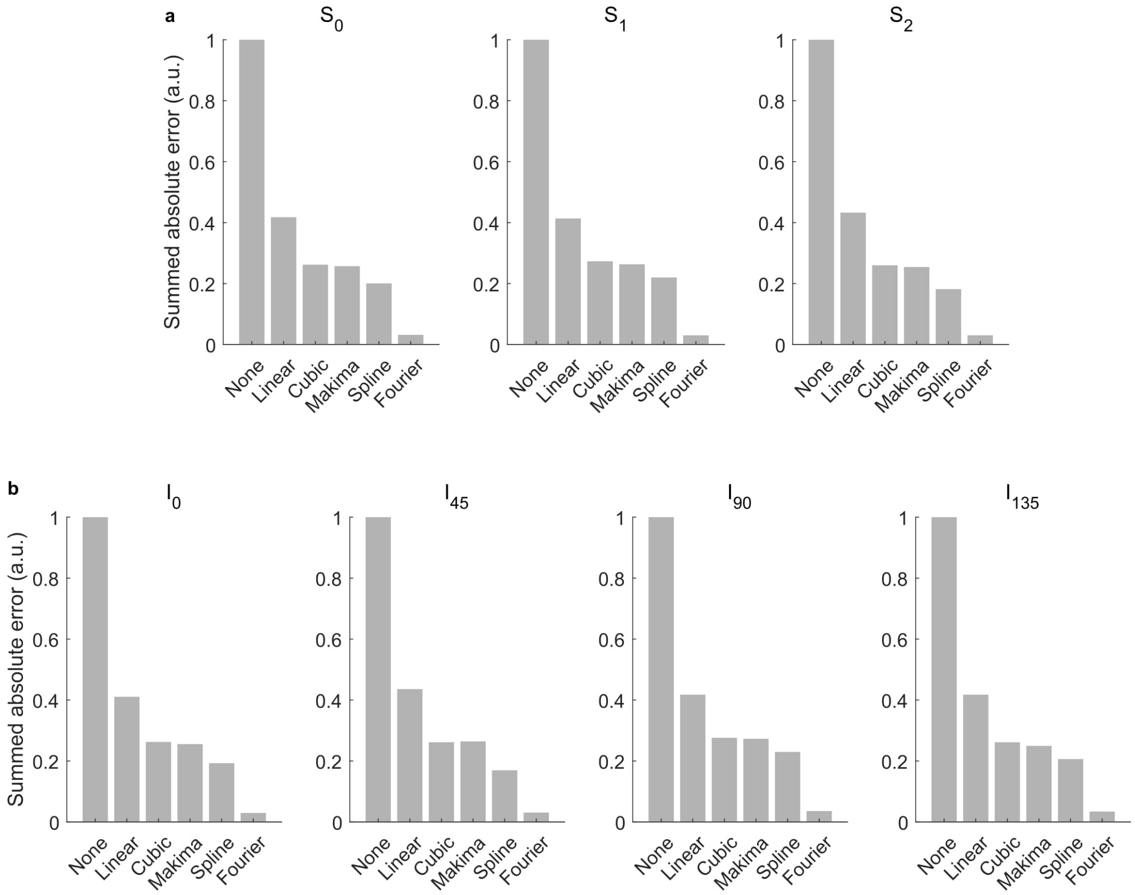

**Figure S24: Comparison of the error of different demosaicking methods. a)** The sum of the absolute value of the bias images shown in Fig. S21- S23. All results are normalized with respect to the worst performing demosaicking method (*i.e.* no interpolation or *None*). **b)** The equivalent of the plots in panel a, but for the four intensity channels instead of the Stokes parameters.

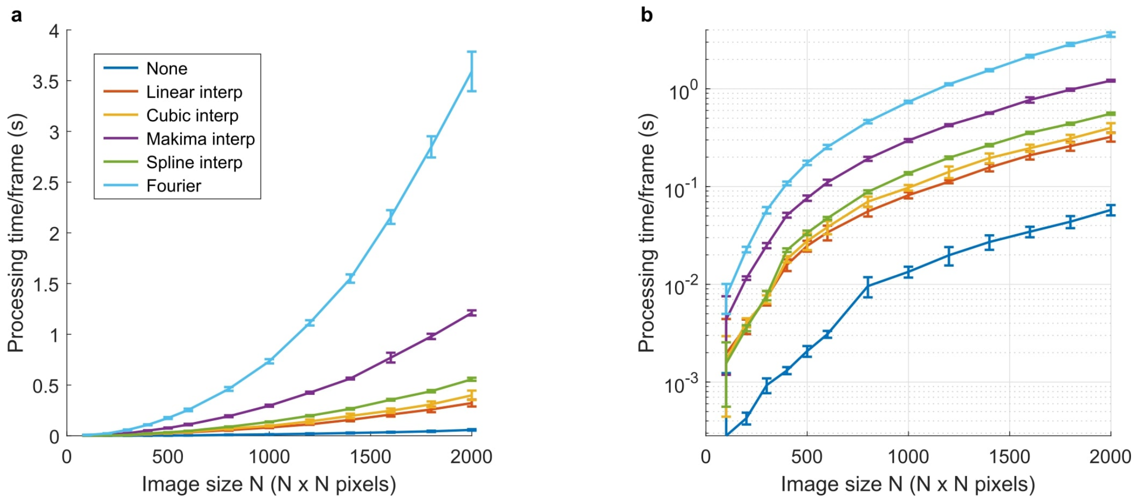

**Figure S25: Scaling of the processing speed of different demosaicking methods with image size. a)** The processing time of different demosaicking methods versus image size. The error bars are the standard deviation on 100 repeated timing measurements. **b)** The same figure as in panel a but with a logarithmic y-axis.

### S7 Single-molecule data analysis

The following describes the image analysis pipeline used for orientation estimation of all single-molecule data presented in this work. All steps in this pipeline are implemented in the form of a MATLAB application POLCAM-SR (Supplementary Fig. S43- S46).

#### S7.1 Pre-processing

##### S7.1.1 Background estimation

A spatially varying and polarized background is estimated by generating a temporal average intensity projection of the stack. The result is split into the four polarized channels. Each channel is then filtered using a median and/or gaussian filter of a specified size. Next, the smoothed channels are combined again using the original mosaic tiling of the pixels. The resulting background estimate can be subtracted from an unprocessed image.

If during image acquisition, the dataset is split into multiple substacks, the background estimation is performed separately on each substack, resulting in a spatially varying background estimation.

##### S7.1.2 Pixel-dependent camera offset and gain correction

The estimated background is subtracted from raw images. The result is then converted to units of photons by subtracting the (pixel-dependent) camera offset, and dividing by the (pixel-dependent) camera gain and quantum efficiency:

$$I_p = \frac{I_c - b - O}{g \cdot \text{QE}} \tag{42}$$

where $I_p$ is the image in units of photons, $I_c$ is the image in ADU counts, $b$ the background in ADU counts, $O$ the camera offset in ADU counts, $g$ the gain in ADU counts/photoelectron, and QE the quantum efficiency in photoelectrons/photon.

#### S7.2 Localization and orientation estimation

##### S7.2.1 Stokes estimation

Stokes estimation is performed using cubic spline interpolation or a Fourier-based method, both of which are explained in detail in section S6.2. From the estimated Stokes images ($S_0$, $S_1$, $S_2$), the angle of linear polarization (AoLP) and degree of linear polarization (DoLP) images are calculated using:

$$\text{AoLP} = \frac{1}{2}\tan^{-1}\left(\frac{S_2}{S_1}\right), \qquad \in [-\pi/2, \pi/2] \tag{43}$$

$$\text{DoLP} = \sqrt{\frac{S_1^2 + S_2^2}{S_0^2}}, \qquad \in [0, 1] \tag{44}$$

For completely linearly polarized light DoLP $= 1$ and for unpolarized light DoLP $= 0$.

##### S7.2.2 Localization

Localization is performed on the Stokes parameter image $S_0$, as it is an estimate of the intensity. Localization is performed using least squares fitting of a rotated asymmetric Gaussian. From the fitted parameters, other properties are calculated such as the intensity $I = 2\pi A\sigma_x\sigma_y$ in photons, and an estimate of the localization uncertainty using the following formula:

$$\sigma_{xy} = \sqrt{\frac{s^2 + (a^2/12)}{N} + \frac{8\pi s^4 b^2}{(Na)^2}} \tag{45}$$

where $s$ is the standard deviation of the fitted Gaussian (in nm), $a$ is the virtual pixel size (in nm), $N$ the numbers of detected signal photons (without the background) and $b$ is the constant background/pixel in photons. Equation (45) was derived by Thompson *et al.* [14] assuming an isotropic emitter, so in this work, we only use it as a rough estimate.

##### S7.2.3 Orientation estimation

For each localization, the orientation is estimated by making local measurements around the $x, y$ coordinates in the Stokes parameter images ($S_0$, $S_1$ and $S_2$), DoLP, and AoLP images.

The in-plane angle $\phi$ is calculated as a weighted average of AoLP in a $n \times n$ pixel region ($n = 5$ in this work) around the localization:

$$\phi = \sum_i w_i \text{AoLP}_i \tag{46}$$

where $w$ are weights based on the intensity image ($S_0$) and for which $\sum_i w_i = 1$. The fact that AoLP is defined on the periodic interval $[-\pi/2, \pi/2]$ is taken into account when averaging.

The out-of-plane angle $\theta$ is calculated using the following equation

$$\theta = \sin^{-1}\left(\sqrt{\frac{A \cdot \text{netDoLP}}{C - B \cdot \text{netDoLP}}}\right) \tag{47}$$

where $A$, $B$, and $C$ are constants that are a function of the half maximum collection angle of the objective $\alpha$ as defined in Eq. 31. The net degree of linear polarization (netDoLP) is calculated as:

$$\text{netDoLP} = \sqrt{\frac{\langle S_1\rangle^2 + \langle S_2\rangle^2}{\langle S_0\rangle^2}} \tag{48}$$

where the brackets $\langle ... \rangle$ refer to a weighted averaging in a $m \times m$ pixel region ($m = 13$ in this work) around the localization:

$$\langle S_0\rangle = \sum_i M_i S_{0,i} \tag{49}$$

$$\langle S_1\rangle = \sum_i M_i S_{1,i} \tag{50}$$

$$\langle S_2\rangle = \sum_i M_i S_{2,i} \tag{51}$$

where $M$ is a binary mask that gives a zero-weight to the corners in the $m \times m$ pixel region, and for which $\sum_i M_i = 1$. For netDoLP estimation, a value of $m = 15$ was used, motivated on simulations. For the specific optical setup used in this work, this corresponds to the area on the detector that just about contains the image of a single emitter with a dipole moment oriented parallel to the optical axis (i.e., the orientation with the largest spatial footprint).

As a proxy for rotational mobility, we use a weighted average of the degree of linear polarization (avgDoLP) in a $n \times n$ pixel region ($n = 5$ in this work) around the localization:

$$\text{avgDoLP} = \sum_i w_i \text{DoLP}_i \tag{52}$$

where $w$ are weights based on the intensity image ($S_0$) and for which $\sum_i w_i = 1$. The relationship between avgDoLP and the rotational mobility parameter $\gamma$ is numerically explored in Fig. S27.

#### S7.3 Post-processing: Filtering and drift-correction

The following default filtering is performed on all datasets:

- All localizations where 500 or fewer photons were detected. This threshold is motivated by simulations, as this is the point where orientation estimation breaks down (at typical experimental background levels).
- All localizations with a width that is too far away from the expected width ($\lambda/2$NA) are also discarded. This removes poor fits and out-of-focus localizations.

Drift-correction is performed using fiducial correction (Tetraspeck beads in the sample) or using RCC (redundant cross-correlation [15]).

#### S7.4 Rendering $\phi$-colorcoded single-molecule data

When rendering $\phi$-colorcoded single-molecule data, points with a low average degree of linear polarization (avgDoLP) —*i.e.* localizations with high degree of rotational mobility— and a low net degree of linear polarization (netDoLP) —*i.e.* localizations with a large out-of-plane component— are filtered out as in these cases the in-plane component of the dipole moment is small and $\phi$ becomes difficult to estimate or is meaningless (*e.g.* a rapidly rotating emitter or immobilized emitter oriented perpendicular to the sample plane).

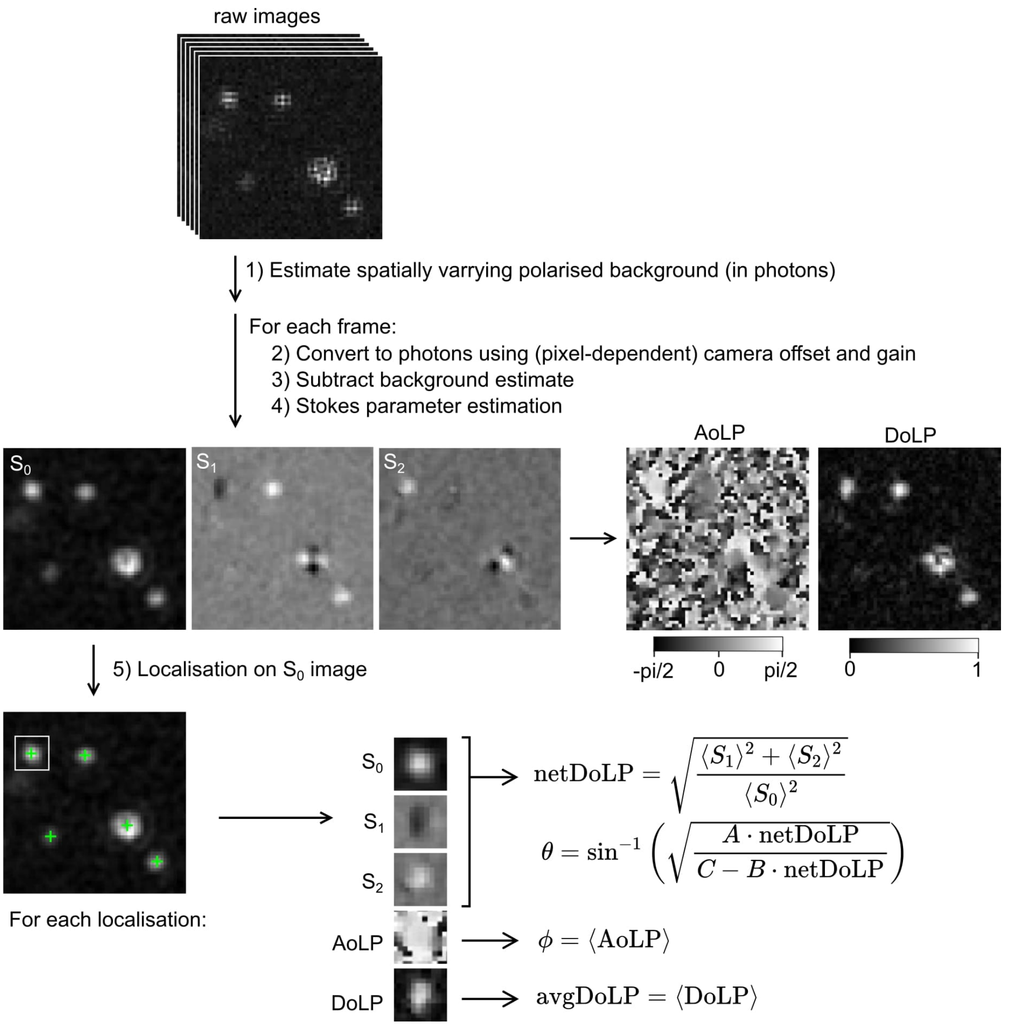

**Figure S26: Single-molecule data analysis pipeline.** From top to bottom, a background is estimated from the raw data and subtracted from all frames. The background-corrected images are converted to units of photons using (pixel-dependent) camera parameters. The resulting images are converted to Stokes parameters images ($S_0$, $S_1$ and $S_2$), from which the angle of linear polarization (AoLP) and degree of linear polarization (DoLP) images can be calculated. Localization is performed on the $S_0$ images as these are estimates of the incident intensity on the micropolarizer array. Around each identified emitter, local $S_0$-weighted measurements are taken in the Stokes, AoLP, and DoLP images that can be plugged into analytical equations to generate an estimate for $\phi$, $\theta$, and a rotational mobility parameter proxy avgDoLP.

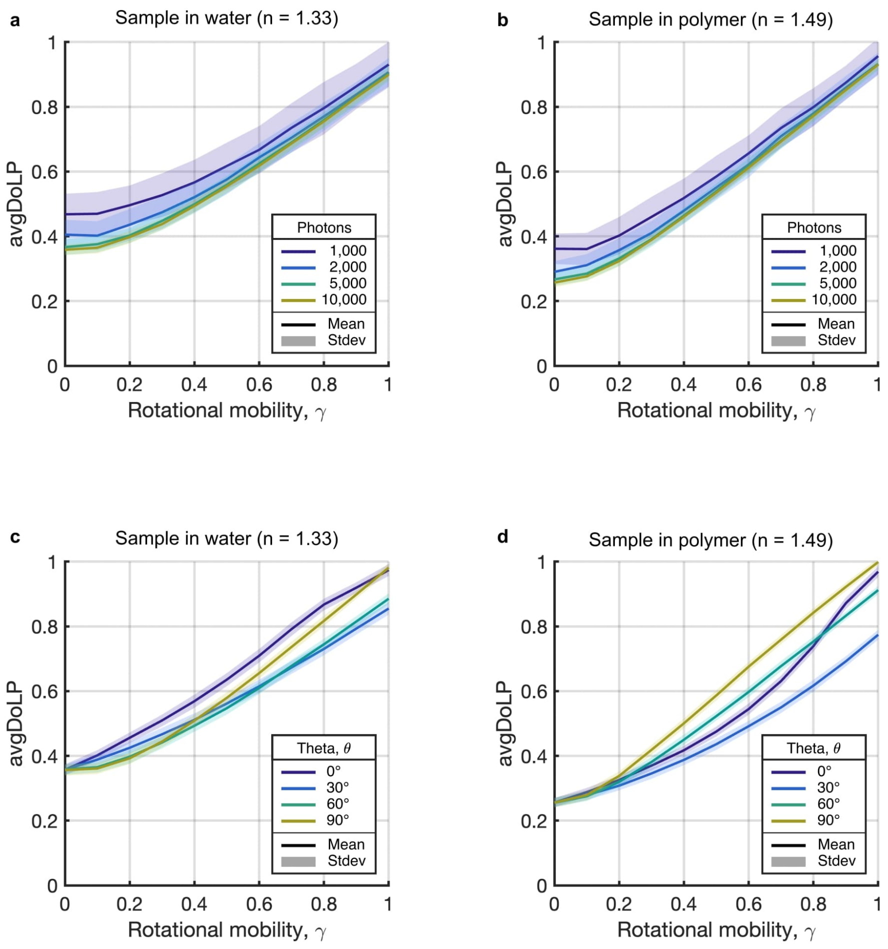

**Figure S27: Relationship between the rotational mobility parameter gamma and measured avgDoLP.** These plots were generated by estimating avgDoLP of realistic simulated images of single molecules with a known rotational mobility parameter. The simulations include realistic camera noise for the camera used in this work (CS505MUP, Thorlabs). For each image, an avgDoLP estimate was generated. For each rotational mobility value, this process was repeated (N = 200 independent noise repeats) to calculate a mean (lines) and standard deviation (envelope). The following system parameters were used: 650 nm emission wavelength, $60\times$ oil-immersion objective with a numerical aperture of 1.42, a tube lens with a focal length of 200.0 mm, physical camera pixel size of 3.45 µm. The molecules are placed on a glass-water (or glass-polymer) refractive index interface ($n_{\text{glass}} = n_{\text{oil}} = 1.518$, $n_{\text{polymer}} = 1.49$, $n_{\text{water}} = 1.33$) and in focus. A CMOS camera noise model was used (Supplementary Note S1.7). a) The measured avgDoLP versus gamma (simulation input) for various signal-to-background ratios (background photons/pixel is fixed to 5 photons), for molecules in water. Theta was fixed to 90 degrees (i.e., only in-plane emitters). b) The same as panel a, but for molecules in polymer. c) The measured avgDoLP versus gamma (simulation input) for various out-of-plane angles for molecules in water, at a high SNR (10,000 signal photons/molecule and 5 background photons/pixel). d) The same as panel c, but for molecules in polymer.

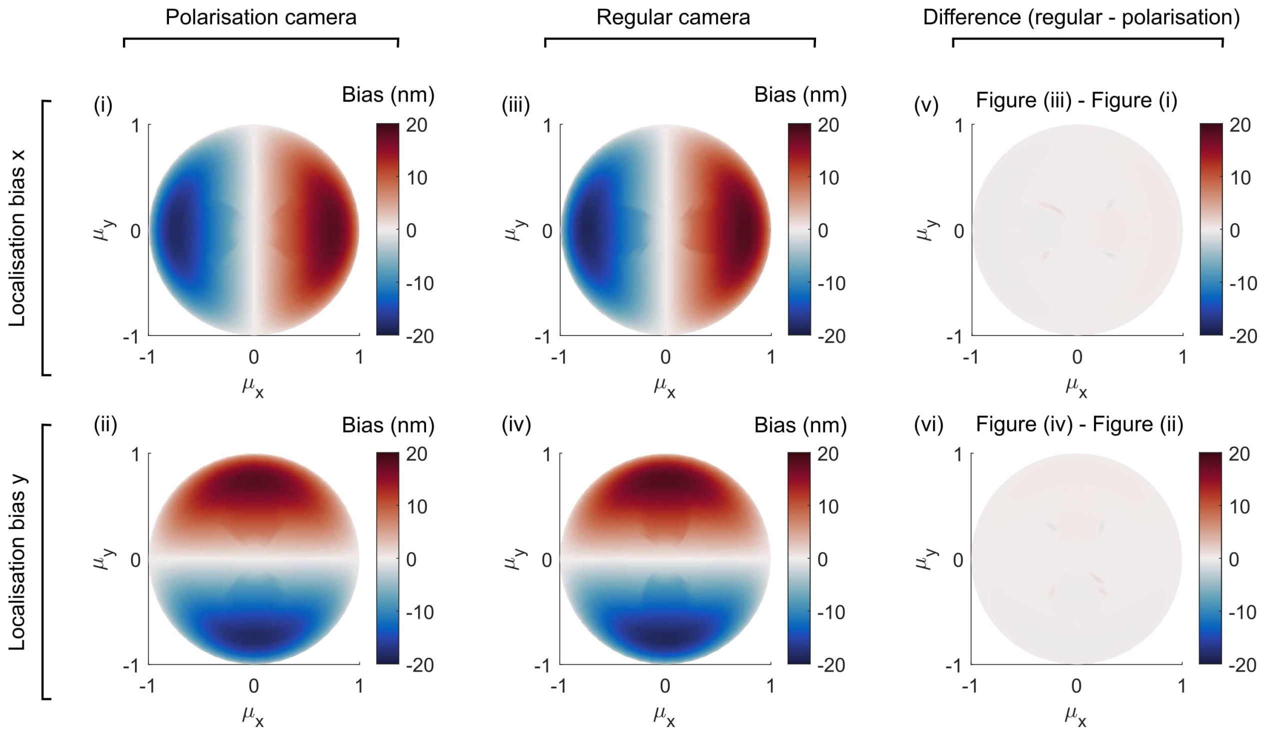

**Figure S28: Comparison of localization bias due to a dipole emitter for detection with a polarization camera and regular camera.** The localization bias due to a dipole emitter was compared between a polarization camera and a regular camera. Images of immobilized, single dipole emitters were simulated at different 3D orientations in a hemisphere ($\mu_z \geq 0$), and localized using least-squares fitting of a rotated asymmetric Gaussian. **i)** The localization bias in the x-direction, for a polarization camera. **ii)** The localization bias in the y-direction, for a polarization camera. **iii)** The localization bias in the x-direction, for a regular camera. **iv)** The localization bias in the y-direction, for a regular camera. **v)** The difference between the data in plots (iii) and (i). **vi)** The difference between the data in plots (iv) and (ii).

### S8 Experimental protocol

When performing a POLCAM experiment, the following points need to be considered:

#### S8.1 Polarization of the excitation beam at the sample plane

If the polarization of the excitation beam at the sample plane is not circularly or randomly polarized (random polarization here taken to mean *effectively unpolarized* at the timescale of the measurements), photoselection will occur; fluorescent emitters whose transition absorption dipole moment is more aligned with the dominant axis of polarization will be preferentially excited. If this photoselection effect is strong, there will be a biased detection of fluorescent emitters with certain orientations. To avoid this, it is important to characterize the polarization of the excitation beam at the sample plane. This can be done by placing a linear polarizer in the excitation beam right after the objective lens, and measuring the power of the beam after the polarizer while rotating the polarizer. In an ideal case, the power will remain constant regardless of the polarizer's orientation. In practice, it is highly likely that there will be some oscillation. We found that using a multi-mode optical fiber resulted in a more random polarization at the sample plane compared to the more typical arrangement of a linear polarizer followed by a quarter waveplate (Fig S29).

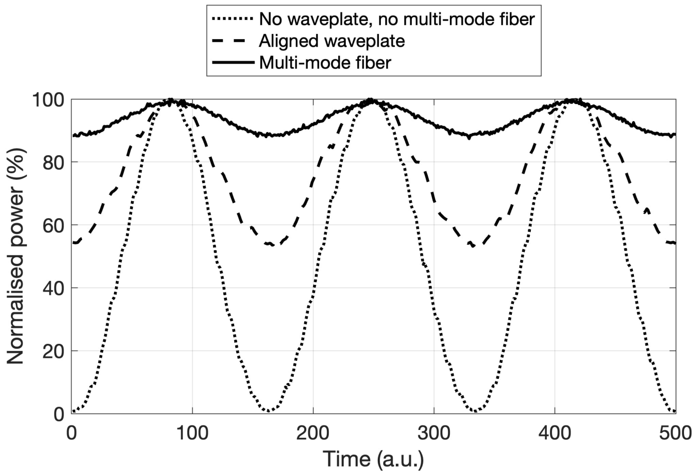

**Figure S29: Polarization of the excitation beam.** Measured power after a rotating polarizer.

During experiments performed in this work, the software POLCAM-Live (`https://github.com/ezrabru/POLCAM-Live`) was used to reconstruct data live during acquisition. If the polarization of the excitation is not circular or random, this will usually give rise to the background (and sample) looking polarized in one direction when displaying the images in *polarisation colormap* or *HSVmap* mode. This will be especially apparent when doing non-single-molecule experiments, or when imaging samples with a lot of fluorescence background. This can be corrected by tweaking the alignment of any polarization-controlling optics in the excitation path while reconstructing the images live. The alignment should be optimal when the AoLP of the background appears random, or DoLP is minimized. We found that when a multi-mode optical fiber is used for excitation, this was not necessary and excellent results could be achieved without this step. This is not necessarily true when imaging outside of EPI illumination mode (*e.g.*, HILO, or TIRF), as the polarization at the sample plane will become more challenging to control. As a result, most data was acquired using EPI illumination.

#### S8.2 Determining the orientation of micropolarizers

For correct image processing, it is important to correctly assign the orientation of the transmission axis of the micropolarizer covering each pixel. This can be performed by illuminating the camera chip with light with a known polarization (*e.g.*, incandescent light bulb or room light followed by a linear polarizer with a known transmission axis such as LPVISE100-A, Thorlabs). Coarsely sampling different polarization orientations will allow the correct assignment of the orientation of the transmission axes of the micropolarizers. Alternatively, a test sample with known polarization can be imaged and the assignment back-calculated. As long as the coordinates of the region of interest are known, this step does not have to be repeated for each experiment.

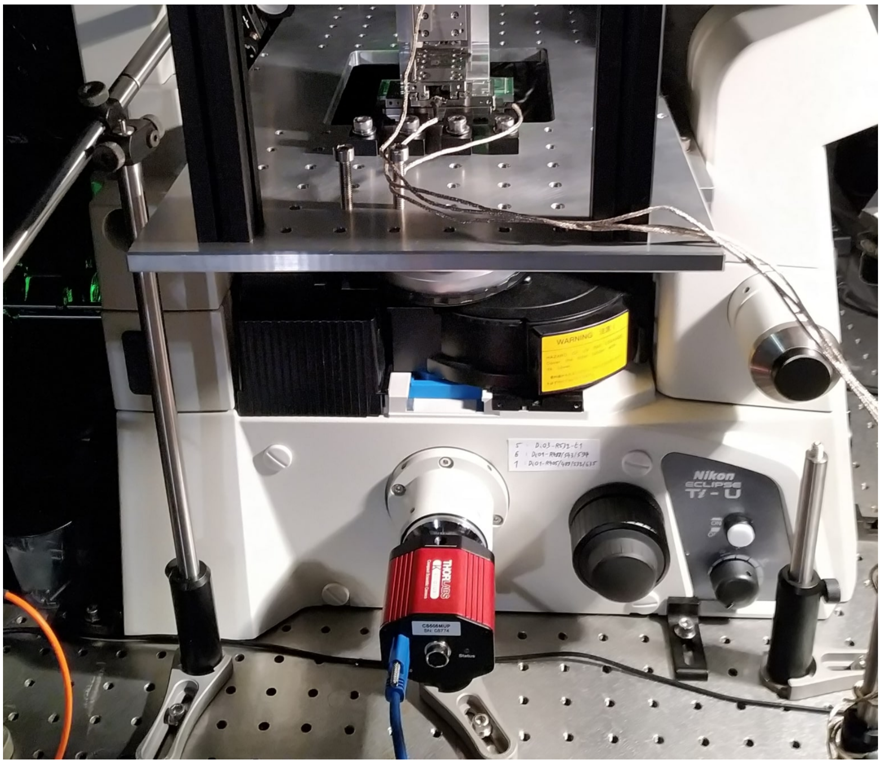

**Figure S30: Optical setup.** A picture of the optical setup. The camera is directly mounted at the camera port of a Nikon Eclipse Ti-U microscope body.

| Figure Ref. | Sample | Dye | Dichroic mirror | Emission filter(s) | Wavelength laser (nm) | Power density (kW/cm$^2$) | Exposure time (ms) |
|---|---|---|---|---|---|---|---|
| 1a-c | Dye on glass | SYTOX Orange | Di03-R532-t1 | FF01-582/64 | 520 | 0.36 | 100 |
| 1e, 6a | Lipid-bilayer bead | Di-8-ANEPPS | Di03-R514-t1 | FF01-515/LP, FF01-650/200 | 514 | 2.02 | 200 |
| 1f | Lipid-bilayer bead | NR4A | Di03-R514-t1 | FF01-515/LP, FF01-650/200 | 514 | 6.08 | 50 |
| 1g | Lipid-bilayer bead | Nile red | Di03-R514-t1 | FF01-515/LP, FF01-650/200 | 514 | 6.08 | 50 |
| 1i,j | Dye in polymer | Alexa Fluor 647 | Di03-R405/488/561/635-t1 | BLP01-635R | 638 | - | 10-2,000 |
| 2 | $\alpha$-syn fibrils | Nile red | Di03-R532-t1 | BLP01-532R, FF01-650/200 | 532 | 2.42 | 50 |
| 3a-c, 3e-g, 4 | dSTORM F-actin HeLa | Alexa Fluor 488 | Di02-R488 | BLP01-488R, FF01-582/64 | 488 | 5.04 | 30 |
| 3b, 3d-f | dSTORM F-actin HeLa | Alexa Fluor 647 | Di03-R405/488/561/635-t1 | BLP01-635R | 638 | 3.51 | 30 |
| 6c | F-actin COS7 | SiR-actin | Di03-R405/488/561/635-t1 | BLP01-635R | 638 | - | 100 |
| 6d-e | membrane live T cell | NR4A | Di03-R514-t1 | FF01-515/LP, FF01-650/200 | 514 | 0.004 | 200 |
| 6f | membrane live T cell | NR4A | Di03-R514-t1 | FF01-515/LP, FF01-650/200 | 514 | 0.004 | 200 |
| S5 | Dye on glass | SYTOX Orange | Di03-R532-t1 | FF01-582/64 | 520 | 0.36 | 100 |
| S13c-d | Lipid-bilayer bead | DiI | Di03-R514-t1 | FF01-515/LP, FF01-582/64 | 514 | 0.81 | 200 |
| S13e | Lipid-bilayer bead | Nile red | Di03-R514-t1 | FF01-515/LP, FF01-650/200 | 514 | 2.02 | 200 |
| S13f | Lipid-bilayer bead | Di-8-ANEPPS | Di03-R514-t1 | FF01-515/LP, FF01-650/200 | 514 | 2.02 | 200 |
| S13g-h | Lipid-bilayer bead | NR4A | Di03-R514-t1 | FF01-515/LP, FF01-650/200 | 514 | 6.08 | 50 |
| S14 | $\alpha$-syn fibrils | Nile red | Di03-R532-t1 | BLP01-532R, FF01-650/200 | 532 | 2.42 | 50 |
| S15 | Dye in polymer | Alexa Fluor 647 | Di03-R405/488/561/635-t1 | BLP01-635R | 638 | - | 10-2,000 |
| S21 | dSTORM F-actin HeLa | Alexa Fluor 488 | Di02-R488 | BLP01-488R, FF01-582/64 | 488 | 5.04 | 30 |
| S22 | Lipid-bilayer bead | NR4A | Di03-R514-t1 | FF01-515/LP, FF01-650/200 | 514 | 2.02 | 200 |
| S23-S25 | $\alpha$-syn fibrils | Nile red | Di03-R532-t1 | BLP01-532R, FF01-650/200 | 532 | 2.42 | 50 |

**Table 3: Imaging parameters**. Experimental parameters used for imaging. All dichroic mirrors and emission filters were bought from Semrock. If an entry in the column *Power density* is labeled '-', it means that no power density was measured during that specific experiment. The power densities were estimated at the end of experiments by removing the sample and measuring the total power of the excitation beam exiting the objective using a power meter. This value was divided by the area of the illuminated area, which was estimated from the illumination profile visible in the images themselves (which was possible as the image of the illuminated area was smaller than the camera sensor).

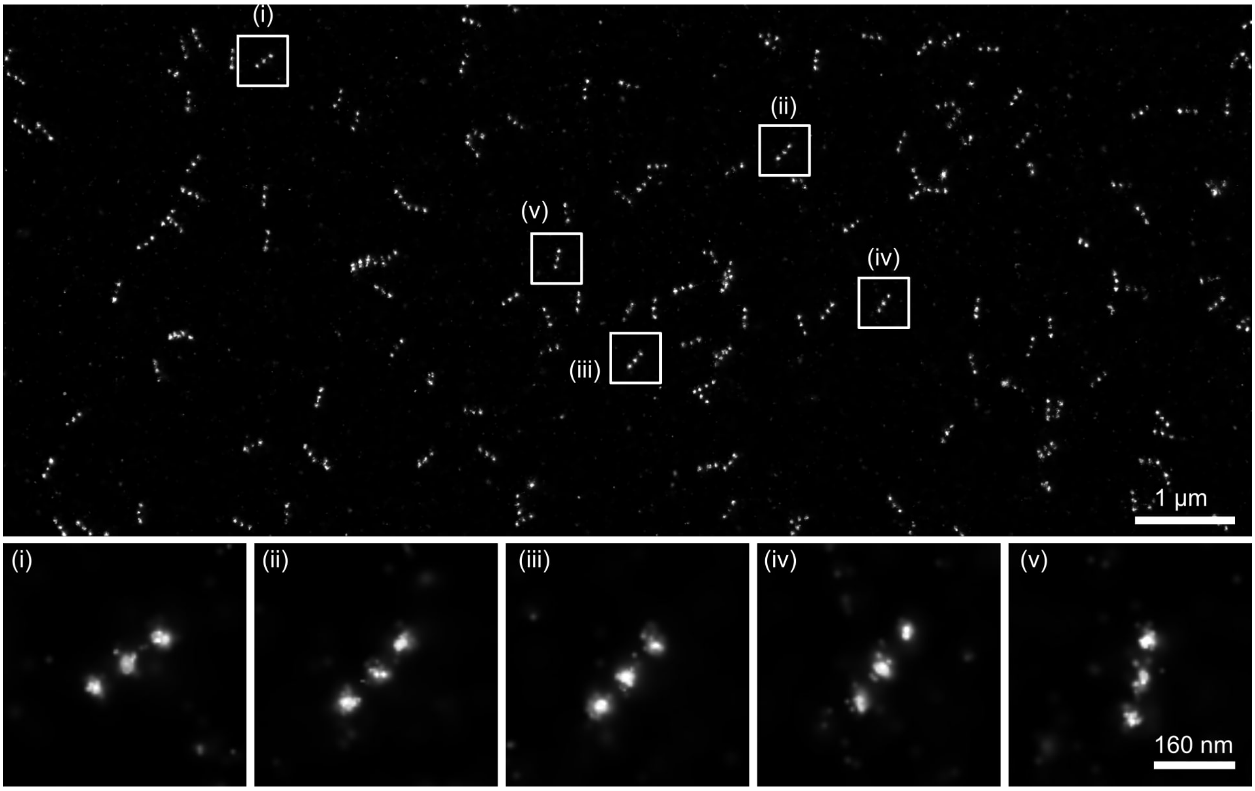

**Figure S31: DNA-PAINT of DNA-origami nanorulers nanorulers reconstructed in Picasso.** Reconstruction of DNA-origami nanorulers (GATTA-PAINT 80R, Gattaquant) that were imaged using the polarization camera (20.000 frames, 200 ms exposure time, 4.8 kW/cm$^2$ in TIRF). The nanorulers have three docking sites with a 80 nm spacing between docking sites. The imager strands are labeled with ATTO 655. Raw polarisation camera images were converted to S0 and then analysed in Picasso [16].

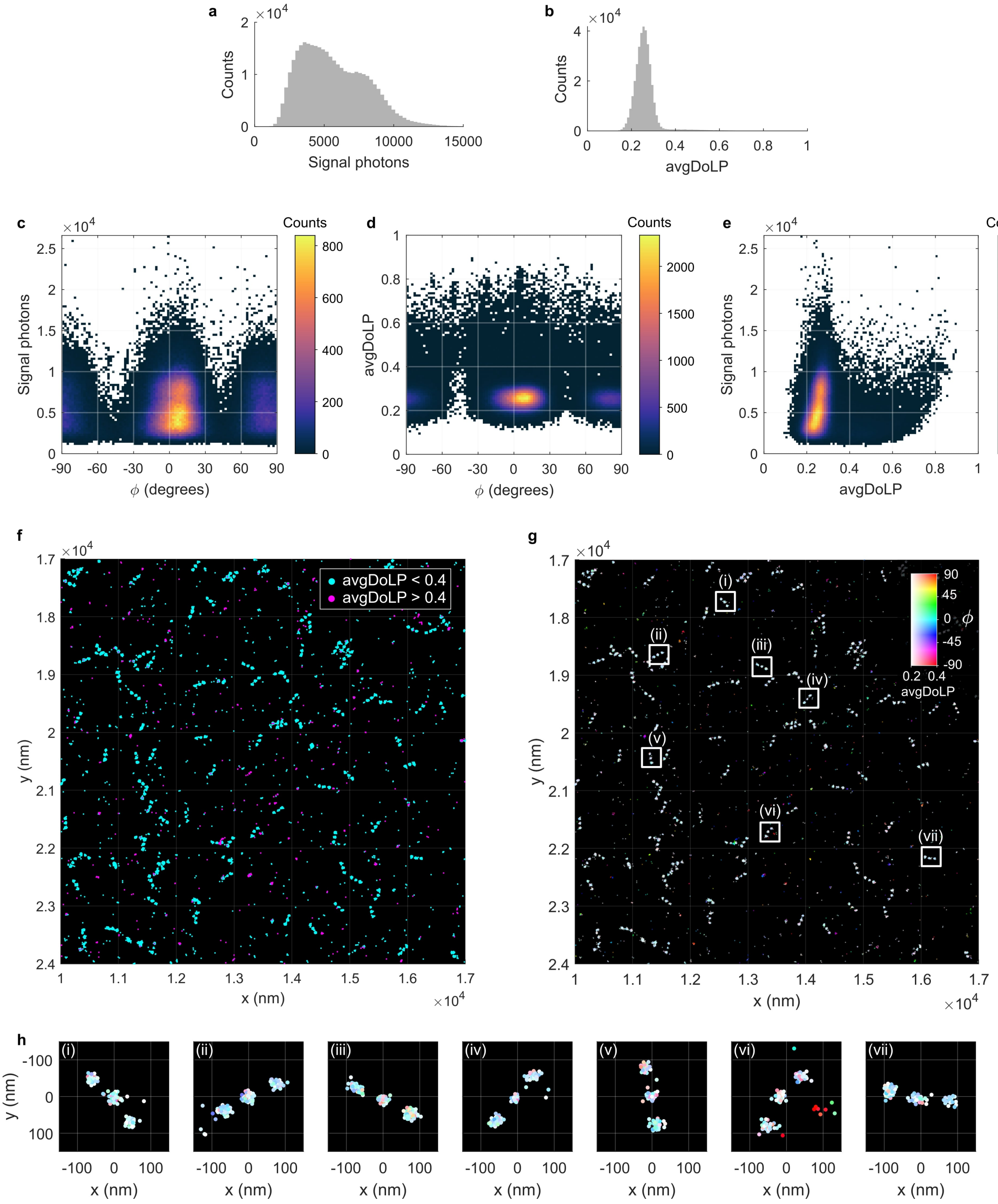

**Figure S32: DNA-PAINT of DNA-origami nanorulers reconstructed in POLCAM-SR.** Reconstruction of DNA-origami nanorulers (GATTA-PAINT 80R, Gattaquant) that were imaged using the polarization camera (20.000 frames, 200 ms exposure time, 4.8 kW/cm$^2$ in TIRF). The nanorulers have three docking sites with a 80 nm spacing between docking sites. The imager strands are labeled with ATTO 655. Raw polarisation camera images were processed using POLCAM-SR. The sample drift was corrected using RCC [15]. **a)** The distribution of detected photons per localisation/frame. **b)** The distribution of the avgDoLP of localisations. The avgDoLP is low compared to other datasets presented in this work (e.g., Fig. 3g and 4e in the main manuscript), suggesting that the dyes on the imager strands are rotationally free relative to the DNA origami structure. **c)** Heatmap of the estimated in-plane angle versus the detected photons per localisation/frame. **d)** Heatmap of the estimated in-plane angle versus avgDoLP. This distribution is characteristic of rotationally-free emitters (*i.e.*, peaks around 0, 90 and -90 degrees at low avgDoLP). **e)** Heatmap of the detected photons per localisation/frame versus avgDoLP. **f)** Scatter plot of the reconstructed DNA-origami data in two colours: All localisations with an avgDoLP larger than 0.4 are magenta, and all with an avgDoLP smaller than 0.4 are cyan. Nearly all localisations associated with origami are cyan, magenta localisations are likely probably free imager strand binding non-specifically to the surface. **g)** The same data as shown in panel f, but with a 2D colormap that encodes both the avgDoLP and the in-plane angle $\phi$. The map can be interpreted as follows: white dots have low avgDoLP, and coloured dots have high avgDoLP. If localisations in a cluster have the same colour, they are oriented in the same direction. Many non-specific binding events seem to have high avgDoLP and be oriented in the same direction, supporting the idea that these might be free dye binding to the surface. **h)** Representative examples of individual origami from panel g.

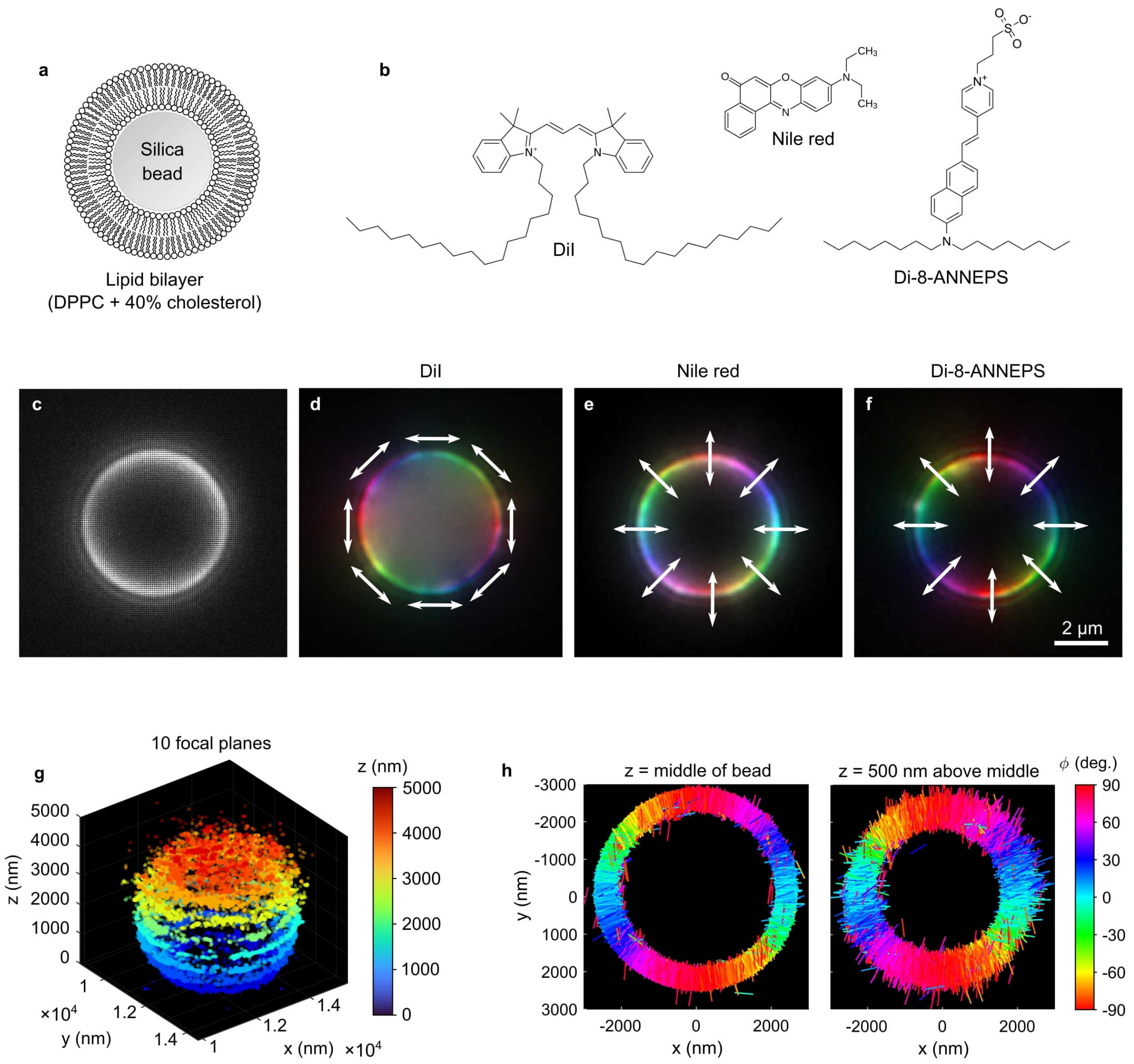

**Figure S33: Lipid-coated silica beads and lipid probes.** **a)** Diagram of a silica bead coated in a lipid bilayer. **b)** The molecular structures of three different lipid membrane dyes. **c)** Unprocessed polarization camera image of a lipid-bilayer (DPPC + 40% cholesterol) coated silica bead (5 μm diameter) labeled with DiI, focused approximately at a plane in the middle of the bead. **d)** A polarization colormap rendering of the bead is shown in panel c. **e)** A polarization colormap rendering of a bead as shown in panel c, but labeled with Nile red. **f)** A polarization colormap rendering of a bead as shown in panel c, but labeled with the dye Di-8-ANNEPS. **g)** A *semi*-3D rendering of a PAINT reconstruction of a lipid bilayer-coated bead labeled with the Nile red derivative NR4A. The data of 10 different z-planes are shown together. (The data is not truly 3D, because if the data were observed from the zy or zx plane, the point cloud would look like 10 lines.) **h)** Reconstruction of a lipid-coated bead labeled with NR4A and imaged in PAINT mode at two different focal planes. Each rod represents a detected emitter. The direction and color of each rod indicate the measured in-plane angle $\phi$.

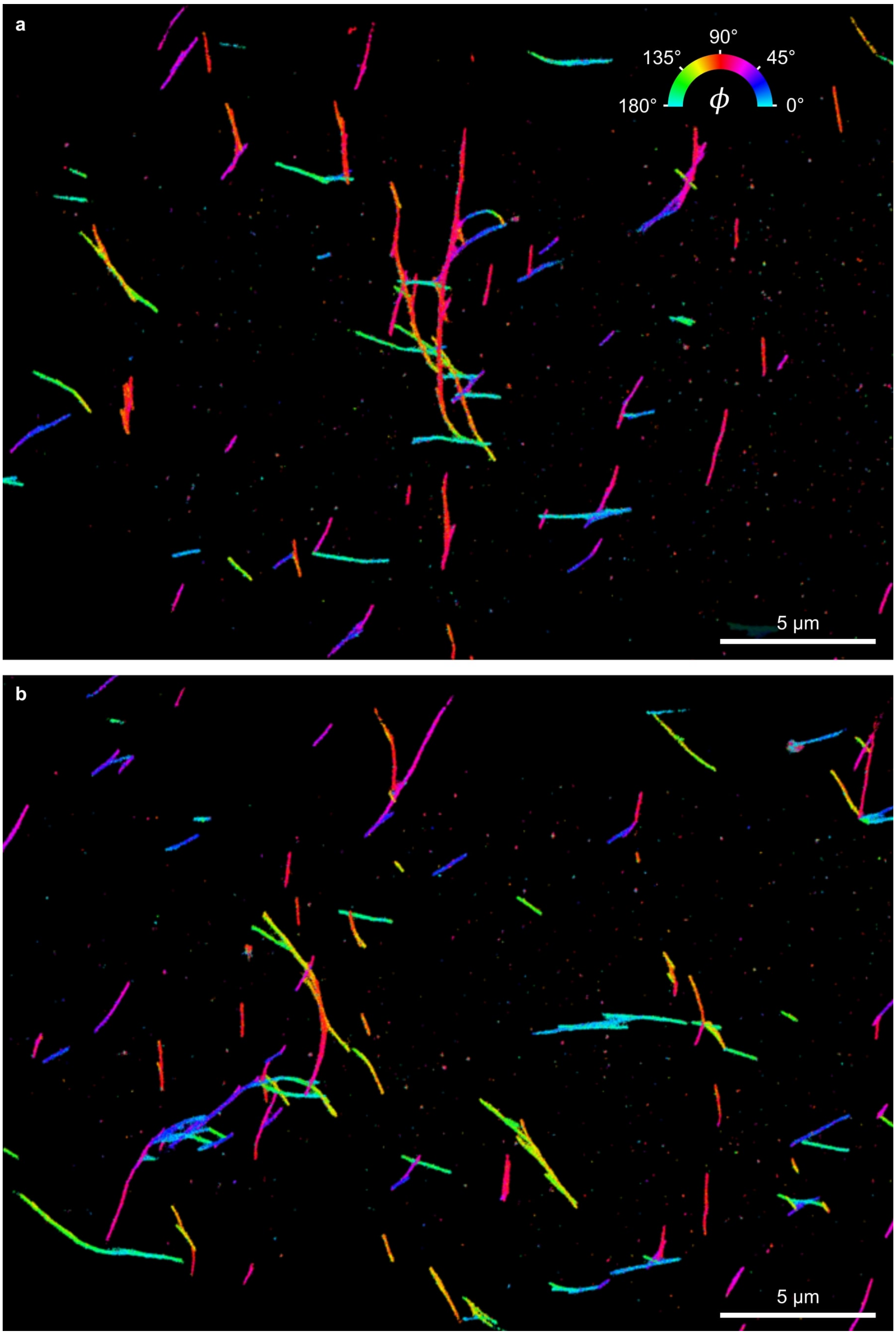

**Figure S34: Representative Nile red TAB-PAINT reconstructions of alpha-synuclein fibrils.**

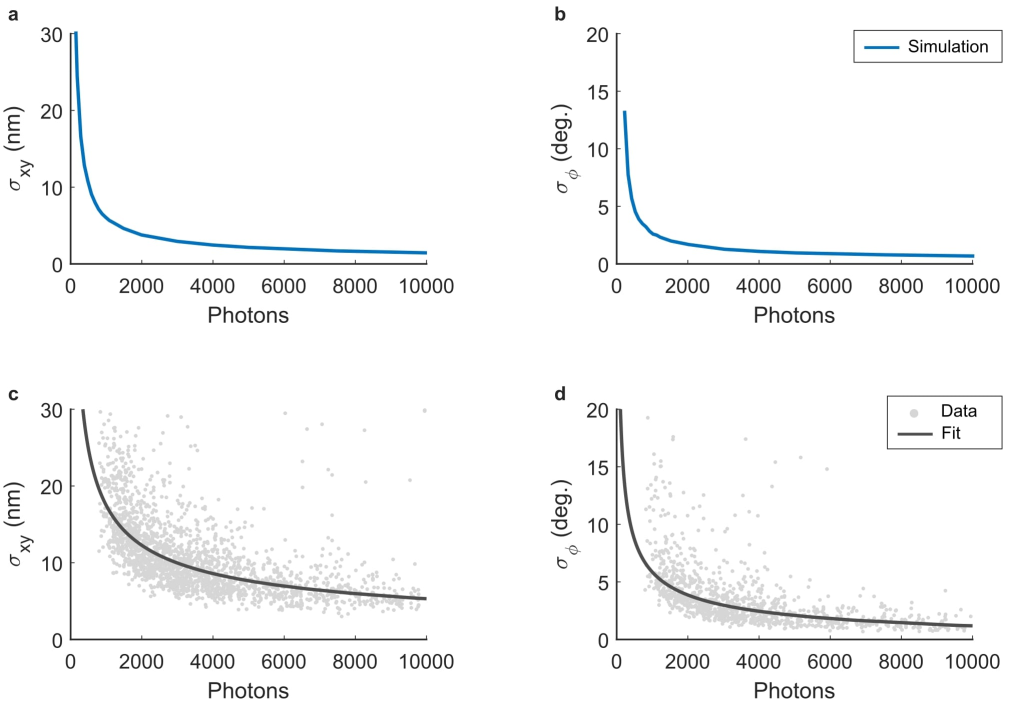

**Figure S35: Comparison of experimental precision with simulations. a)** Localization precision calculated as the standard deviation on the repeated estimate of the position of an emitter from simulated images with a realistic camera noise model. All simulation parameters were set to match the experiment presented in Figure 2i,j, *i.e.* AF647 immobilized in PVA. All molecules were assumed to be oriented parallel to the cover glass ($\theta = 90°$), have rotational mobility of $\gamma = 0.75$ (as previously measured for molecules immobilized in PVA), background of 10 photons/pixel, emission wavelength of 665 nm, sample medium PVA with a refractive index of 1.477. **b)** In-plane angle $\phi$ precision calculated as the standard deviation on the repeated estimate of $\phi$ from simulated images. The same parameters were used as described for panel a. **c)** The experimental localization precision, calculated as the standard deviation on the repeated estimate of the position of single AF6467 molecules immobilized in PVA. **d)** The experimental $\phi$ precision, calculated as the standard deviation on the repeated estimate of the in-plane angle of single AF6467 molecules immobilized in PVA. The data in panels c and d are the same data as shown in Figures 2i and 2j in the main manuscript.

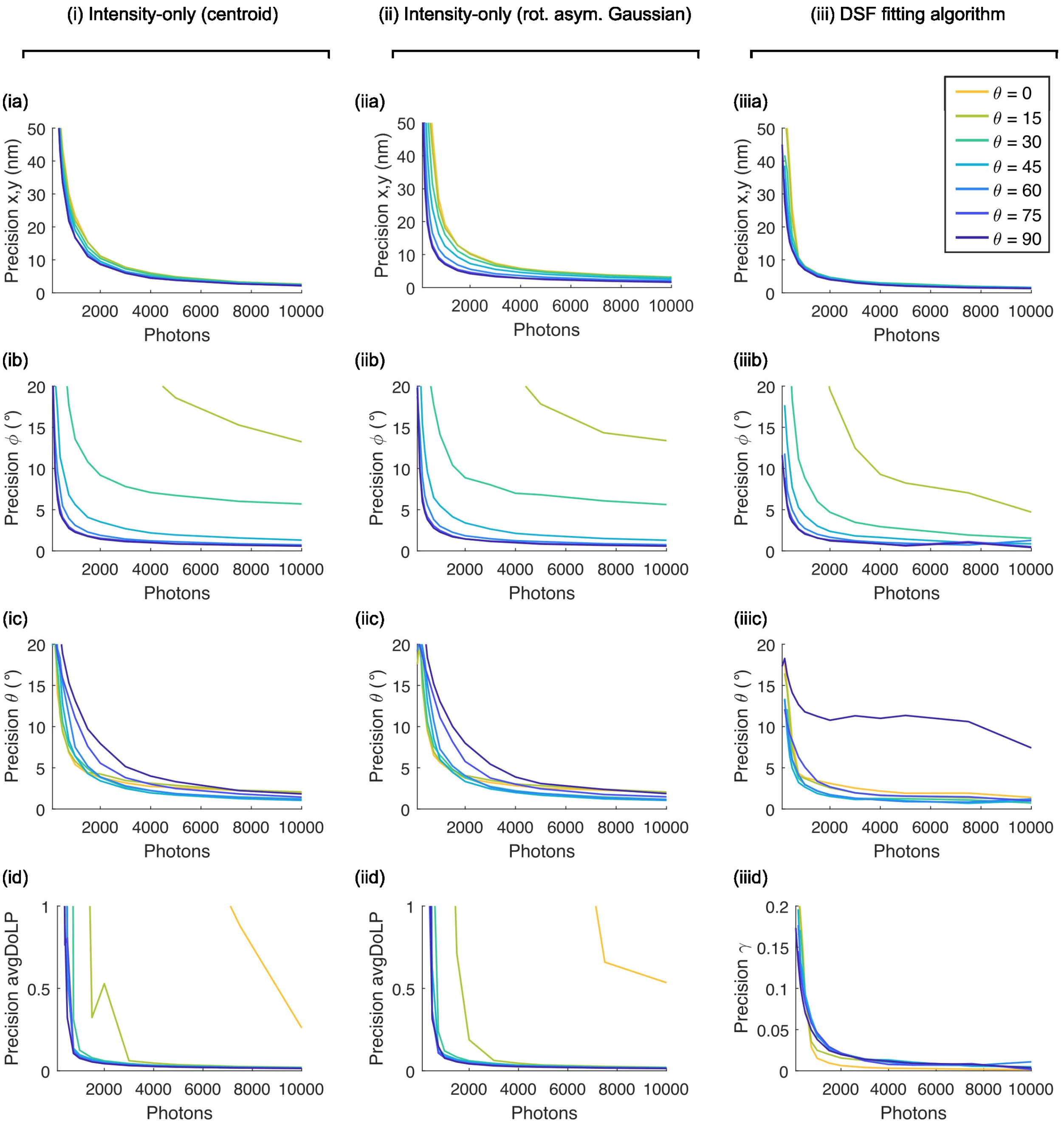

**Figure S36: Precision as a function of photon number (10 bkgnd photons/pixels, $\gamma = 1$).** The precision (standard deviation on a repeated estimate) of different parameters as a function of the detected number of photons. Each plot contains multiple lines, representing a set of simulations at a fixed $\theta$. The plots are organized in three columns, each representing the results for a different algorithm: (i) intensity-only algorithm using centroid localization, (ii) intensity-only algorithm using least squares fitting of a 2D rotated asymmetric Gaussian and (iii) DSF-fitting algorithm.

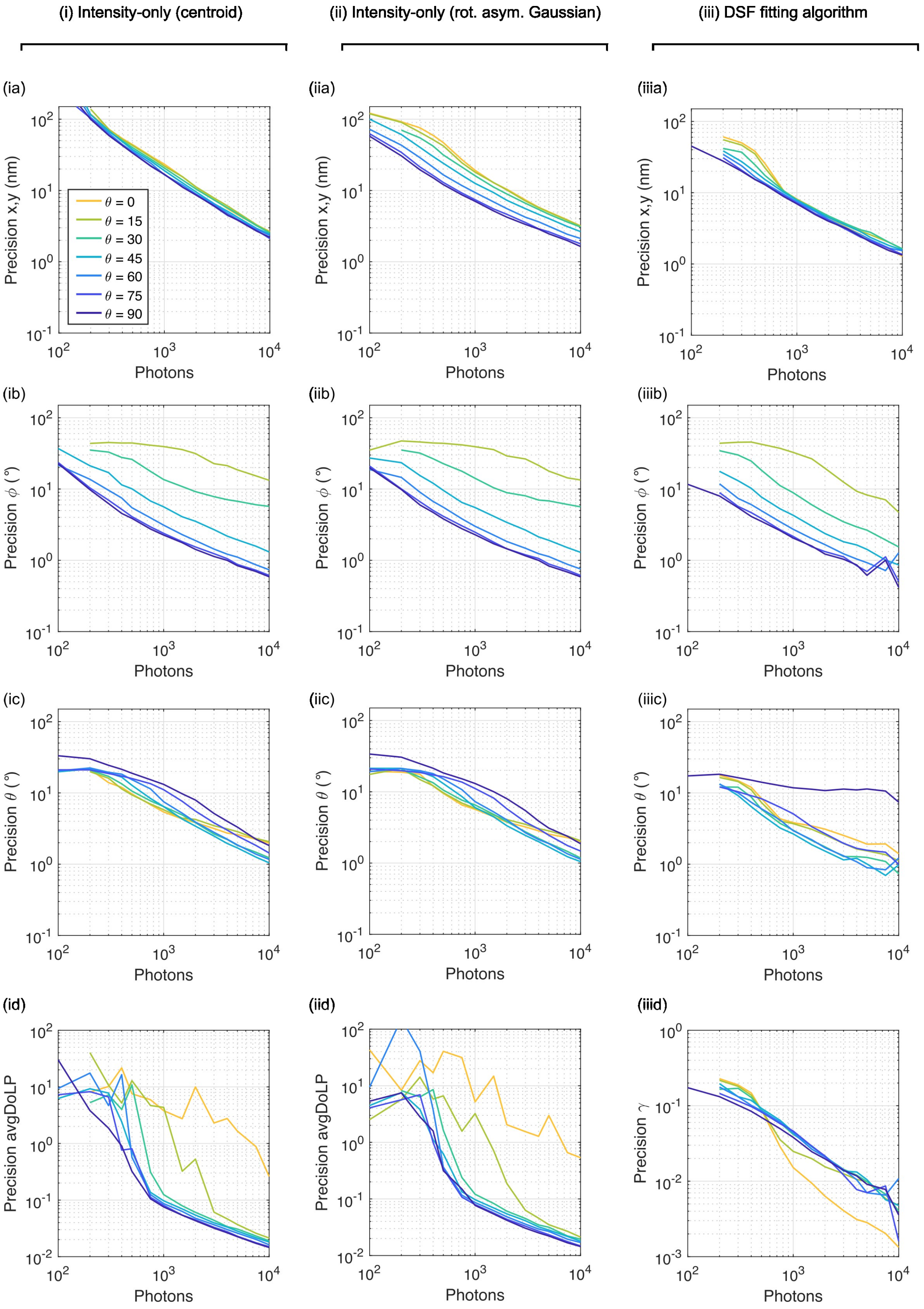

**Figure S37: Precision as a function of photon number (10 bkgnd photons/pixels, $\gamma = 1$).** The same data as shown in figure S36, but using a log-scale. The precision (standard deviation on a repeated estimate) of different parameters as a function of the detected number of photons. Each plot contains multiple lines, representing a set of simulations at a fixed $\theta$. The plots are organized in three columns, each representing the results for a different algorithm: (i) intensity-only algorithm using centroid localization, (ii) intensity-only algorithm using least squares fitting of a 2D rotated asymmetric Gaussian and (iii) DSF-fitting algorithm.

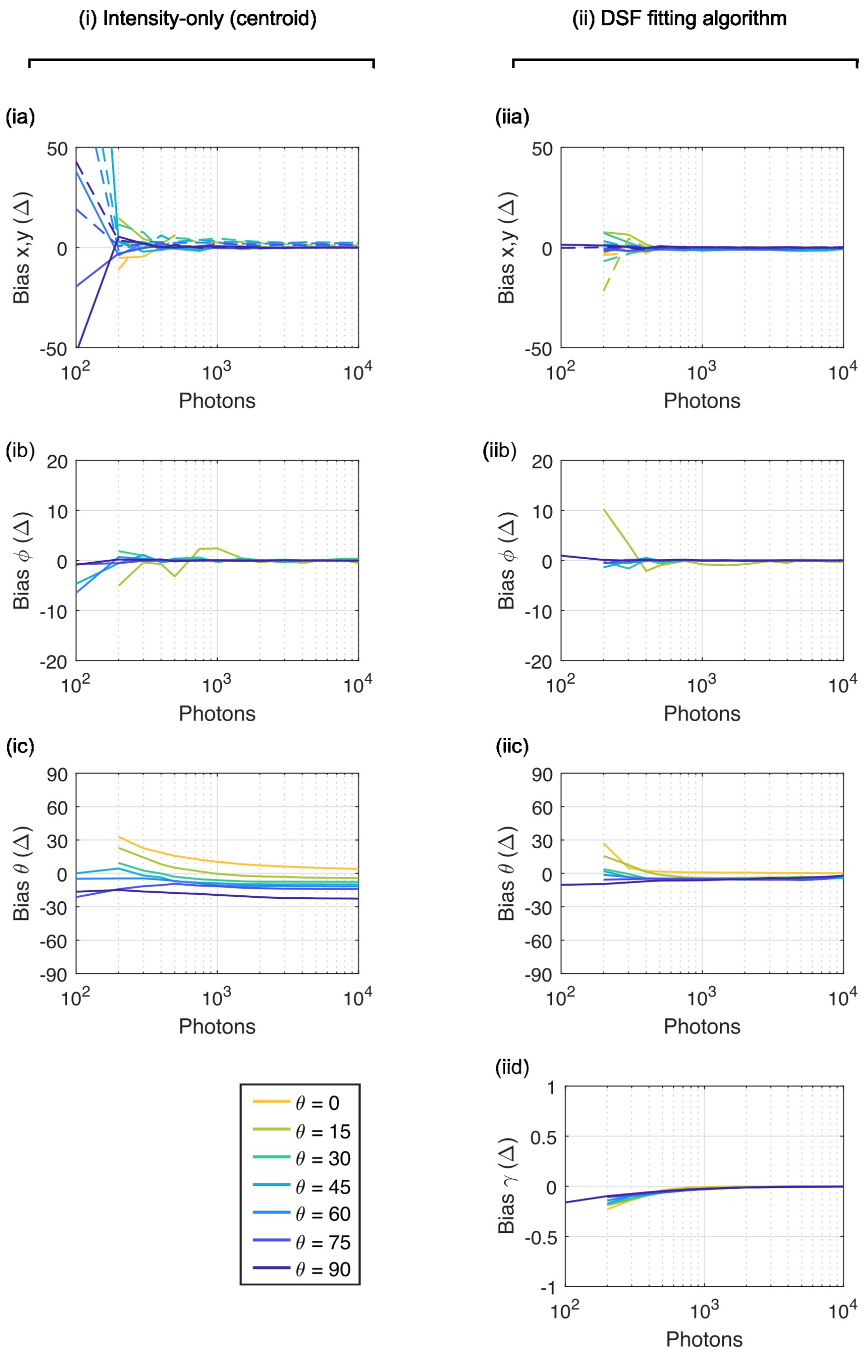

**Figure S38: Bias as a function of photon number (10 bkgnd photons/pixels, $\gamma = 1$).** The bias (difference between the true value and estimated value) of different parameters as a function of the detected number of photons. Each plot contains multiple lines, representing a set of simulations at a fixed $\theta$. The plots are organized in three columns, each representing the results for a different algorithm: (i) intensity-only algorithm using centroid localization, (ii) intensity-only algorithm using least squares fitting of a 2D rotated asymmetric Gaussian and (iii) DSF-fitting algorithm.

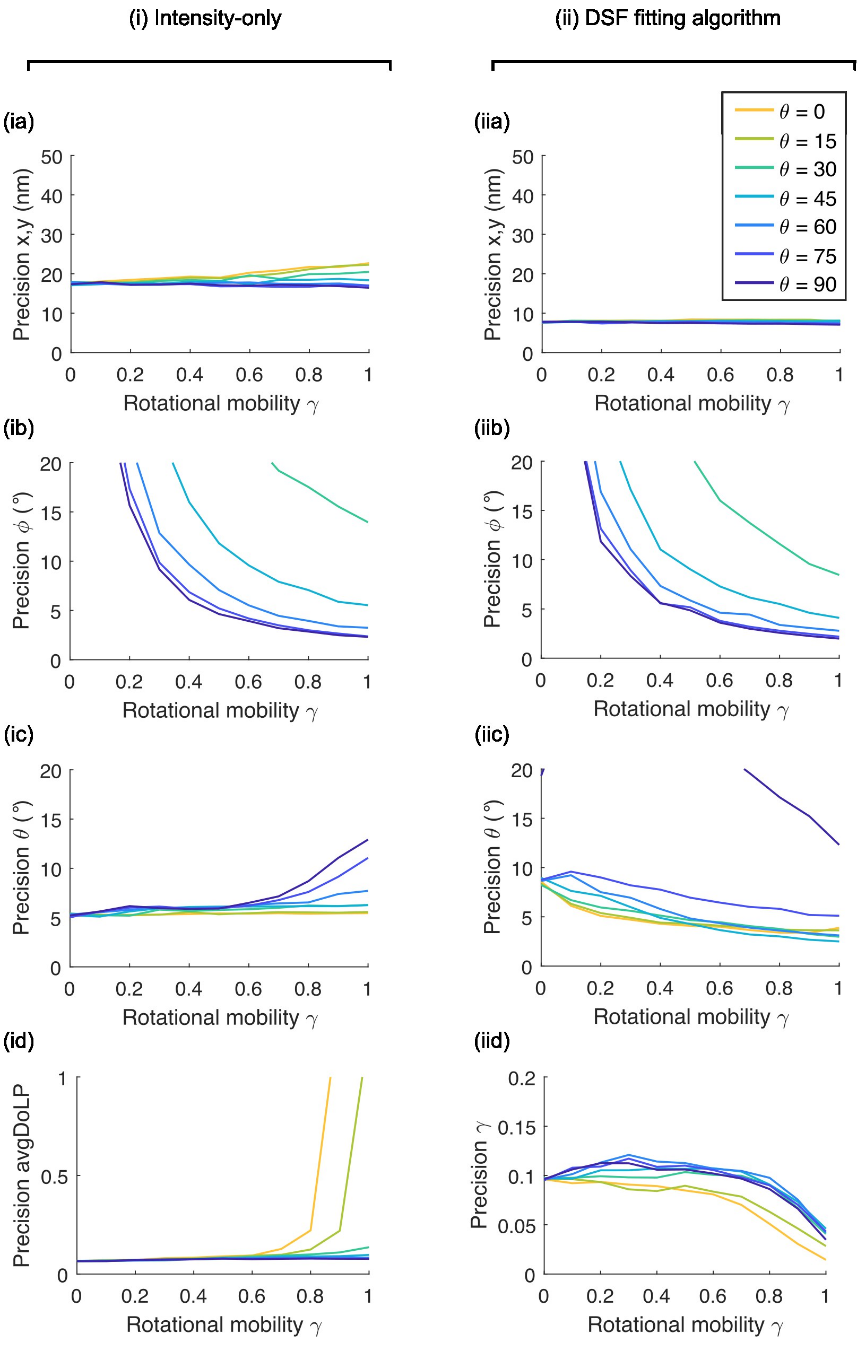

**Figure S39: Precision as a function of rotational mobility parameter $\gamma$ (10 bkgnd photons/pixels, 1000 signal photons).** The precision (standard deviation on a repeated estimate) of different parameters as a function of rotational mobility. Each plot contains multiple lines, representing a set of simulations at a fixed $\theta$. The plots are organized in three columns, each representing the results for a different algorithm: (i) intensity-only algorithm using centroid localization, (ii) intensity-only algorithm using least squares fitting of a 2D rotated asymmetric Gaussian and (iii) DSF-fitting algorithm.

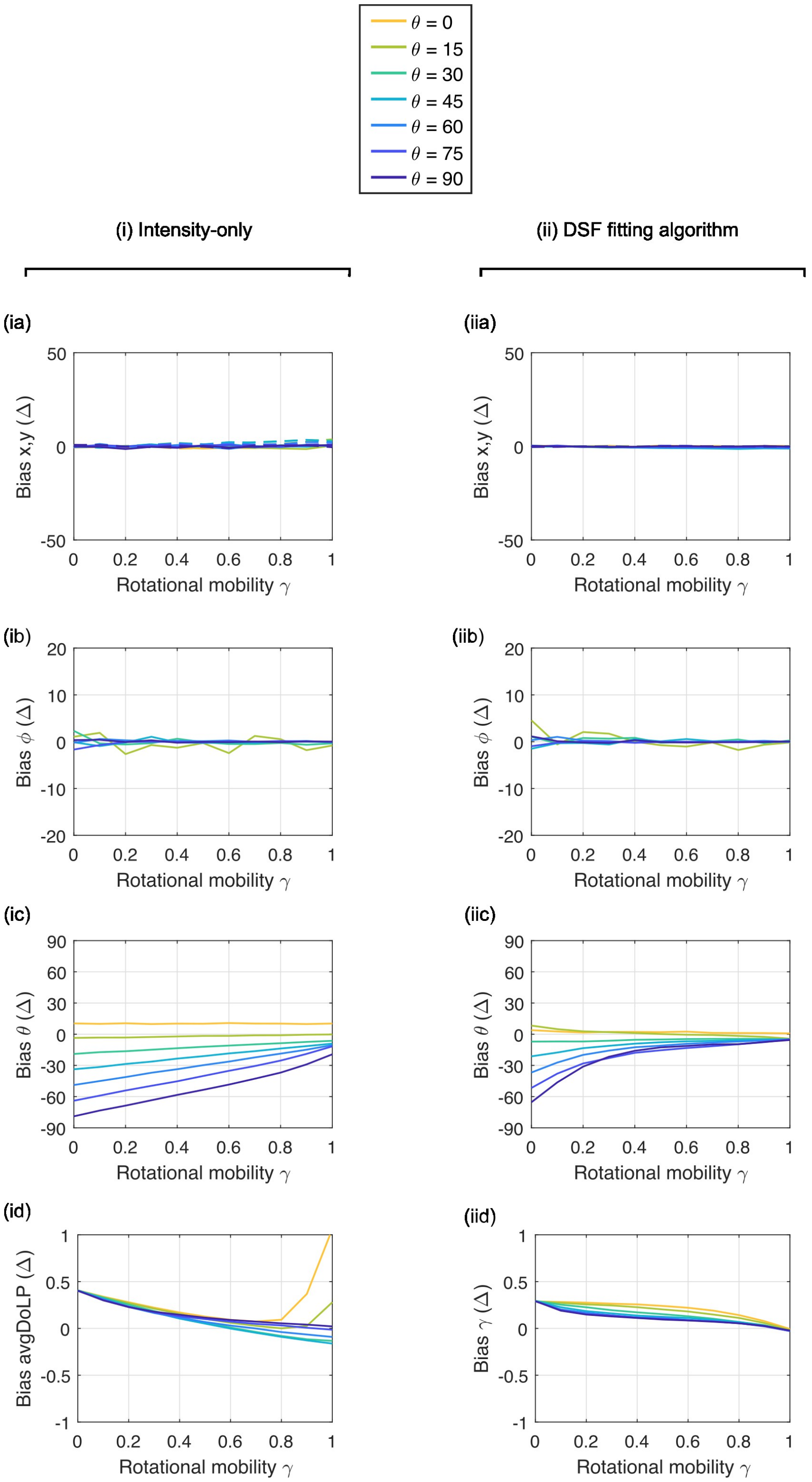

**Figure S40: Bias as a function of rotational mobility parameter $\gamma$ (10 bkgnd photons/pixels, 1000 signal photons).** The bias (difference between the true value and estimated value) of different parameters as a function of rotational mobility. Each plot contains multiple lines, representing a set of simulations at a fixed $\theta$. The plots are organized in three columns, each representing the results for a different algorithm: (i) intensity-only algorithm using centroid localization, (ii) intensity-only algorithm using least squares fitting of a 2D rotated asymmetric Gaussian and (iii) DSF-fitting algorithm.

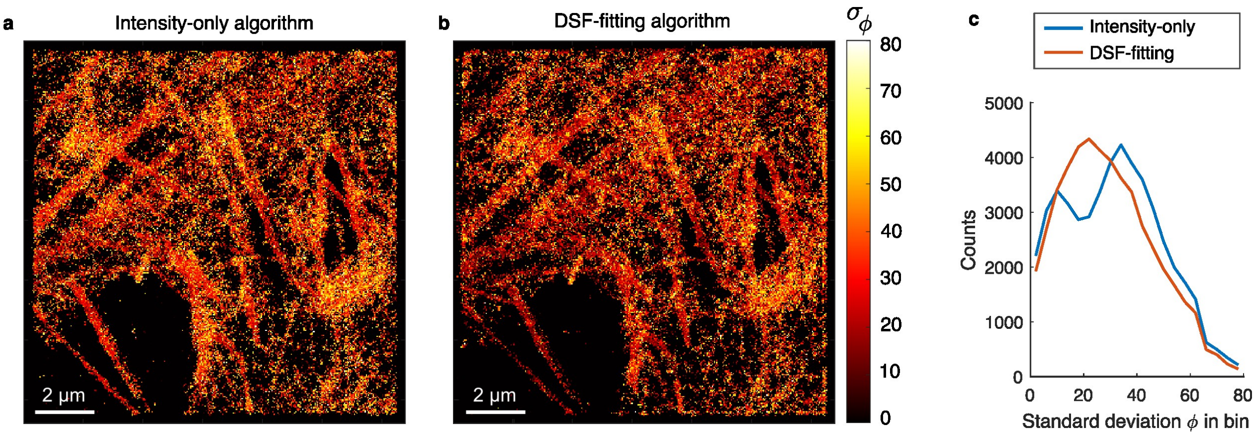

**Figure S41: Comparison of SMOLM reconstruction from different algorithms. a)** A $\phi$ standard deviation image of the dSTORM actin region from figure 5 in the main manuscript. The value of each pixel of the image represents the standard deviation on the in-plane angle of all localizations that fall within the pixel. The pixels represent 40 nm by 40 nm bins. If fewer than 5 emitters fall inside a bin, no standard deviation is calculated and the pixel gets a value of zero. **b)** The same as in panel a, but for the results from the DSF-fitting algorithm. **c)** Histograms of all non-zero values in the images in panels a and b.

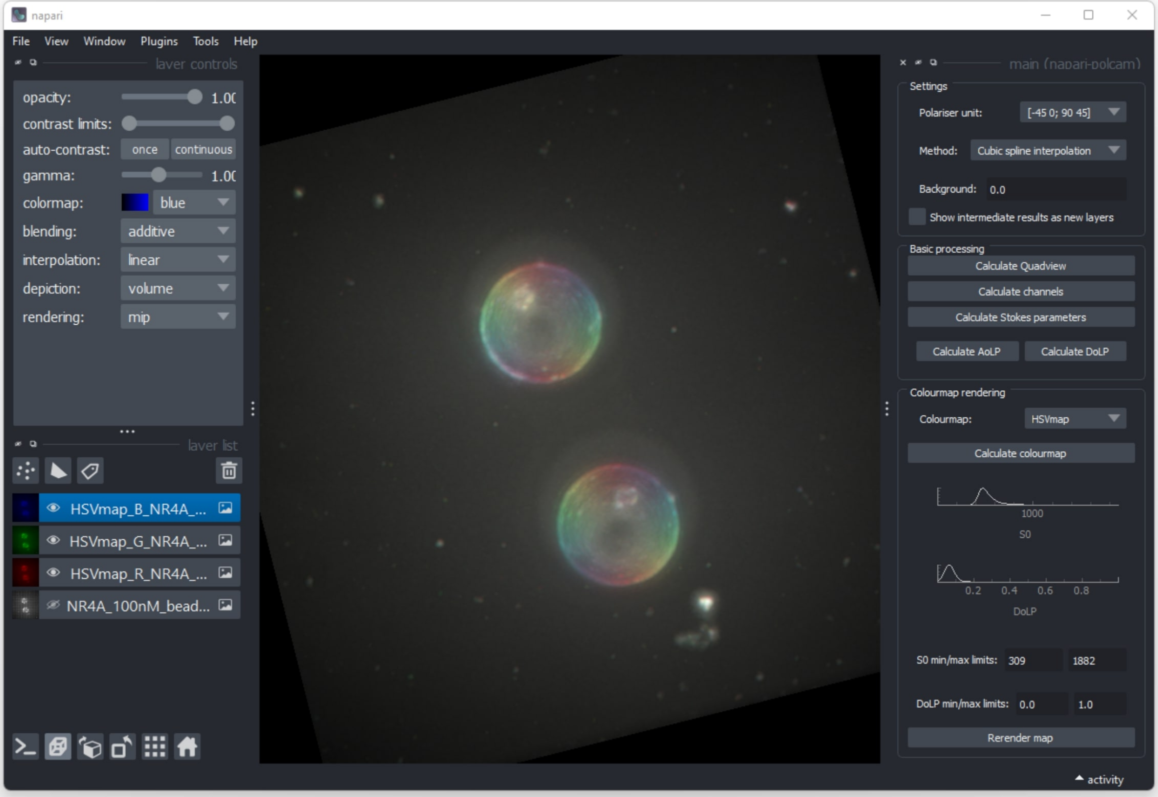

**Figure S42: Screenshot of the gui of the napari plugin napari-polcam.** The open image is an HSV polarization colormap rendering of a z-stack (maximum intensity projection view from the top) of two 5 µm diameter silica beads coated with a lipid bilayer (DPPC + 40% cholesterol) that was stained with the Nile red derivative NR4A. An unprocessed .tif stack can be loaded as normal in napari as a new layer, and when selected, the layer can be converted to different formats in napari-polcam (*e.g.* QuadView, interpolated polarised channels, Stokes parameter images, or an HSV polarization or DoLP colormap) which will appear as new layers that can be used for further image processing.

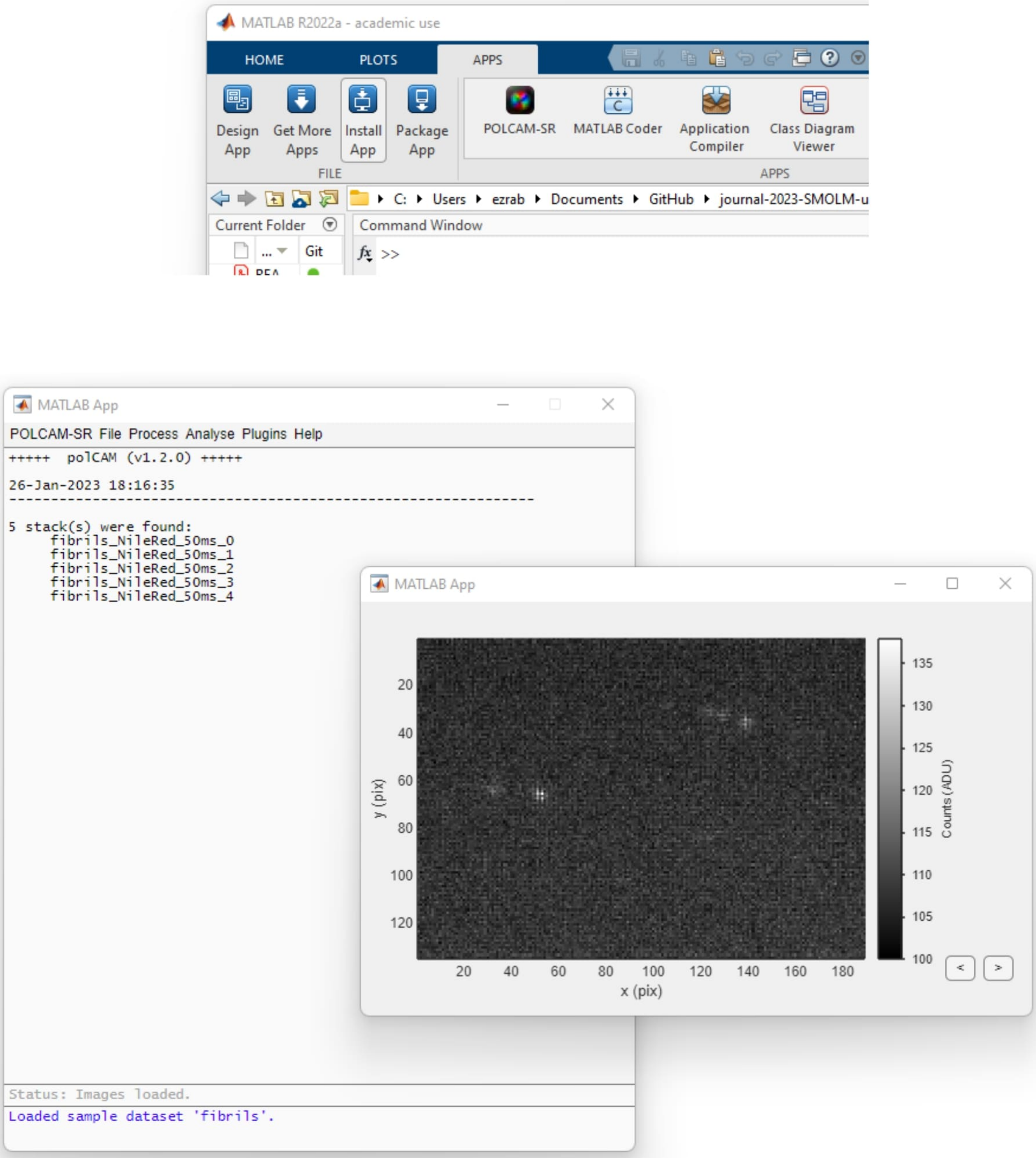

**Figure S43: Screenshot of the gui of the MATLAB plugin POLCAM-SR.** Top, after installation, the POLCAM-SR app will appear in the Apps ribbon in MATLAB. Bottom, the main gui of the POLCAM-SR app. When opening a new image dataset, an image viewing window appears. Shown here is a small test dataset (TAB-PAINT using Nile red of an alpha-synuclein fibril) which is included with the app and can be opened using *File ¿ Open samples... ¿ Fibrils* or 'ctrl + shift + F'.

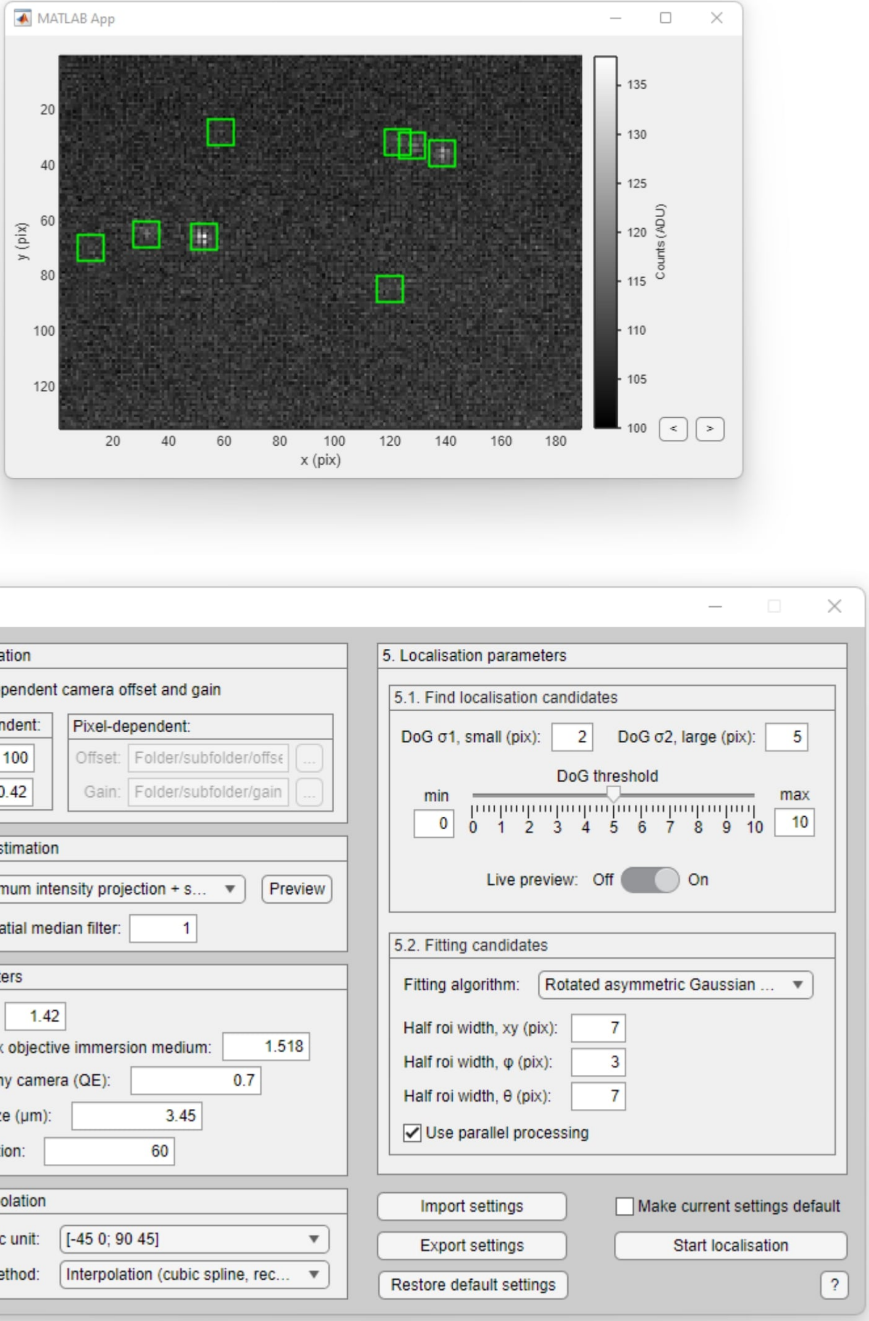

**Figure S44: Screenshot of the Localization gui of POLCAM-SR.** When an image dataset is loaded, localization can be performed by opening the Localization window using *Process ¿ Single-molecule... ¿ Localize*, or 'ctrl + L'. The localization window has different sections on 1) camera calibration parameters, 2) background estimation, 3) setup parameters, 4) the channel interpolation approach, and 5) localization. A live preview can be turned on that overlays on the image data the localization candidates (green boxes) that the software finds using the current parameter choice.

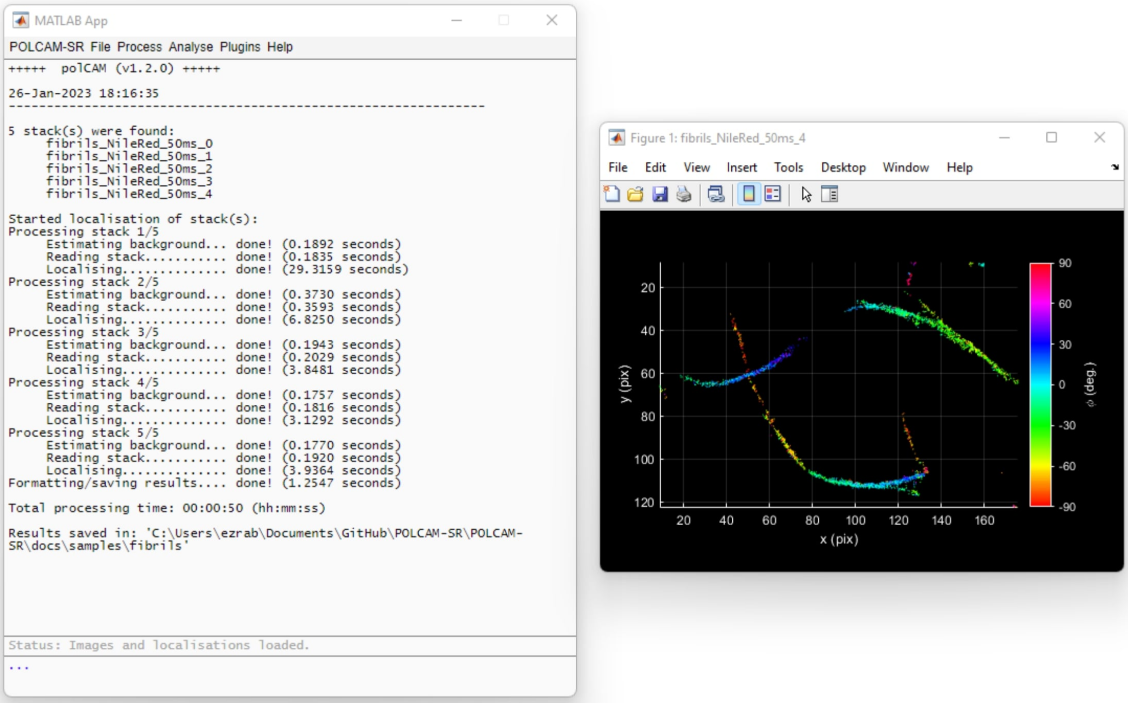

**Figure S45: Screenshot of the gui of POLCAM-SR after localization is finished.** During localization, the progress and timing are printed to the main gui of the POLCAM-SR app. Once done, a $\phi$-colourcoded scatter plot of the localizations will appear. The localization file can be filtered, drift-corrected and rendered in a range of different formats inside the app.

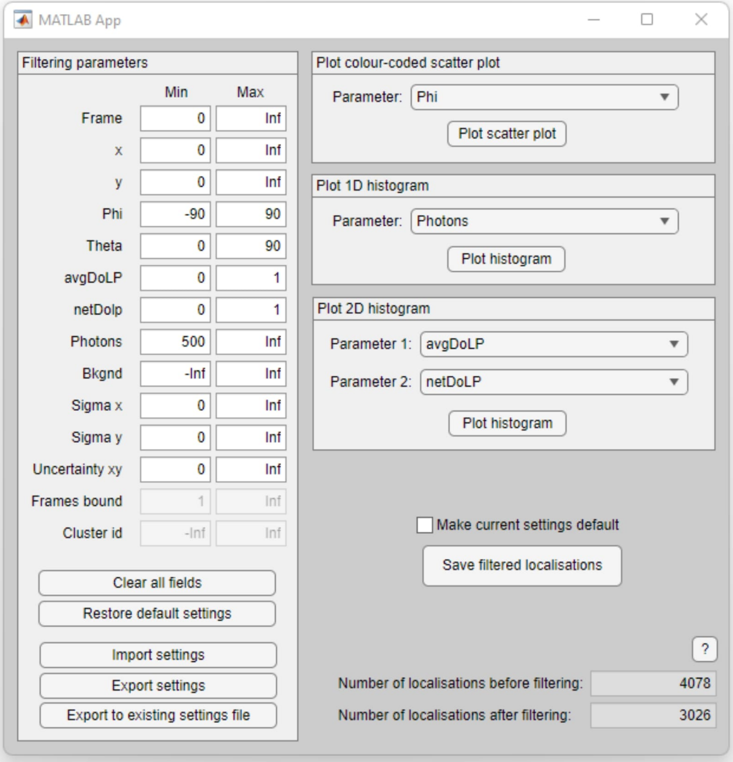

**Figure S46: Screenshot of the Filtering gui in POLCAM-SR.** When localization are loaded (because localization was just performed, or a localization file was opened), the Filtering window can be used using *Process ¿ Single-molecule... ¿ Filter*, or 'ctrl + F'. The filtering window allows setting minimum and maximum thresholds on all parameters that are generated by the localization algorithm that was used. To help with picking thresholds, color-coded scatter plots, 1D and 2D histograms using different variables from the localization file can be generated.

#### References

1. Backer, A. & Moerner, W. Extending single-molecule microscopy using optical fourier processing. *J Phys Chem B* **118,** 8313–8329 (2014).
2. Novotny, L. & Hecht, B. *Principles of Nano-Optics* (Cambridge University Press, New York, 2007).
3. Kim, D., Zhang, Z. & Xu, K. Spectrally resolved super-resolution microscopy unveils multipath reaction pathways of single spiropyran molecules. *J. Am. Chem. Soc.* **139,** 9447–9450 (2017).
4. Hecht, E. *Optics* 4th (Addison Wesley, 2002).
5. Backer, A. S. & Moerner, W. E. Determining the rotational mobility of a single molecule from a single image: a numerical study. *Optics Express* **23(4),** 4255–4276 (2015).
6. Huang, F. *et al.* Video-rate nanoscopy using sCMOS camera–specific single-molecule localization algorithms. en. *Nature Methods* **10,** 653–658. ISSN: 1548-7105. (2022) (July 2013).
7. Rueden, C., Schindelin, J. & Hiner, M. e. a. ImageJ2: ImageJ for the next generation of scientific image data. *BMC Bioinformatics* **18:529** (2017).
8. Herbert, A. *GDSC Single Molecule Light Microscopy (SMLM) ImageJ Plugins* original-date: 2014-06-19T13:52:37Z. Dec. 2022. https://github.com/aherbert/gdsc-smlm (2022).
9. Schaefer, B., Collett, E., Smyth, R., Barrett, D. & Fraher, B. Measuring the Stokes polarization parameters. *Am. J. Phys.* **75(2),** 163–168 (2007).
10. Hauge, P. Survey of methods for the complete determination of a state of polarization. *SPIE* **88,** 3–10 (1976).
11. Collett, E. Measurement of the four Stokes polarization parameters with a single circular polarizer. *Optics Communications* **52(2),** 77–80 (1984).
12. Fourkas, J. T. Rapid determination of the three-dimensional orientation of single molecules. *Opt. Lett.* **26,** 211–213. http://opg.optica.org/ol/abstract.cfm?URI=ol-26-4-211 (2001).
13. Tyo, J., LaCasse, C. & Ratliff, B. Total elimination of sampling errors in polarization imagery obtained with integrated microgrid polarimeters. *Optics Letters* **34(20)** (2009).
14. Thompson, R., Larson, D. & Webb, W. Precise nanometer localization analysis for individual fluorescent probes. *Biophysical Journal* **82(5),** 2775–2783 (2002).
15. Wang, Y. *et al.* Localization events-based sample drift correction for localization microscopy with redundant cross-correlation algorithm. *Opt Express* **22(13),** 15982–15991 (2014).
16. Schnitzbauer, J., Strauss, M., Schlichthaerle, T., Schueder, F. & Jungmann, R. Super-resolution microscopy with DNA-PAINT. *Nature Protocols* **12(6),** 1198–1228 (2017).

### References

1. Backer, A. & Moerner, W. Extending single-molecule microscopy using optical fourier processing. *J Phys Chem B* **118,** 8313–8329 (2014).
2. Novotny, L. & Hecht, B. *Principles of Nano-Optics* (Cambridge University Press, New York, 2007).
3. Kim, D., Zhang, Z. & Xu, K. Spectrally resolved super-resolution microscopy unveils multipath reaction pathways of single spiropyran molecules. *J. Am. Chem. Soc.* **139,** 9447–9450 (2017).
4. Hecht, E. *Optics* 4th (Addison Wesley, 2002).
5. Backer, A. S. & Moerner, W. E. Determining the rotational mobility of a single molecule from a single image: a numerical study. *Optics Express* **23(4),** 4255–4276 (2015).
6. Huang, F. *et al.* Video-rate nanoscopy using sCMOS camera–specific single-molecule localization algorithms. en. *Nature Methods* **10,** 653–658. ISSN: 1548-7105. (2022) (July 2013).
7. Rueden, C., Schindelin, J. & Hiner, M. e. a. ImageJ2: ImageJ for the next generation of scientific image data. *BMC Bioinformatics* **18:529** (2017).
8. Herbert, A. *GDSC Single Molecule Light Microscopy (SMLM) ImageJ Plugins* original-date: 2014-06-19T13:52:37Z. Dec. 2022. https://github.com/aherbert/gdsc-smlm (2022).
9. Schaefer, B., Collett, E., Smyth, R., Barrett, D. & Fraher, B. Measuring the Stokes polarization parameters. *Am. J. Phys.* **75(2),** 163–168 (2007).
10. Hauge, P. Survey of methods for the complete determination of a state of polarization. *SPIE* **88,** 3–10 (1976).
11. Collett, E. Measurement of the four Stokes polarization parameters with a single circular polarizer. *Optics Communications* **52(2),** 77–80 (1984).
12. Fourkas, J. T. Rapid determination of the three-dimensional orientation of single molecules. *Opt. Lett.* **26,** 211–213. http://opg.optica.org/ol/abstract.cfm?URI=ol-26-4-211 (2001).
13. Tyo, J., LaCasse, C. & Ratliff, B. Total elimination of sampling errors in polarization imagery obtained with integrated microgrid polarimeters. *Optics Letters* **34(20)** (2009).
14. Thompson, R., Larson, D. & Webb, W. Precise nanometer localization analysis for individual fluorescent probes. *Biophysical Journal* **82(5),** 2775–2783 (2002).
15. Wang, Y. *et al.* Localization events-based sample drift correction for localization microscopy with redundant cross-correlation algorithm. *Opt Express* **22(13),** 15982–15991 (2014).
16. Schnitzbauer, J., Strauss, M., Schlichthaerle, T., Schueder, F. & Jungmann, R. Super-resolution microscopy with DNA-PAINT. *Nature Protocols* **12(6),** 1198–1228 (2017).
